# Expanding the Triangle of U: The genome assembly of *Hirschfeldia incana* provides insights into chromosomal evolution, phylogenomics and high photosynthesis-related traits

**DOI:** 10.1101/2024.05.16.593662

**Authors:** Nam V. Hoang, Nora Walden, Ludovico Caracciolo, Sofia Bengoa Luoni, Moges Retta, Run Li, Felicia C. Wolters, Tina Woldu, Frank F. M. Becker, Patrick Verbaarschot, Jeremy Harbinson, Steven M. Driever, Paul C. Struik, Herbert van Amerongen, Dick de Ridder, Mark G.M. Aarts, M. Eric Schranz

**Affiliations:** Biosystematics Group, Wageningen University and Research, Droevendaalsesteeg 1, 6708 PB Wageningen, The Netherlands; Centre for Organismal Studies, Heidelberg University, Heidelberg, Germany; Laboratory of Biophysics, Wageningen University and Research, Stippeneng 4, 6708 WE Wageningen, The Netherlands; Laboratory of Genetics, Wageningen University and Research, Droevendaalsesteeg 1, 6708 PB Wageningen, The Netherlands; Centre for Crop Systems Analysis, Wageningen University and Research, P.O. Box 430, 6700 AK Wageningen, The Netherlands; Bioinformatics Group, Wageningen University and Research, Droevendaalsesteeg 1, 6708 PB Wageningen, The Netherlands

**Keywords:** *Hirschfeldia incana*, Brassicaceae, Brassica, whole-genome duplication, photosynthesis evolution, polyploidy, gene fractionation, sub-genome dominance

## Abstract

The Brassiceae tribe encompasses many economically important crops and exhibits high intraspecific and interspecific phenotypic variation. After a shared whole-genome triplication (WGT) event (*Br-α*, ∼15.9 million years ago), different lineages and species underwent differential chromosomal rearrangements (diploidization) leading to diverse patterns of gene retention and loss (fractionation). Lineage diversification and genomic changes contributed to an array of divergence in morphology, biochemistry, and physiology underlying photosynthesis-related traits. The C_3_ species *Hirschfeldia incana* is studied as it displays high photosynthetic rates under high-light conditions. We present an improved chromosome-level genome assembly for *H. incana* (Nijmegen, v2.0) using nanopore and chromosome conformation capture (Hi-C) technologies, with 409Mb in size and an N50 of 52Mb (a 10× improvement over the previously published scaffold-level v1.0 assembly). The updated assembly and annotation allowed to investigate the WGT history of *H. incana* in a comparative phylogenomic framework from the Brassiceae ancestral genomic blocks and related diploidized crops. *Hirschfeldia incana* (x=7) shares extensive genome collinearity with *Raphanus sativus* (x=9). These two species share some commonalities with *Brassica rapa* and *B. oleracea* (A genome, x=10 and C genome, x=9, respectively) and other similarities with *B. nigra* (B genome, x=8). Phylogenetic analysis revealed that *H. incana* and *R. sativus* form a monophyletic clade in between the *Brassica* A/C and B genomes. We postulate that *H. incana* and *R. sativus* genomes are results of reciprocal hybridization combinations of the *Brassica* A/C and B genome types. Our results might explain the discrepancy observed in published studies regarding phylogenetic placement of *H. incana* and *R. sativus* in relation to the “Triangle of U” species. Expression analysis of WGT retained gene copies revealed sub-genome expression divergence, likely due to neo- or sub-functionalization. Finally, we highlighted genes associated with physio-biochemical-anatomical adaptive changes observed in *H. incana* which likely facilitate its high-photosynthesis traits under high light.

## INTRODUCTION

The Brassicaceae family contains the model plant *Arabidopsis thaliana* and many economically important vegetable, root and oil crops. Many of these crops are members of the Brassiceae tribe (*Brassicas*) that underwent a meso-hexaploidy *Brassica α* whole-genome triplication event (*Br-α* WGT), which occurred ∼15.9 million years ago (MYA) in the middle of the Miocene epoch (Jiao et al., 2011; Liu et al., 2014). This has resulted in a massive inter- and intraspecific phenotypic variation due to differential chromosomal rearrangements (i.e., diploidization) and gene retention and loss (i.e., fractionation) (Cheng et al., 2014; Cheng et al., 2016). While no true C_4_ photosynthesis in Brassiceae species has been reported so far, the tribe consists of species that exhibit a wide range of light-saturated photosynthetic rates, and utilize both the C_3_ and C_3_-C_4_ photosynthetic pathways (Schluter et al., 2023). Among these, the C_3_ species *Hirschfeldia incana* (grey mustard, n=x=7) was reported to display high photosynthetic rates (i.e., carbon assimilation) under high-light conditions (Canvin et al., 1980; Garassino et al., 2022). This, together with its close evolutionary proximity to the model plant *Arabidopsis* and to *Brassica* crops, positions *H. incana* as a good model to study how high-photosynthesis traits have evolved in the Brassiceae tribe.

In the last two decades, significant genomic resources were developed for *Arabidopsis* and *Brassica* crops, including *B. rapa*, *B. oleracea*, *B. nigra* and their allo-tetraploid hybrids which are part of the “Triangle of U” (Wu et al., 2022). However, genomic data available for *H. incana* and its wild relatives in the Brassiceae tribe is still limited, which hinders our understanding of the genetic basis of its high photosynthesis under high irradiance conditions. More recently, high-quality chromosome-level genome sequences of the close relatives *Raphanus sativus* and *Sinapis arvensis* were released (Cho et al., 2022; Xu et al., 2023; Yang et al., 2023). Notably, a new genomic panel was developed for a total of 18 Brassiceae species at scaffold-level with different photosynthesis types, including C_3_ and C_3_-C_4_ (Guerreiro et al., 2023). These resources are expected to support large-scale comparative phylogenomic studies in combination with high-throughput phenotyping, focusing on the *Nigra - Rapa/Oleracea - Raphanus* clades to untangle the evolution of the high-photosynthesis traits in the Brassiceae tribe. These are also critical for resolving the nuclear genome-based phylogenetic relationships and designation of Brassiceae species, which still remain controversial (Huang et al., 2016; Hendriks et al., 2023). To date, there are at least two scaffold-level genome assemblies of different *H. incana* accessions (Nijmegen, NIJ, and HIR1) (Garassino et al., 2022; Guerreiro et al., 2023) and a few transcriptome datasets (Mabry et al., 2020; Garassino et al., 2022; Hasnaoui et al., 2022; Garassino et al., 2023), which have facilitated studies on comparative genomics and the expression of photosynthesis-related genes. However, for genome evolution and synteny-based studies, it is imperative to have chromosome-level assemblies to uncover the evolutionary trajectory that led to high-photosynthesis traits in this species. Genome synteny, i.e. conserved gene order across different genomes (Liu et al., 2018), is therefore crucial for evolution studies, since it allows the detection of chromosome-level reorganisation events, including whole-genome duplication/triplication (WGD/WGT), that gave rise to different taxonomical clades.

It has been proposed that the Brassiceae genomes underwent a “two-step” hybridization (Cheng et al., 2012; Cheng et al., 2014; Hao et al., 2021) that resulted in three distinct sub-genomes of different origins, namely the least-fractionated (LF), medium-fractionated (MF_1_) and most-fractionated (MF_2_) sub-genomes. During this process, the MF_1_ and MF_2_ sub-genomes initially emerged together through an auto-tetraploidization event, followed by a first round of diploidization and fractionation. Then, the LF sub-genome was subsequently added to form an allo-hexaploid genome, which was also followed by another round of diploidization and fractionation. Pioneering work suggested that these sub-genomes were derived from the common ancestral tPCK (translocated Proto-Calepineae Karyotype) of all Brassiceae species (Schranz et al., 2006; Lysak et al., 2016). Typically, the three sub-genomes display differential gene fractionation rates and a gene expression bias, with the LF sub-genome being dominant. From a broader perspective, this sub-genome dominance phenomenon has been observed in several families including Brassicaceae (Cheng et al., 2012; Liu et al., 2014; Perumal et al., 2020; Yang et al., 2023), Cleomaceae (Hoang et al., 2023), Poaceae (International Wheat Genome Sequencing, 2014) and Asteraceae (Barker et al., 2016).

The tribe Brassiceae, and likewise the Brassicaceae family, is notorious for having phylogenetic trees with poorly supported bootstrap values due to their complicated history of allo-polyploidization, incomplete lineage sorting, and rampant introgressive hybridization (Hendriks et al., 2023). It is also expected that nuclear genes were selected for cyto-nuclear compatibility in the hybrid genotypes, which in turn led to cytonuclear phylogenetic discordance (Forsythe et al., 2020). This could mislead our understanding of species relationships and evolution. Regarding the nuclear phylogenetic placement of *H. incana* in relation to the species in the “Triangle of U”, published studies appear to be inconsistent. *Hirschfeldia incana* was either grouped closely to *B. rapa*/*B. oleracea* (A/C genome type) in Huang et al. (2016) or with *B. nigra* (B genome type) in Garassino et al. (2022). Interestingly, this is similar to the case of radish (*R. sativus*) (Cho et al., 2022; Yang et al., 2023). It was previously revealed that the *R. sativus* genome exhibits intermediate characteristics between the *Brassica* A/C and B genome types (Jeong et al., 2016; Cho et al., 2022), which might explain the observed phylogenetic incongruency among nuclear trees in published studies. For example, different contributions of the progenitor genomes to a set of genetic markers that are used for phylogenetic studies could possibly change the species’ phylogenetic placement. With a high-quality genome, it would be possible to investigate the placement of *H. incana* and its relationships with the Brassiceae species through chromosome and genome evolution.

Due to its importance, photosynthesis has become one of the most studied processes in plant science with the aim to increase agricultural production. While the model C_3_ plant *A. thaliana* was used in most fundamental research and discoveries, several other model systems of different photosynthesis types have also been established, including that of the C_4_ (Brown et al., 2005), C_3_-C_4_ (Gowik et al., 2011) and CAM (Edwards, 2019) types. These studies have resulted in the identification of targets for photosynthesis improvement through manipulation of biochemical metabolic pathways, canopy architecture and leaf anatomy, as well as the underlying mechanism of natural photosynthesis variation (Lawson et al., 2012; Tholen et al., 2012; Zhu et al., 2013; Theeuwen et al., 2022). Key genetic factors responsible for these target features would be pivotal to allow redesigning crops with desirable high-photosynthesis traits. Regarding *H. incana*’s ability to maintain a high photosynthetic efficiency in high-light conditions, recent advances in high-throughput phenotyping and sequencing technologies could facilitate the investigation of its genetic basis and thereby suggest potential targets for photosynthesis improvement. This will also potentially explain the evolution that gave rise to its high-photosynthesis traits.

In this study, we present an improved chromosome-level assembly (v2.0) of the *H. incana* Nijmegen accession based on a combination of Oxford Nanopore Technology (ONT) sequencing and chromosome conformation capture (Hi-C) data. We also provide an updated genome annotation that includes more gene models than the previous version and is comparable in gene model number to those of Brassiceae genomes that underwent the *Br-α* WGT event. The improved *H. incana* genome assembly and annotation allowed to elucidate the genome evolution of this species in relation to the *Brassica* species and other species within the Brassiceae tribe. We showed that, like the *Brassica* genomes, the *H. incana* genome was also derived from the common ancestral tPCK genomic blocks. The triplicated ancestral genomic blocks within the *H. incana* genome could be classified into three sub-genomes, with the LF sub-genome showing dominance in gene retention as well as gene expression. The *H. incana* genome appears to be similar to that of *R. sativus* in terms of collinearity, and displays intermediate characteristics of *Brassica* A/C and B genome types. The results might explain the discrepancy observed in the published studies regarding the phylogenetic placement of *H. incana* and *R. sativus* in relation to species within the “Triangle of U”. Finally, we highlight genes that are associated with the physio-biochemical-anatomical adaptive changes observed in *H. incana* which likely facilitate its high rate of carbon assimilation when grown under high-light intensity. The updated assembly and annotation will be a valuable resource for future research to explore the genetic basis of this interesting species in terms of retaining a high light-use efficiency at high-light intensity, for example, through exploring the natural genetic variations within the *H. incana* accessions or interspecific comparative genomics/transcriptomics to pinpoint the underlying mechanism responsible for its variation in photosynthetic efficiency.

## RESULTS AND DISCUSSION

### A chromosome-level assembly and annotation of the *Hirschfeldia incana* genome

The scaffold-level genome assembly v1.0 of the Nijmegen accession of C_3_ species *H. incana* was previously reconstructed based on PacBio SMRT long-read, 10X Genomics linked-read, and Illumina short-read data (Garassino et al., 2022). Here, we first improved the draft genome assembly v1.0 through two rounds of scaffolding using ONT and Hi-C sequencing, respectively. We generated a total of about 20 Gb long-read ONT data (N50 = 26 kb, ∼48x genome coverage) and 124 million Hi-C Illumina reads (150 bp, ∼44x genome coverage) (**Supplemental Table S1** and **Fig. S1**). The ONT data were used to link v1.0 scaffolds into larger contigs to obtain an ONT-derived assembly (termed as v1.5). We subsequently used v1.5 as input for a second round of scaffolding based on the Hi-C data to produce the final chromosome-level genome assembly (termed as v2.0). Compared to the previous assembly v1.0 (size 398 Mb, N50 of 5 Mb), both v1.5 and v2.0 have a slightly larger assembly size of 409 Mb, and significantly improved N50 lengths, of 14 Mb and 52 Mb, respectively (**Table 1**). Overall, the size of the three assemblies is close to our re-estimated genome size of 421 Mb using *k*-mer analysis (**Supplemental Fig. S2** and **Methods**) and smaller than the flow cytometry estimate of 487 Mb (Garassino et al., 2022). Our v2.0 assembly N50 length (52 Mb) and chromosome size are comparable to other chromosome-level assemblies of related Brassiceae species including *B. rapa* v4.0 and v4.1 (Zhang et al., 2023), *B. oleracea* JZS v2.0 (Cai et al., 2020) and *B. nigra* N100 v2.0 (Perumal et al., 2020), with N50 lengths ranging from 43 to 61 Mb (**Supplemental Table S2**) and individual chromosome sizes of 30 to 75 Mb (**Supplemental Fig. S3**).

**Table 1.** Summary statistics of the genome assembly and annotation of *H. incana* and relatives.

| Genome features | *B. rapa* v4.0 (Chiifu) | *B. oleracea* v2.0 (JZS) | *B. nigra* v2.0 (NI100) | *H. incana* v1.0 (NIJ) | *H. incana* v1.5 (NIJ) | *H. incana* v2.0 (NIJ) |
| --- | --- | --- | --- | --- | --- | --- |
| Chromosome number (n=x) | 10 | 9 | 8 | 7 | 7 | 7 |
| Assembled genome size (Mb) | 424.59 | 561.16 | 506.00 | 398.50 | 408.86 | 408.93 |
| GC content (assembly, %) | 37.59 | 36.75 | 38.21 | 36.18 | 36.20 | 36.20 |
| Number of scaffolds | 10 | 649 | 58 | 384 | 246 | 358 |
| Number of pseudomolecules | 10 | 9 | 8 | N/A | N/A | 7 |
| N50 length (Mb) | 43.05 | 57.88 | 60.82 | 5.11 | 13.76 | 52.38 |
| N90 length (Mb) | 30.43 | 47.84 | 55.08 | 0.83 | 1.99 | 48.08 |
| Longest scaffold (Mb) | 73.37 | 74.51 | 70.85 | 15.00 | 30.16 | 63.71 |
| N's per 100 kb | 0.05 | 26.05 | 2.47 | 13.45 | 1.70 | 18.08 |
| BUSCO assembly (%) | 99.5 | 99.5 | 99.2 | 99.3 | 99.5 | 99.2 |
| Number of genes | 47,531 | 59,064 | 59,852 | 32,312 | 54,459 | 54,457 |
| GC content (main transcript, %) | 46.24 | 46.13 | 45.92 | 46.53 | 46.19 | 46.19 |
| N50 length (main transcript, bp) | 1,473 | 1,428 | 1,449 | 1,515 | 1,377 | 1,377 |
| Mean length (main transcript, bp) | 1,147 | 1,155 | 1,026 | 1,247 | 1,035 | 1,035 |
| BUSCO (main transcripts, %) | 97.2 | 98.7 | 98.2 | 96.2 | 97.8 | 97.7 |

The final genome assembly v2.0 has 358 scaffolds with the majority (91.2% assembly length and 96.0% predicted genes) anchored onto seven super scaffolds (**Fig. 1A** and **B**, **Table 1**) which correspond to the seven (x=7) reported chromosomes for *H. incana* (Garassino et al., 2022). All seven chromosomes contain centromere-specific repeat sequences, CENTs (Jeong et al., 2016; Cho et al., 2022), detected within the repeat-rich regions (**Fig. 1B**). By mapping back the 102 million whole-genome sequencing (WGS) Illumina reads generated by Garassino et al. (2022), it was found that all three assemblies have comparable mapping rates of 91.4% (v1.0) and 91.5% (v1.5 and v2.0) (**Supplemental Table S3**). The BUSCO (Benchmarking Universal Single-Copy Orthologs) completeness score (Simao et al., 2015) of all three assemblies ranged from 99.2% to 99.5% (**Supplemental Table S4**). This indicates that we successfully incorporated v1.0 sequences into our final chromosome-level assembly v2.0, and all three assemblies represent the gene content of the *H. incana* genome well.

**Figure 1.**
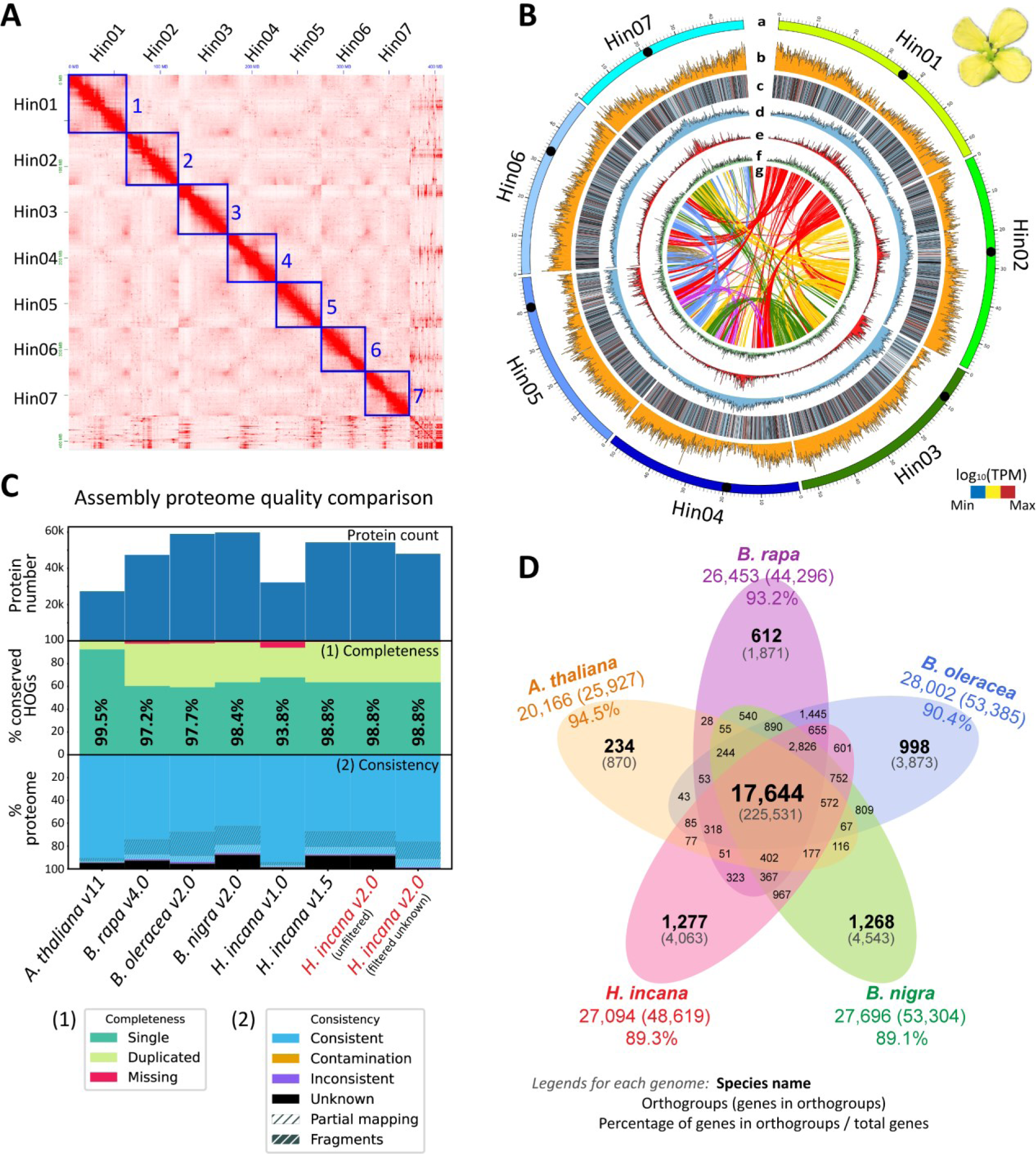
The assembly of *H. incana* Nijmegen genome v2.0, intra-genomic synteny, quality assessment and gene orthogroup clustering. **(A)** Chromosome-level Hi-C contact map of the *H. incana* genome, highlighting 7 blocks that correspond to 7 chromosomes. **(B)** Circos plot showing 7 chromosomes of the *H. incana* genome (track a), gene density (track b), gene expression of the predicted gene models (track c), distribution of all repetitive elements (track d), distribution of LTR elements (track e), DNA transposon elements (track f) and intra-genomic synteny (*minspan = 4 genes*) (track g). Distributions were estimated for each window of 100 kb. Gene expression was calculated using the transcriptome data from Garassino et al. (2023) as log_10_(average transcripts per million transcripts, TPM) over all 10 samples from two conditions, low-light and high-light. Ribbon links in the inner track of the Circos plot represent intra-genomic syntenic regions among chromosomes. For a dotplot, see **Supplemental Fig. S9**. Centromere regions are indicated by black dots on each chromosome. Length is in Mb. **(C)** Comparisons of proteomes among selected Brassicaceae genomes by OMArk tool, including proteome count, completeness, and annotation consistency against a total of 17,999 conserved orthologs of the Brassicaceae family in the OMA database. Percentage of completeness for each genome is the total percentage of single and duplicated completeness of the hierarchical orthologous groups (HOGs). For *H. incana* v2.0, two proteome datasets, unfiltered and filtered, are shown. The filtered dataset was obtained from the initial proteome after the removal of unknown proteins that were not found in the OMA database. **(D)** Venn diagram showing shared and unique orthogroups in *A. thaliana*, *B. rapa, B. oleracea, B. nigra* and *H. incana*. Numbers in brackets denote genes included in the orthogroups (as explained in the figure). Percentages were calculated based on the total genes annotated in each selected genome. For panels **C** and **D**, the genomes of *A. thaliana* Col-0, *B. rapa* Chiifu, *B. oleracea* JZS, *B. nigra* NI100 and *H. incana* NIJ were used (see **Methods** for more details).

As expected, the repeat content of the *H. incana* genome assembly v2.0 is very similar to that of v1.0, with 50.3% of the sequences being masked as repetitive elements, present in two major classes, the long terminal repeat retrotransposons (LTR-RT) and DNA transposons (**Fig. 1B** and **Supplemental Table S5**). By integrating various gene prediction approaches, we annotated a total of 54,457 protein-coding gene models (59,417 total transcripts) and 1,262 tRNAs (**Supplemental Figs. S4** and **S5**) with a 97.7% BUSCO completeness score (**Supplemental Table S6**). The v2.0 annotation is highly syntenic with that of v1.0 as indicated by the syntenic path assembly in their dotplot (**Supplemental Fig. S6**). Most of the gene models (99.9%) matched with sequences in at least one of the public protein databases or assembled transcripts (**Supplemental Table S7**), including 71.9% matching with Swiss-Prot (O’Donovan et al., 2002), 91.2% with TrEMBL (O’Donovan et al., 2002), 74.2% with InterPro (Zdobnov and Apweiler, 2001), 62% with gene ontology (GO) (Ashburner et al., 2000), 55.0% with Kyoto Encyclopedia of Genes and Genomes (KEGG) (Kanehisa and Goto, 2000), 88.4% with OMA database (Nevers et al., 2022), and 87.9% with assembled transcripts from *H. incana*.

To further assess our gene annotation, we employed the OMArk tool (Nevers et al., 2022) to compare our proteome completeness and consistency with that of four closely related species using 17,999 conserved hierarchical orthologous groups (HOGs) of the Brassicaceae family. The completeness score of the *H. incana* proteome v2.0 was 98.8% (compared to 93.8% for v1.0), and comparable to that of other Brassicaceae species proteomes (**Figure 1C** and **Supplemental Fig. S7** and **Table S8**). The percentage of our proteome that matched the conserved Brassicaceae HOGs in the OMA database is slightly higher than *B. nigra* (88.4% vs. 87.7%), but lower than *B. rapa*, *B. oleracea* and *A. thaliana* (92.8%, 95.7% and 94.7%, respectively). Additional ortholog clustering by OrthoFinder (Emms and Kelly, 2019) of our proteome and the proteomes of the aforementioned Brassicaceae genomes resulted in 48,619 *H. incana* genes (89.3% of total genes) being classified into 27,094 orthogroups, of which 17,644 were commonly shared with four other Brassicaceae proteomes (**Figure 1D** and **Supplemental Table S9**). Collectively, these assessments indicate that our improved genome assembly and annotation of *H. incana* v2.0 is of good quality and could be used together with that of its Brassiceae relatives for synteny-based and trait evolution studies of high photosynthetic rates in the Brassiceae tribe.

### The *Hirschfeldia incana* genome exhibits the typical triplicated structure of the Brassiceae tribe

The meso-hexaploidy *Br-α* WGT event was previously reported based on the genomes of several Brassiceae species including those from the genera *Brassica* (Wang et al., 2011; Liu et al., 2014; Perumal et al., 2020), *Raphanus* (Jeong et al., 2016; Xu et al., 2023) and *Sinapis* (Yang et al., 2023). As a member of the Brassiceae tribe, it is expected that the *H. incana* genome also underwent the *Br-α* WGT event, even though it is not clear if it shared the same progenitors involved in the two-step speciation process (Cheng et al., 2012) that gave rise to other hexaploid species within the Brassiceae. Here, using the updated chromosome-scale assembly and annotation and the total syntenic gene pairs between *A. thaliana* and *H. incana*, we find a clear 1:3 syntenic pattern between the two respective species. More specifically, 87% of the *H. incana* genes have one syntenic block in the *A. thaliana* genome, while 8%, 33% and 54% of the *A. thaliana* genes have 1, 2 and 3 syntenic blocks in the *H. incana* genome, respectively (**Fig. 2A** and **Supplemental Fig. S8**). Those *H. incana* genes syntenic to 3 *A. thaliana* syntenic blocks are likely located within the well retained genomic regions, while *H. incana* genes syntenic to 2 blocks or 1 block are likely those within more fractionated regions. The syntenic relationship between *H. incana* and *A. thaliana* genomes resembles the relatedness between each of the three *Brassica* genomes (*B. rapa*, *B. oleracea* and *B. nigra*) and *A. thaliana*, as illustrated in **Fig. 2B**. Additionally, a clear triplicated intra-genomic syntenic pattern could be observed within the *H. incana* genome (**Supplemental Fig. S9**), while a 3-to-3 (or 1-to-1, if only true orthologs were considered) syntenic relationship between *H. incana* and *B. rapa* was observed when comparing the two genomes (**Supplemental Fig. S10**). Our results support the notion that the *H. incana* genome, like other Brassiceae genomes, also experienced the Brassiceae tribe-specific hexaploidy *Br-α* WGT event.

**Figure 2.**
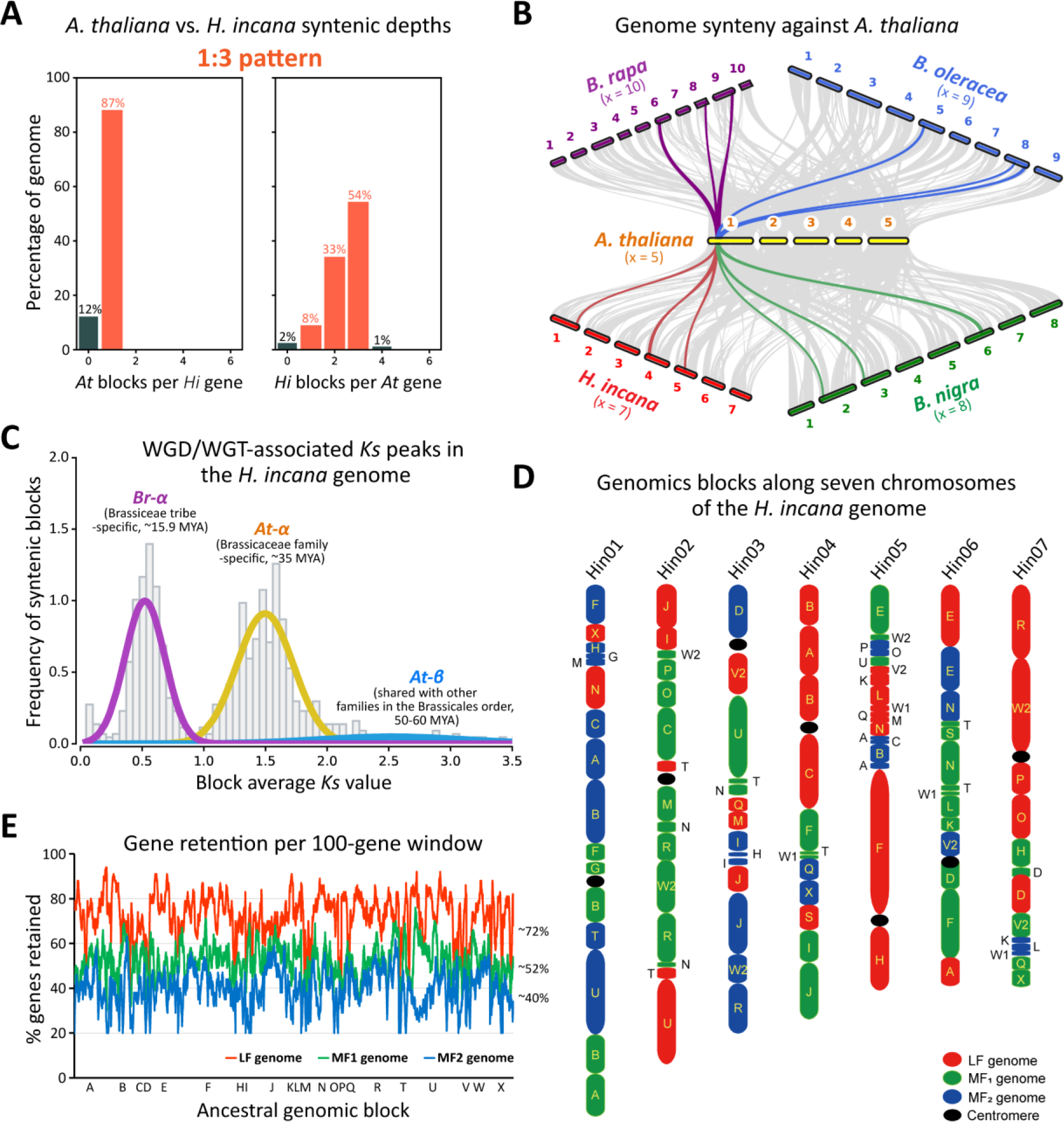
Genomic architecture of the *H. incana* Nijmegen genome. **(A)** Ratio of syntenic depth between *A. thaliana* and *H. incana*. Syntenic blocks of *A. thaliana* per *H. incana* gene (left) and syntenic blocks of *H. incana* per *A. thaliana* gene (right) are shown which suggest a clear 1:3 pattern between the two genomes. **(B)** Genome macro-synteny between *A. thaliana* and four Brassiceae genomes (*B. rapa, B. oleracea, B. nigra* and *H. incana*) that underwent the *Br-α* WGT event, showing a similar 1:3 syntenic pattern. A syntenic block located on *A. thaliana* chromosome 1 was chosen to illustrate the triplicated pattern in the Brassiceae genomes. The genomes of *A. thaliana* Col-0, *B. rapa* Chiifu, *B. oleracea* JZS, *B. nigra* NI100 and *H. incana* NIJ were used. **(C)** WGT/WGD events identified in the *H. incana* genome by fitting the *Ks* distributions for WGT/WGD-derived gene pairs using a Gaussian Mixture Model (GMM). *Ks* peaks correspond to the *At-β* (shared with other families within the Brassicales), *At-α* (Brassicaceae-specific) and *Br-α* (Brassiceae-specific) events. Only *Ks* ≤3.5 were included in this analysis. MYA denotes million years ago. Date estimates were derived from Jiao et al. (2011); Kagale et al. (2014); Liu et al. (2014); Edger et al. (2018). **(D)** Ancestral genomic blocks along the seven *H. incana* chromosomes. The updated tPCK ancestral genome from He et al. (2021) was mapped onto the *H. incana* genome to identify intervals of the triplicated conserved genomic blocks, then these triplicated blocks were classified into three sub-genomes (LF, MF_1_ and MF_2_) based on their gene retention pattern analyzed by FractBias (Joyce et al., 2017). Sub-genomes were colored as red, green, and blue, respectively. Centromeres are represented by black ovals. **(E)** Gene fractionation bias in the three *H. incana* sub-genomes. The ancestral tPCK genomic blocks were used as reference. Gene retention (%) was calculated in sliding windows of 100 genes across the tPCK genomic blocks based on all identified syntenic genes.

We further elucidated the WGD/WGT history of *H. incana* by fitting the distribution of *Ks* values (the ratio of number of substitutions per synonymous site) from its WGD/WGT-derived gene pairs (**Supplemental Fig. S11**) using a Gaussian Mixture Model (GMM) (Qiao et al., 2019). Here, we revealed three major *Ks* peaks corresponding to the three recent WGD/WGT events in the genome (**Fig. 2C**). The youngest peak (in purple) represents the more recent *Br-α* WGT event that was shared among the Brassiceae species (Brassiceae tribe-specific, ∼15.9 MYA), while the two more ancient peaks (in yellow and blue) respectively represent the more ancient *At-α* WGD event (Brassicaceae family-specific, ∼35 MYA, i.e., late Eocene to early Oligocene epochs) and *At-β* WGD event (shared among Brassicaceae and other families in the Brassicales order, 50-60 MYA, i.e., the mid-Palaeocene to early Eocene epochs) (Jiao et al., 2011; Kagale et al., 2014; Liu et al., 2014; Edger et al., 2018). The fitted peaks are consistent with those identified in the *Brassica* genomes previously reported using the same method (Qiao et al., 2019; Yim et al., 2022; Hoang et al., 2023).

The Brassiceae genomes are known to be derived from the ancestral tPCK karyotype (Schranz et al., 2006; Lysak et al., 2016). We thus used the updated ancestral genomic blocks of the *Brassica* genomes (He et al., 2021) to determine the intervals and boundaries of the 26 ancestral genomic blocks in the *H. incana* genome (**Fig. 2D**, **Supplemental Table S10** and **Supplemental Fig. S12** and **S13**). We were able to identify most of the triplicated blocks (except small blocks G, S and V1) within the *H. incana* genome that are syntenic to the ancestral genomic blocks. Based on the gene retention rate, these triplicated genomic blocks were then classified into three sub-genomes, the LF, MF_1_ and MF_2_ (**Fig. 2D**). The average retention rates of these sub-genomes compared to the ancestral tPCK genome were 72%, 52% and 40% for LF, MF_1_ and MF_2_, respectively (**Fig. 2E**). This sub-genome biased fractionation, as a result of the two-step polyploidization process, has been reported in other Brassiceae genomes (Cheng et al., 2012; Liu et al., 2014; Perumal et al., 2020; Cho et al., 2022; Yang et al., 2023), underscoring the notion that the *H. incana* genome was also derived from the common tPCK ancestral genome of the Brassiceae tribe. By reconstructing an intermediate common ancestral genome of the six Brassiceae species, we found that, compared to the genomes of other Brassiceae species, the *H. incana* genome showed a similar level of rearrangement as that of *B. nigra* and *S. arvensis* (36-41 fissions and 38-44 fusions), but higher than that of *B. rapa*, *B. oleracea* and *R. sativus* (29-32 fissions and 31-33 fusions) (**Supplemental Fig. S14**).

Taken altogether, the updated genome assembly of *H. incana* allowed us to elucidate its genome structure, WGD/WGT history, and genome evolution at the sub-genome level. The *H. incana* genome was derived from the recent Brassiceae-specific *Br-α* WGT event that resulted in three distinct sub-genomes that display differential gene retention rates. These sub-genomes were shown to have originated from the ancestral tPCK karyotype of the Brassiceae, similar to that of the other genomes from the *Brassica*, *Raphanus* and *Sinapis* genera.

### Phylogenomic analysis and genome synteny reveal close relationships among *Brassica (A/C/B), Hirschfeldia (H)*, and *Raphanus (R)*

Several chromosome-level genome assemblies of Brassiceae species have been released, including those outside of the *Brassica* “Triangle of U”, such as *R. sativus* (Cho et al., 2022; Xu et al., 2023) and *S. arvensis* (Yang et al., 2023). Large-scale comparative genomics and phylogenetic analyses have provided insights into relationships among these species. However, due to the rampant hybridization among Brassiceae species, that hinders phylogenetic inference, a consensus phylogenetic tree for this tribe is still not resolved. There are on-going debates as to whether to include species from the *Raphanus* and *Sinapis* genera in the *Brassica* genus (Cho et al., 2022; Yang et al., 2023). While the relationships between the *Brassica* A/C and B genome types appear to be more consistent in published nuclear phylogenetic trees, this is not the case for *R. sativus* (Huang et al., 2016; Cho et al., 2022; Yang et al., 2023) and *H. incana* (Huang et al., 2016; Garassino et al., 2022; Guerreiro et al., 2023); as these two species were placed closer to either *Brassica* A/C or B genome type species. Interestingly, Jeong et al. (2016) and Cho et al. (2022) revealed that the *R. sativus* genome displayed intermediate characteristics between A/C and B genome types, which highlights the possibility that *R. sativus* may have been derived from the *Brassica* genus (i.e., a sibling of the *Brassica* species).

We reconstructed a phylogenetic tree using 1,504 single-copy genes identified across 19 genome accessions from a total of 10 species in an attempt to resolve the discrepancy in the phylogenetic placement of *H. incana* and *R. sativus* (**Fig. 3A**). The 10 species included several from the “Triangle of U” *Brassica* species (A, B and C genome types), *Raphanus*, *Sinapis* and *Hirschfeldia* genera (Liu et al., 2014; Parkin et al., 2014; Jeong et al., 2016; Belser et al., 2018; Perumal et al., 2020; Guo et al., 2021; Cho et al., 2022; Xu et al., 2022; Guerreiro et al., 2023; Xu et al., 2023; Yang et al., 2023; Zhang et al., 2023). We also included *B. tournefortii*, and a *Sinapis* sp. (the latter formerly labeled as the *H. incana* accession HIR3), two species that are closest to *H. incana* for which draft genome sequences are available, and the *H. incana* HIR1 accession (Guerreiro et al., 2023). Our phylogenetic tree revealed that all *Brassica* genome types A and C formed a monophyletic clade that represents the *Rapa/Oleracea* clade of the Brassiceae, while *S. arvensis* and all other *B. nigra* B genomes formed another monophyletic clade. This is consistent with previous studies which suggested a close relationship between *B. rapa* and *B. oleracea* (Cheng et al., 2014), and between *B. nigra* and *S. arvensis* (Yang et al., 2023). Overall, our results support *R. sativus* and *H. incana* (referred to collectively as R/H genome type) being closer to the *Brassica* B genome type (i.e. *B. nigra*) than to the A/C type (*B. rapa/B. oleracea*). Notably, our tree topology is consistent with that in Cho et al. (2022) regarding the placement of *R. sativus* and *Brassica* species, and with that in Garassino et al. (2022) regarding the placement of *H. incana* and *Brassica* species. However, our tree topology is different from the tree constructed by Yang et al. (2023) in which *R. sativus* is grouped together with the *Brassica* A/C genome type and inconsistent with the tree topology proposed by Huang et al. (2016) in which *R. sativus* and *H. incana* are grouped together with the *Brassica* A genome type. It is also important to note that a different topology exists. For example, Guerreiro et al. (2023) reported a species tree of Brassiceae species that shows *R. sativus* and *H. incana* in a distinct clade, separated from all three *Brassica* A/C/B in another clade. The incongruency among nuclear species trees might have stemmed from different gene sets included in different studies, and this will be analyzed and discussed in the next section.

**Figure 3.**
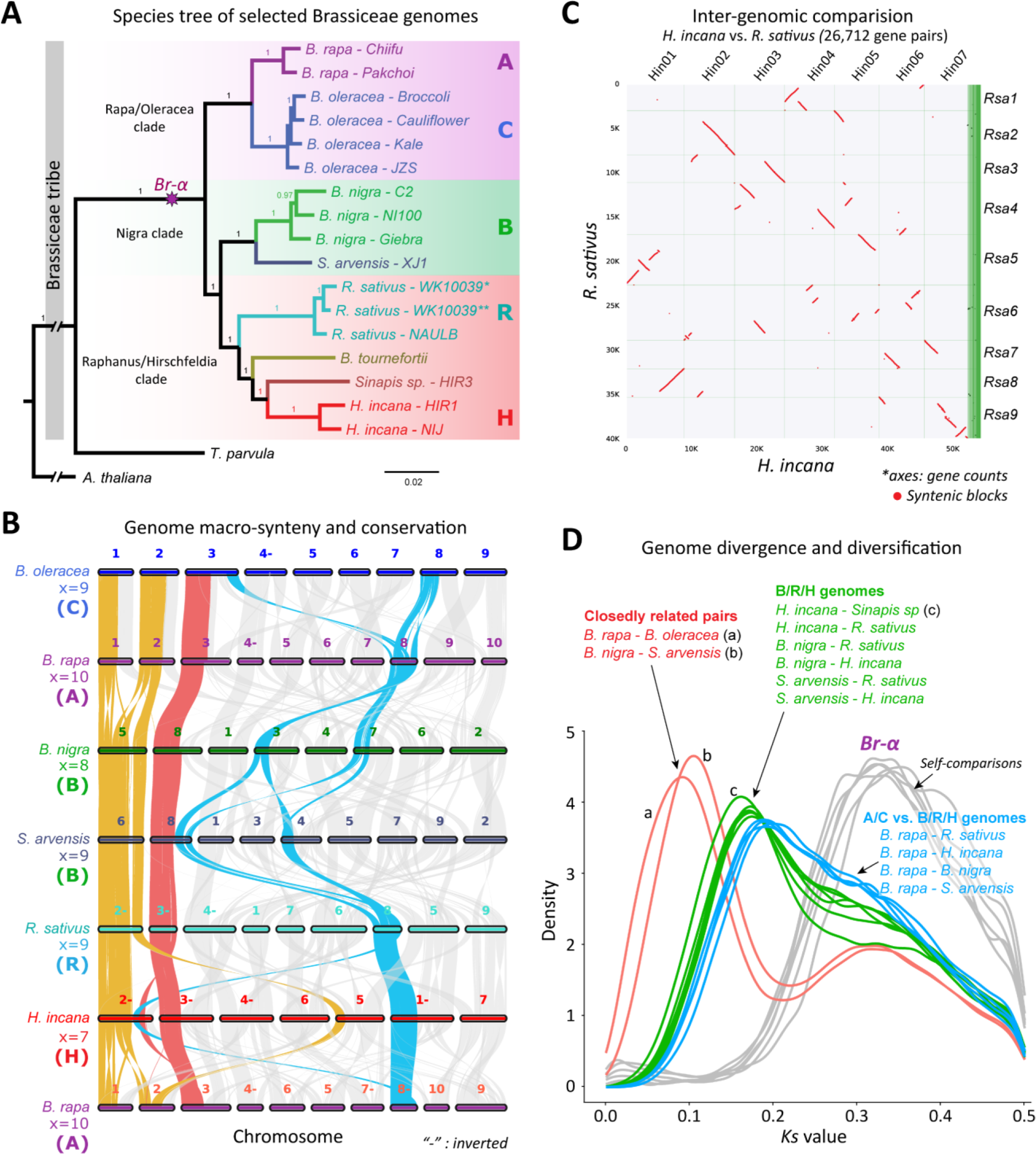
Relationships of *H. incana* and other Brassiceae species. **(A)** Species tree of 10 selected Brassiceae species including several genome accessions for each species. The tree was reconstructed based on 1,504 single-copy genes identified across the selected genomes. It was rooted using the *A. thaliana* (and *Thellungiella parvula*) genomes as outgroup. Supporting values at each node are bootstrap scores. (**B**). Genome macro-synteny plot of six representative Brassiceae genomes for each group in Panel **A**, showing intermediate characteristics of *R. sativus* and *H. incana* genomes compared to *Brassica* A/C/B genome types. For example, a conserved syntenic region across all six species (red); a conserved region that is well retained in *Brassica* B genome type, *R. sativus* and *H. incana* (yellow); and a conserved region that is well retained in *Brassica* A/C genome type, *R. sativus* and *H. incana* (blue). Syntenic blocks (*minspan = 30 genes*) were used for each pair-wise comparison among the selected genomes. Chromosome length is not at scale. Chromosome order is rearranged based on the syntenic relationship between two adjacent genomes. “-” denotes inverted sequence. **(C)** Inter-genomic synteny comparison between the *H. incana* and *R. sativus* genomes. Syntenic blocks (*minspan = 30 genes*) between genomes were aligned. Only true orthologs are shown. Axes show gene count for each genome. **(D)** Genome divergence among the selected species. *Ks* distributions were colored for four groups, comparisons among closely related genome pairs (red), among B/R/H genomes (green), among A/C and B/R/H genomes (blue) and self-comparisons of the included genomes (grey). The self-comparison of each highlights the recently shared *Br-α* WGT event in these genomes (similar to that in Figure 2C), while pairwise genome comparisons highlight species divergence. For panels **B-D** and other results where only one genome accession was included for each species, the genomes of *A. thaliana* Col-0, *B. rapa* Chiifu, *B. oleracea* JZS, *B. nigra* NI100, *S. arvensis* XJ1, *R. sativus* NAULB and *H. incana* NIJ were used. The *R. sativus* WK10039* and ** denotes v1.0 and v2.0 assemblies, respectively.

We also analyzed global genome synteny among six selected Brassiceae species for which chromosome-level assemblies are available (**Fig. 3B**). These six genomes formed three pairs that showed a high level of synteny to each other, including *B. rapa* (x=10) – *B. oleracea* (x=9), *B. nigra* (x=8) – *S. arvensis* (x=9) and *H. incana* (x=7) – *R. sativus* (x=9). These represent an array of chromosome numbers that was derived from the ancestral tPCK karyotype (x = 7, Schranz et al. (2006); Lysak et al. (2016)). Interestingly, the *H. incana* genome exhibited a greater degree of synteny with the *R. sativus* genome than with any other genome (**Fig. 3B, C** and **Supplemental Fig. S15**). We find support for intermediate genome structure of *H. incana* and *R. sativus* in relation to the *Brassica* A/C and B genome types. For example, a highly syntenic region (yellow) was detected across *B. nigra* (chr5), *S. arvensis* (chr6), *H. incana* (chr2), *R. sativus* (chr2), while it was rearranged in both *B. rapa* and *B. oleracea* genomes (chr1 and chr2) (**Fig. 3B**). On the other hand, another highly syntenic region (blue) is conserved across *B. rapa* (chr8), *B. oleracea* (chr8), *H. incana* (chr1) and *R. sativus* (chr8), while it was rearranged in *B. nigra* and *S. arvensis*. We believe these have a shared evolutionary origin, but their independent origin through breakpoint reuse (Li et al., 2016) cannot be ruled out. We also observed several blocks that are well conserved across the six genomes (e.g., red block), reflecting their common ancestry. By analyzing the distribution of *Ks* values of orthologous genes among the selected genomes, we reconstructed their history of lineage divergence after their shared *Br-α* WGT event (**Fig. 3D**). After divergence from the *Brassica* A/C, the *Brassica* B species was then separated from the *Raphanus* and *Hirschfeldia* species. The smallest *Ks* peaks of comparisons between *B. rapa* and *B. oleracea* (A/C genomes) or between *B. nigra* and *S. arvensis* (B genomes) suggest that they are closer to each other than to any other species and the split of species in these two pairs occurred more recently. Among all included R/H genomes, *Sinapis sp.* was the closest species to *H. incana*, even though their *Ks* peak was only slightly smaller than that of other R vs. H or B vs. R/H genome comparisons. Overall, in light of the phylogenetic and genome synteny analyses, our results support the proposal to include the *Raphanus* and *Hirschfeldia* genera into an expanded *Brassica* genus.

### Intermediate characteristics of *Hirschfeldia* and *Raphanus* explain their incongruent phylogenetic placement in relation to the *Brassica* species

Based on the plastid phylogeny, *Raphanus* and the *Brassica* A/C species belong to the *Rapa/Oleracea* clade, while *Hirschfeldia* and *Brassica* B species belong to the *Nigra* clade within the Brassiceae tribe (Arias and Pires, 2018; Guerreiro et al., 2023). As discussed earlier, recent nuclear-based phylogenetic trees for the Brassiceae species still display different topologies regarding the placement of *R. sativus* and *H. incana* in relation to the *Brassica* A/C and B genomes (Cho et al., 2022; Garassino et al., 2022; Guerreiro et al., 2023; Yang et al., 2023). Our new phylogenetic analysis recovered a tree topology in which *R. sativus* and *H. incana* were placed closer to the B genome than to the A/C genome types (**Fig. 3A**). Because their genome structures also suggest that *R. sativus* and *H. incana* exhibit intermediate characteristics between *Brassica* A/C and B genome types, we wondered if hybridization/introgression could explain the discrepancy in the nuclear phylogenetic placement of these species among published studies. Additionally, these genomes underwent the Brassiceae-specific *Br-α* WGT event followed by a subsequent gene fractionation and rediploidization, making them have three distinct sub-genomes (LF, FM_1_ and MF_2_) of different gene retention rates. It is possible that due to differential gene loss rates, different paralogous genes were retained during these processes, leading to the occurrence of pseudo-orthologs, which are single-copy homologs originally from different sub-genomes (Cheng et al., 2014; Smith and Hahn, 2021). The inclusion of such pseudo-orthologs can have profound effects on species tree estimation, in particular if gene loss occurs along internal branches (Xiong et al., 2022). We therefore also checked if the observed incongruency among nuclear species trees is potentially due to different contributions of sub-genome specific paralogous genes to sets of single-copy orthologs that were used for phylogenetics.

The first underlying question is that, as these genomes contain genetic material originating from both A/C and B progenitors, different gene selection methods might have recovered different gene proportions from A/C and B types that could affect the phylogenetic inference. We thus analyzed gene tree topologies of all single-copy orthologs identified by OrthoFinder that are conventionally used to reconstruct species trees. To simplify, here we included only six genomes of three *Brassica* A/C/B types, *R. sativus*, *H. incana*, and *A. thaliana,* which resulted in a species tree of similar topology to that in **Fig. 3A** for these species. A total of 5,675 single-copy genes were identified among these genomes (**Supplemental Table S11**) and their topologies are shown in **Fig. 4A**. Aside from the major topology (blue), other topologies were also recovered (green and red). We detected more conflicts in nodes involving *B. nigra*, *R. sativus* and *H. incana,* with concordance between 42% and 52% (**Fig. 4A** and **Supplemental Fig. S16**). It is noteworthy that, based on the plastid tree, *R. sativus* probably received the maternally inherited chloroplast genome from an A/C type mother while *H. incana* received it from a B genome type mother. Here, following the topology quantification approach of Forsythe et al. (2020), we counted gene trees based on three topologies that represent the plastid topology (**Fig. 4B**) and the two most dominant nuclear topologies (**Fig. 4C, D**) corresponding to two major species trees in published studies. Our results (**Fig. 4E**) revealed that among 3,566 filtered single-copy gene trees (bootstrap support ≥50%), the most common topology (29.4%) supported *R. sativus/H. incana* being closer to the *Brassica* B genome (nuclear topology 1), while 21.2% of trees supported them being closer to A/C type (nuclear topology 2). Only 4.4% of gene trees followed the plastid topology, which could be those nuclear genes that shared the evolutionary history of the chloroplast genes as a result of selection for cyto-nuclear compatibility (Forsythe et al., 2020). Nevertheless, our results might explain the incongruency observed in published species trees regarding the placement of *R. sativus* and *H. incana* in relation to the species within the *Brassica* “Triangle of U”.

**Figure 4.**
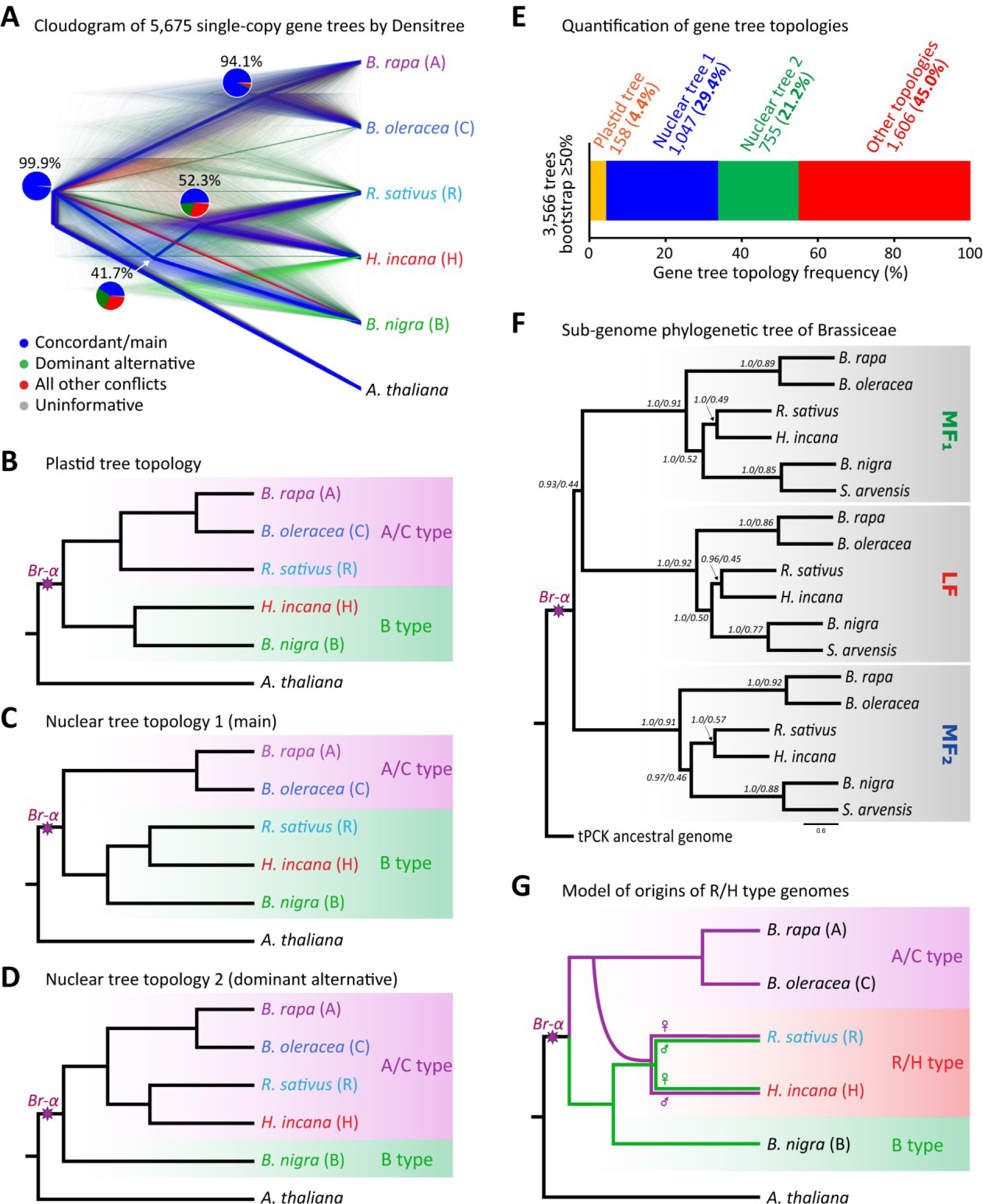
Species tree incongruency and gene tree topology analysis of selected Brassiceae genomes. **(A)** Cloudogram of gene trees derived from a total of 5,675 single-copy orthologs identified among five selected Brassiceae and the *A. thaliana* genomes showing concordant (main, blue), dominant alternative (green) and other conflict (red) topologies. Pie charts derived from the PhyParts analysis (Smith et al., 2015) show percentage of trees supporting main/alternative/conflict topologies at each node. The tree was rooted using the *A. thaliana* as outgroup. **(B-D)** Three focal gene tree topologies that were analyzed among the above species including plastid, main nuclear, and dominant alternative nuclear topologies, respectively. **(E)** Quantification of gene tree topologies (in panels **B-D**) in subset of 3,566 filtered gene trees from the initial set shown in panel **A**, keeping only gene trees with bootstrap support ≥50%. **(F)** Sub-genome tree of six selected Brassiceae species. The tree was reconstructed using the species-tree approach based on 90 genes located on the tPCK genomic block F found across all sub-genomes of the six genomes. The tree was rooted using the ancestral tPCK genome as outgroup. Supporting values at each node are posterior probability and quartet scores, respectively. Branch length represents coalescence units. **(G)** A model for the possible origins of the *R. sativus* and *H. incana* genomes deriving from the *Brassica* A/C and B genome types. The maternal origins were determined based on the plastid tree topology (panel **B**), while the species tree is based on the major nuclear topology (panel **C**). The genomes of *A. thaliana* Col-0, *B. rapa* Chiifu, *B. oleracea* JZS, *B. nigra* NI100, *S. arvensis* XJ1, *R. sativus* NAULB and *H. incana* NIJ were used.

Another potential factor that might affect the species tree topologies based on single-copy orthologous genes is that these genes could have originated from different sub-genomes. Indeed, in the 5,675 single-copy orthologs in the *H. incana* genome, we found that approximately 46.7% are located on the LF sub-genomes, while 24.7% and 15.6% are on MF_1_ or MF_2_, respectively (**Supplemental Table S12**). This suggests that, for genomes that underwent WGD/WGT events followed by re-diploidization, single-copy paralogs from different sub-genomes, if taken as orthologs, could potentially contribute to nuclear species tree discrepancy. More specifically, a significant proportion of these single-copy orthologs could be pseudo-orthologs instead of true orthologs, which could be problematic for species tree inference. Interestingly, when comparing our single-copy genes with the sets of single-copy loci commonly used in phylogenomics (B764 - Brassicaceae-specific dataset: Nikolov et al. (2019), Hendriks et al. (2023); and A353 - Angiosperms universal bait set: Johnson et al. (2019)), we found that a higher proportion of genes (55%) was from the *H. incana* LF sub-genome, potentially due to the more conserved nature of loci that are single-copy across a larger evolutionary timeframe. It is important to note that, while we focused on two main potential factors, introgression and single-copy pseudo-orthologs, our analysis cannot rule out that incomplete lineage sorting could be an alternative hypothesis and additional factors to explain the species tree incongruency.

Recently, Walden and Schranz (2023) recommended using a synteny-based approach for a more reliable identification of true orthologs for phylogenetic studies. We tested if this could aid in resolving the issues with OrthoFinder single-copy orthologs in this case. As this approach utilizes all available triplicated gene copies, it could recover both species trees and sub-genome trees using the same set of markers. For this analysis, we focused on 90 syntenic orthologous gene sets identified across sub-genomes of six selected Brassiceae species that are located on the ancestral tPCK genomic block F (**Supplemental Table S13** and **Supplemental Fig. S17**). Both sub-genome and species trees derived from these 90 selected syntenic orthologs (**Fig. 4F** and **Supplemental Fig. S18**) are consistent with our previous species tree based on OrthoFinder single-copy orthologs (**Fig. 3A**) and with the main nuclear topology (**Fig. 4C**). In the sub-genome tree, all sub-genome types across species were shown to be more similar to each other than different sub-genome types within the same species, and each type formed a monophyletic clade, with the LF and MF_1_ sub-genomes being grouped closer to each other than either to the MF_2_ sub-genome. The results indicate that a synteny-based approach could be used as a reliable method for species tree reconstruction, and further corroborate that the use of single-copy orthologs that were derived from different sub-genomes could potentially change phylogenetic outcomes.

Finally, we proposed a model of the origins of *Raphanus/Hirschfeldia* (R/H) genome types from the *Brassica* genomes (**Fig. 4G**). It is likely that both genome types contain genetic materials from the *Brassica* A/C and B progenitors. As suggested by the plastid-based phylogenetic tree (**Fig. 4B**), *R. sativu*s might have a *Brassica* A/C type mother while *H. incana* might have a B type mother, indicating that these two species might have been the result of reciprocal hybridizations.

### Whole-genome duplication retained genes that show a sub-genome expression bias are associated with distinct biological processes

Compared to its Brassiceae relatives, the C_3_ species *H. incana* was reported to display high photosynthesis rates under high-light conditions (Canvin et al., 1980; Garassino et al., 2022) and the ability to grow in lead contaminated soils (Auguy et al., 2013; Hasnaoui et al., 2022). We hypothesized that this was a result of differential gene retention of WGT gene copies, possibly in a sub-genome biased fashion, followed by neo-/sub-functionalization in the *H. incana* genome that gave rise to its ability to accumulate biomass and thrive in such conditions. To provide a comprehensive analysis of triplicated genes in the *H. incana* genome and their potential associations with its adaptive evolution of high-photosynthesis traits under high-light conditions, we studied three gene categories: triad, dyad and single-copy genes. These loci are those which respectively retained three, two or one homologous gene copy of the triplicates from the ancestral tPCK copy after the *Br-α* WGT event.

First, using sub-genome information in **Fig. 2D**, we identified a total of 2,103 triads and 6,457 dyads. The number of triads identified in *H. incana* is similar to that of other Brassiceae genomes reported in Yang et al. (2023), ranging from 1,531 to 2,183. GO biological process and KEGG pathway analyses of the 2,103 triads and 6,457 dyads showed an enrichment (a false discovery rate, FDR, corrected *p* ≤0.05) for genes related to plant organ development, growth, shoot system development and morphogenesis, responses to hormones and stimuli, as well as photosynthesis and carbon metabolism (**Supplemental Figs. S19** and **S20**). Many of these enriched terms overlap those from an analysis of upregulated genes under high-light conditions (**Supplemental Table S14**). The results suggest that WGT genes retained for these processes followed by neo-/sub-functionalization could have facilitated the adaptive responses to high-light conditions in *H. incana*.

Garassino et al. (2023) generated transcriptome data of *H. incana* whole canopies from plants grown under contrasting low-light and high-light conditions (**Supplemental Fig. S21**). We utilized this data to analyze gene expression patterns of the identified triad and dyad genes, as shown in **Fig. 5A**. To test if there is an expression bias among the identified triads originating from different sub-genomes, we compared pairwise expression patterns (LF vs. MF_1_, LF vs. MF_2_ and MF_1_ vs. MF_2_) using the method outlined in Cheng et al. (2012) and a threshold |fold-change| ≥ 2, *p* ≤0.05. Of the total 2,103 triads, we further filtered out lowly expressed genes to obtain 1,439 triads that had a reliable expression level for cross sub-genome comparisons. Of these, we found that about 40% of triad genes showed dominance in the LF compared to MF_1_ sub-genome under low and high-light conditions, while it was only 25-27% of that in the MF_1_ more dominant over LF sub-genome (**Fig. 5B** and **Supplemental Table S15**). Likewise, when comparing between the LF and MF_2_ sub-genomes, 39% and 25-26% genes showed dominance for the respective sub-genomes. However, the percentage of dominant genes between the MF_1_ and MF_2_ genomes were more similar, ranging from 31-32% and 33-34%, respectively. A similar pattern was observed when comparing a total of 3,722 filtered dyad genes (**Fig. 5C** and **Supplemental Table S15**). When all three sub-genomes were compared, it was found that, of the 1,439 filtered triads, 49% and 28% could be classified as dominant in LF and MF_1_/MF_2_ sub-genomes, respectively (**Supplemental Table S16**). The LF-dominant triad genes were most enriched (FDR-corrected *p* ≤0.05) for “translation” and many GO terms related to “response to hormones/endogenous stimuli” and “reproduction”; while the MF_1_/MF_2_-dominant genes were most enriched for “cell growth”, “developmental growth involved in morphogenesis”, “developmental growth”, “flavonoid biosynthesis process” and “response to light stimuli” (**Fig. 5D**). Among the 3,722 filtered dyads, apart from shared enriched terms related to “response to light stimuli”, “carboxylic acid metabolic”, the LF-dominant dyad genes were most enriched for terms related to “organelle organization” and “protein localization to organelle”; while MF_1_/MF_2_-dominant genes were enriched for “homeostasis”, and “ion transport” processes (**Fig. 5D**).

**Figure 5.**
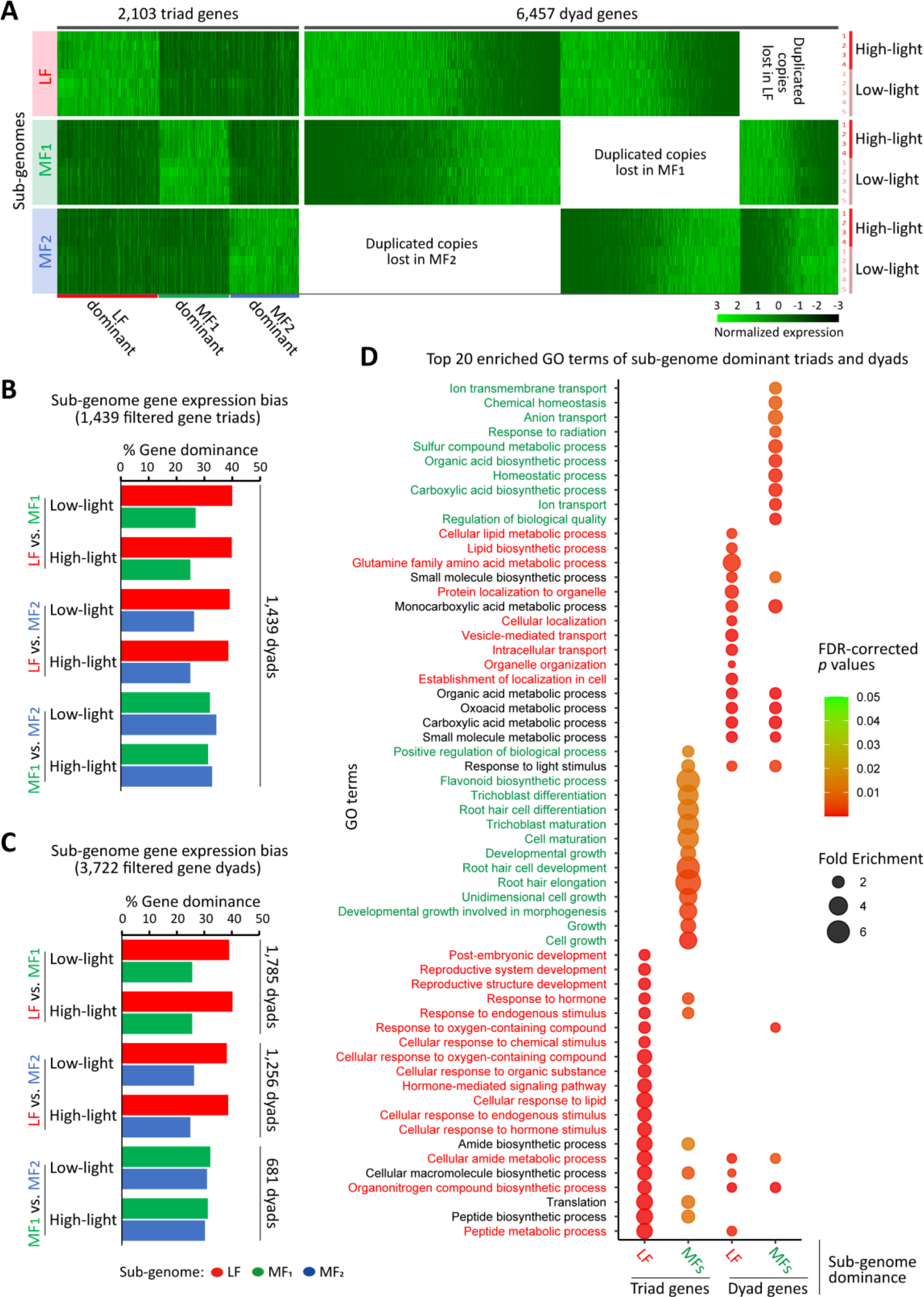
Analysis of sub-genome biased expression of WGD triad and dyad gene copies in the *H. incana* genome. **(A)** Gene expression of the 2,103 identified triads (three syntenic triplicated copies) and dyads (two syntenic copies) in two conditions, low (5 replicates) and high-light (4 replicates), taken from whole canopy transcriptomes (Garassino et al., 2023). Gene expression data are presented based on sub-genomes and sorted based on the expression levels across the three sub-genomes. For each gene (column), the expression data were column-normalized across all samples. **(B, C)** Gene expression dominance in different *H. incana* sub-genome comparisons, LF-MF_1_, LF-MF_2_ and MF_1_-MF_2_ calculated for 1,439 triad and 3,722 dyad gene sets obtained from the initial 2,103 and 6,457 respective gene sets in panel **A** after filtering out lowly/non expressed genes. Gene expression dominance between a pair of duplicated genes was determined using a threshold |fold-change| ≥2, *p* ≤0.05. The numbers of genes used are indicated next to the bar charts. **(D)** Gene ontology (GO) enrichment analysis of triad and dyad genes that showed dominance in the LF and MF_1_/MF_2_ sub-genomes, respectively. Top 20 most significant GO terms (*FDR*-corrected *p ≤ 0.05*) for each gene set are shown. For GO enrichment of the total 2,103 triad and 6,457 dyad genes, see **Supplemental Figs. S19** and **S20**.

While we only tested sub-genome gene expression bias in leaf tissues grown at two conditions in *H. incana*, the results are consistent with the observations in other Brassiceae genomes (Cheng et al., 2012; Yang et al., 2023). There might be different gene copies within each triad and dyad that are expressed in a tissue-specific manner. However, this sub-genome expression bias was also reported in different tissues, for example, in leaf, stem and root tissues of *B. rapa* (Cheng et al., 2012); or in leaf and stem tissues of six Brassiceae genomes (Yang et al., 2023). Altogether, our results indicate a bias in sub-genome gene retention and expression (i.e., LF more dominant over MF_1_/MF_2_) in the *H. incana* genome. Genes that show a sub-genome expression dominance appeared to be associated with different biological processes. More specifically, the LF-dominant genes were most enriched for terms related to responses to stimuli/hormones and organelle organization, while MF_1_/MF_2_-dominant genes were most enriched for growth/development and photosynthesis-related.

Next, we focused on a total of 6,769 single-copy genes that we could confidently assign to the three sub-genomes, of which 3,527 (52%) were found on the LF, 1,932 (29%) on the MF_1_ and 1,310 (19%) on the MF_2_ sub-genome, respectively. These genes were found to be single-copy in both *A. thaliana* and *H. incana* genomes and were derived from a total of 7,952 originally identified in our OrthoFinder analysis. Interestingly, GO enrichment analysis of these gene sets suggested an enrichment (FDR-corrected *p* ≤0.05) for several organelle-related terms, “RNA modification”, “chromosome segregation”, “embryo development”, “cell cycle” and “reproduction” within single-copy genes from the LF sub-genome; “mismatch repair”, “reciprocal meiotic/homologous recombination” and “plastid organization” within those from MF_1_; and “replication fork processing”, “DNA replication maintenance”, “response to DNA damage stimulus”, “DNA repair”, “RNA modification” and “DNA metabolic process” within those from the MF_2_ sub-genome (**Supplemental Fig. S22**). The result is consistent with findings in previous studies that, unlike genes related to transcription factors, ribosomal proteins and kinases; genes related to DNA repair and organelle-targeted pathways tend to return to single-copy after the WGD/WGT event (De Smet et al., 2013). This result highlights the importance of studying the mechanisms that lead to the sub-genome biased retention of single-copy genes, because many chloroplast-targeted genes are involved in the photosynthesis machinery.

Evidence is accumulating that polyploidy could potentially aid plants in adapting to (and thriving in) new challenging environments and stressful climates (Van de Peer et al., 2017; Stevens et al., 2020). For example, Feng et al. (2024) found a strong selection on three-copy retained gene families associated with adaptive response to new environments in the mangrove tree (*Sonneratia alba*). These include genes related to root development and salt tolerance that enhance plant adaptive response to intertidal zones. In relation to photosynthesis traits, several studies, including Wang et al. (2009b) and Hoang et al. (2023), showed the contribution of WGD to the evolution of C_4_ photosynthesis from the C_3_ ancestral state in two evolutionary distant families, Poaceae and Cleomaceae, respectively. Our results on retained WGT genes and their sub-genome dominance, especially those related to plant response to endogenous stimuli, morphogenesis, development, organelle organization, chloroplast-targeted pathways and photosynthesis might reflect important gene families that were involved in the evolution that led to the high-photosynthesis trait at high-light intensity in *H. incana*.

### Analysis of gene families related to leaf physio-biochemical-anatomical changes that potentially facilitate adaptation to high-light intensity in *Hirschfeldia incana*

As our previous analyses suggested several gene groups that are related to adaptive changes that might explain the high-photosynthesis traits in *H. incana* and its ability to withstand high-light conditions, our next focus was particularly on the expression of genes involved in the key changes found in the plants grown under high light. These key changes were based upon the recent findings on the significant physiological, biochemical and anatomical differences in the *H. incana* leaf tissues in response to high light compared to its relatives. More specifically, the changes consist of (1) a higher average gross CO_2_ assimilation rate at high irradiance (Garassino et al., 2022), (2) a higher endoreduplication (endopolyploidy) level (**Fig. 6A** and **Supplemental Table S17**), (3) a smaller antenna size and more chloroplasts of smaller size (Caracciolo et al., *in prep.*, and **Fig. 6B**), and (4) the development of thicker multilayer palisade mesophyll cells, a higher vasculature volume and mesophyll surface area (Retta et al., *under review*). Among these, to our surprise, when compared with *B. rapa* and *B. nigra*, *H. incana* showed a much higher endoreduplication level in mature leaves under high light (50% vs. ≤10% nuclei at 8× ploidy level, **Fig. 6A**). Our hypothesis was that these changes are the results of differential expression of genes associated with the photosynthesis machinery (i.e., response to high-light stimuli and photosystem proteins), cell division (i.e., cell cycle, endoreduplication and plastid division) and leaf development (mesophyll formation, vein development and hormone signaling pathways). To this end, we identified genes differentially expressed between low-light and high-light conditions in the whole canopy transcriptome data of *H. incana* (Garassino et al., 2023) using our updated gene models. A total of 2,012 up- and 2,126 downregulated genes were found using a threshold FDR corrected *p*-value ≤0.05 and |fold-change| ≥2 (see **Supplemental Table S18** for a list and **Supplemental Table S14** for GO enrichment analysis).

**Figure 6.**
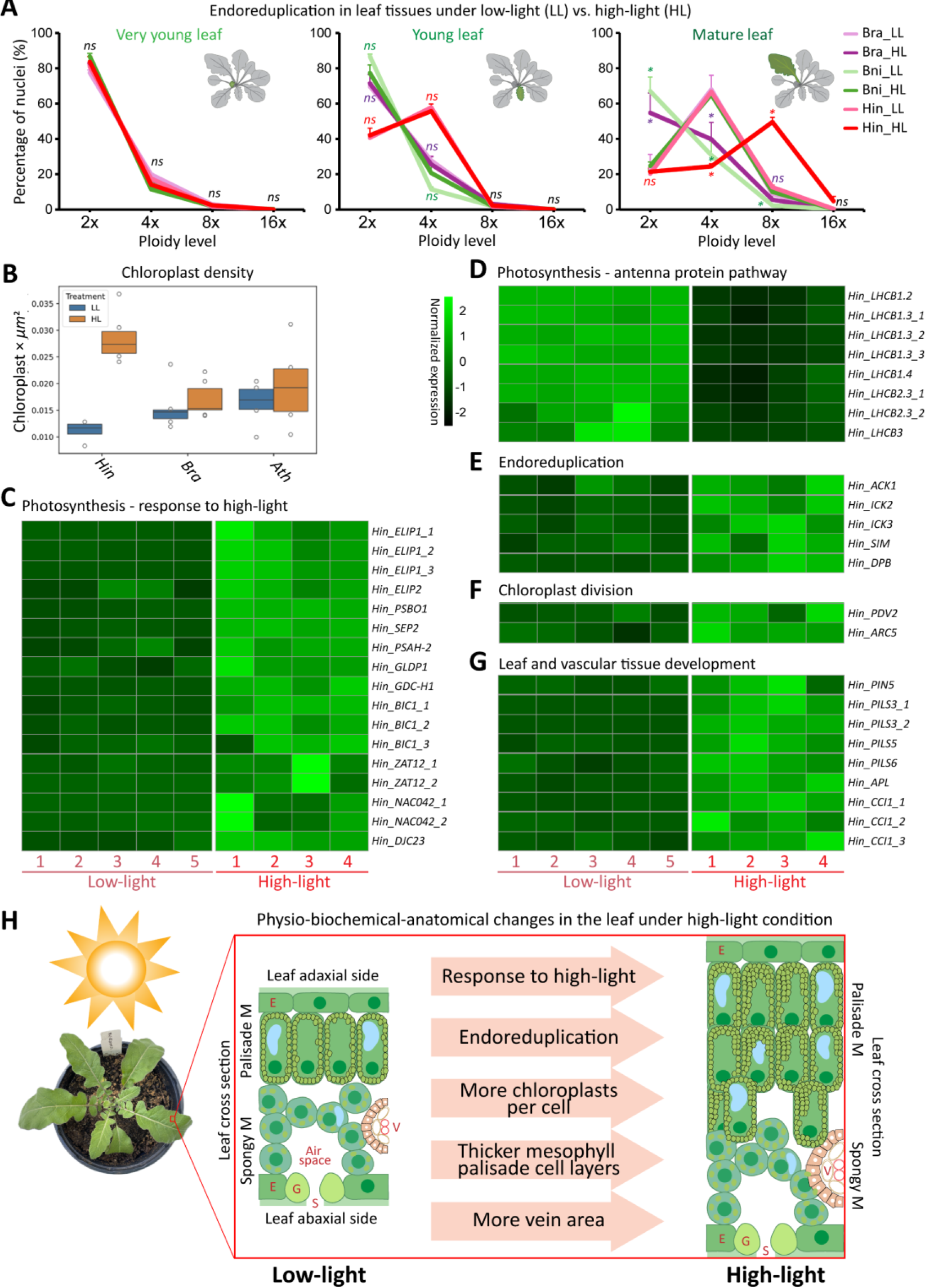
Analysis of gene families associated with adaptive evolution to high-light conditions in the *H. incana* genome. **(A)** Endoreduplication level of plants grown under low-light and high-light conditions measured by flow cytometry. For each timepoint, three samples were used. Data are presented as mean ± SEM (standard error of the mean). Error bars indicate SEM. Ns: non-significant. Asterisk (*): statistical significance at *p* ≤0.05 by Student’s *t*-test. **(B)** Chloroplast density of plants grown under low-light and high-light. **(C-G)** Expression analysis of gene families associated with physio-biochemical-anatomical adaptive changes that likely facilitate the high photosynthesis in *H. incana*, including genes related photosynthesis, endoreduplication, chloroplast division, leaf and vascular development. Only genes that showed an upregulation (FDR corrected *p*-value ≤0.05 and |fold-change| ≥2) in the whole canopy transcriptomes of low and high-light conditions (Garassino et al., 2023) were included. Gene expression data are TPM values and were row-normalized (per gene). **(H)** Model of *H. incana* leaf physio-biochemical-anatomical adaptive changes in response to high-light conditions. E denotes epidermis. G: guard cells, M: mesophyll, S: stomata, V: vasculature. Ath: *A. thaliana*, Bra: *B. rapa*, Bni: *B. nigra* and Hin: *H. incana*. LL: low light, HL: high light.

Interestingly, among the upregulated genes related to photosynthesis and response to high-light intensity (**Fig. 6C** and **D**), several genes encoding for transcription factors and proteins were found, including *ELIPs (EARLY LIGHT-INDUCIBLE PROTEINs)*, *BIC2 (BLUE-LIGHT INHIBITOR OF CRYPTOCHROMES 2), ZAT12/RHL41 (RESPONSIVE TO HIGH LIGHT 41)*, *NAC042 (NAC DOMAIN CONTAINING PROTEIN 42)*, and *DJC23 (DNA J PROTEIN C23)*. Besides ELIPs that is known to have roles in photoprotection and chlorophyll biosynthesis (Hutin et al., 2003), these might be genes that contribute to the adaptation to the high-light conditions and facilitate the high photosynthetic efficiency in *H. incana*. For example, ZAT12 was shown to play a central role in high-light acclimation and response to oxidative stress in *A. thaliana* (Davletova et al., 2005), while the transcription coactivator BIC1 promotes brassinosteroid signaling and plant growth (Yang et al., 2021). NAC042, a reactive oxygen species-responsive transcription factor enhances stress tolerance and delay senescence (Wu et al., 2012), while *DJC23* is involved in optimizing photosynthetic reactions (Chen et al., 2010). Additionally, we noticed that most of the genes encoding photosystem II LHCBs (LIGHT HARVESTING CHLOROPHYLL A/B-BINDING PROTEINs) and photosystem I LHCAs (PHOTOSYSTEM I LIGHT HARVESTING COMPLEX PROTEINs) were, to some extent, suppressed by high-light conditions. However, we found only *LHCBs* genes (i.e., *LHCB1.2, LHCB1.3s, LHCB1.4, LHCB2.3s* and *LHCB3*) to be significantly downregulated (**Fig. 6D**). This might explain the reduction of functional antenna size of photosystem II but not that of photosystem I, as observed in Caracciolo et al., (*in prep.*).

Among the upregulated genes related to cell division, we found several interesting genes associated with cell cycle control “CYCLIN-dependent protein kinase inhibitor activities - CKI” (**Fig. 6E**) that might be responsible for the high endoreduplication level by inhibiting cell division (Vieira et al., 2014). These include *ACK1 (ARABIDOPSIS CDK INHIBITOR 1), ICKs (KIP-RELATED PROTEIN), SIM (SIAMESE)* and *DPP (DP* protein). It was found that plants could sustain growth under different potential stressful conditions (e.g., high UV-B irradiation) by endoreduplication, which is a result of one or more rounds of genome replication without mitosis (Scholes and Paige, 2015; Zedek et al., 2020). The upregulation of these cell division inhibitor genes might be related to the high endoreduplication level found in *H. incana* mature leaves under high-light conditions. Additionally, there were two genes involved in plastid division, *PDV2 (PLASTID DIVISION2)* and *ARC5 (ACCUMULATION AND REPLICATION OF CHLOROPLAST 5)* (**Fig. 6F**). PDV1 and PDV2 are known to function together with ARC5 in the chloroplast to mediate chloroplast division (Miyagishima et al., 2006). Among genes related to leaf and vascular system development (**Fig. 6G**), *PIN5 (PIN-FORMED 5*), several *PILS (PIN-LIKES) 3, 5* and *6, APL (ALTERED PHLOEM DEVELOPMENT)* and *CCI1 (CLAVATA COMPLEX INTERACTOR 1)* were upregulated in high-light samples. PIN5 and PILS were shown to be involved in auxin transport pathways controlling vein patterning (Mravec et al., 2009; Sawchuk et al., 2013) and auxin signaling during environmental stimuli-induced growth adaptation (Waidmann et al., 2023). APL is a transcription factor that regulates vascular tissue development in Arabidopsis (Bonke et al., 2003), while CCI1 is involved in the WUSCHELL/CLAVATA signaling pathway functioning in shoot meristem development (Gish et al., 2013).

Overall, our gene expression analysis supports the key physiological, biochemical and anatomical adaptive changes observed in the *H. incana* leaf tissues in response to high light. As the transcriptome data were derived from one time-point of the whole canopy (i.e., an averaged sample of different developmental stages), our analysis might have missed important genes that were expressed in a more tissue or cell type-specific or developmental stage-specific manner (i.e., transcription factor genes, hence diluted in a bulk sample). In this case, transcriptomes of dissected tissues or single-cells or time-series might be needed to identify key players in these processes. Additionally, our analysis was based solely on transcriptional evidences that support the observed phenotypic changes, further proteomic experiments might need to be conducted to validate this and also identify other genes involving in these changes but were not detected in the transcriptomic analysis. Finally, based on the evidence from comparative phenotypic, genomic and transcriptomic analyses, we propose a model of key processes related to physio-biochemical-anatomical changes in the leaf of *H. incana* in response to high light that might have facilitated its high-photosynthesis traits (**Fig. 6H**).

## CONCLUSIONS

In summary, we successfully reconstructed an improved chromosome-level genome assembly of *H. incana* (Nijmegen accession, v2.0) based on a combination of ONT sequencing and Hi-C technologies. The improved *H. incana* genome assembly and annotation enabled us to elucidate the WGT history of the *H. incana* genome from the common *Brassica* tPCK ancestor, and genome evolution of this species in relation to other species within the Brassiceae tribe. We were able to assign the triplicated ancestral genomic blocks within the *H. incana* genome into three sub-genomes, with the LF sub-genome showing dominance in gene retention as well as gene expression over the MF_1_/MF_2_ sub-genomes. This sub-genome expression divergence among WGT retained gene copies is likely due to the neo-/sub-functionalization processes. The *H. incana* genome appears to be similar to the *R. sativus* genome, and displays intermediate characteristics of *Brassica* A/C and B genome types. This result might explain the discrepancy observed in the published studies regarding their phylogenetic placement in relation to the “Triangle of U” species. Using the information obtained from comparative physio-biochemical and anatomical studies as a guide, we illustrated the expression changes of the associated gene families which likely facilitate the high-photosynthesis traits under high light in *H. incana*. Overall, the improved genome assembly, annotation and results presented in this work will be a valuable resource for future research to unravel the genetic basis of this exceptional species in terms of light-use efficiency and improvement in photosynthesis for enhanced agricultural production.

## MATERIALS AND METHODS

### Plant materials

Plant materials used in this study were derived from *H. incana* reference line “Nijmegen” which was inbred for more than six rounds by hand-pollination (i.e., F_6_) as previously described by Garassino et al. (2022). For whole-genome and transcriptome sequencing using Nanopore ONT technologies, seeds derived from this line were used. The line was then inbred for another two rounds (i.e., F_8_) in a greenhouse at Wageningen University, the Netherlands, and used for Hi-C sequencing.

### Whole-genome sequencing of the *Hirschfeldia incana* genome

For Nanopore ONT sequencing, 750 mg leaf tissues was collected and used for high-molecular-weight (HMW) genomic DNA extraction according to the LeafGo method (Driguez et al., 2021). An aliquot of 1.2-1.5 ug of HMW gDNA was used for library preparation following the Oxford Nanopore SQK-LSK114 kit. We generated three different libraries without size selection, and with size selection (two bins of >25 kb and >40 kb thresholds). Each library sequenced using a MinION platform on an R10.4.1 flow cell and combined for downstream analyses. Base calling was performed using Dorado Duplex v0.3.4 (https://github.com/nanoporetech/dorado) with default parameters and *“--min-qscore 10*” and the “*dna_r10.4.1_e8.2_400bps_sup@v4.2.0*” model to utilize duplex reads and obtain higher read quality compared to the simplex base-calling method.

For Hi-C sequencing based on the Dovetail Omni-C library protocol, chromatin was first fixed with formaldehyde in the nucleus and then extracted. Extracted chromatin was then digested with *DNAse* I, chromatin ends were repaired and subsequently ligated to a biotinylated bridge adapter followed by proximity ligation of adapter containing ends. After that, crosslinks were reversed, and the DNA was purified. The purified DNA underwent treatment to eliminate any biotin that was not internally bound to ligated fragments. Sequencing libraries were prepared using NEBNext Ultra enzymes and Illumina-compatible adapters. Fragments containing biotin were isolated using streptavidin beads prior PCR enrichment to achieve the final library. The final library was subsequently sequenced on an Illumina HiSeqX platform to produce ∼30× genome coverage. All steps were performed by Dovetail Genomics (Scotts Valley, CA, USA).

### Chromosome-scale assembly of the *Hirschfeldia incana* genome

The genome size of *H. incana* was re-estimated using a total 102 million Illumina WGS data from Garassino et al. (2022) and GenomeScope v2.0 (Vurture et al., 2017). The *k-mer* distribution was generated by KMC v3 (Kokot et al., 2017) with default settings (*k-mer* = 21) and the “*-cx1000000*” option to account for the high-frequency *k-mers* derived from repeat content in the genome. The updated genome size estimation was 421 Mb (**Supplemental Fig. S2**), higher than the previously reported size of 325 Mb by Garassino et al. (2022) which was found using GenomeScope v1.0.

The *H. incana* draft genome v1.0 (Garassino et al. (2022) was used as a starting point for assembly. ONT trimmed reads of a minimum of 10 kb were used for genome scaffolding using ntLink v1.3.9 (Coombe et al., 2023) with the options “*gap-fill, k=32, w=500*”. The ONT-derived assembly was polished for two rounds by RACON v1.4.3 (Vaser et al., 2017) using ONT data from this study and Illumina data from Garassino et al. (2022). The resulting ONT genome assembly is labelled v1.5. Assembly v1.5 and Hi-C reads were used as input data for chromosome-level scaffolding employing Juicer v1.6 (Durand et al., 2016a) with default settings and *“-s none”* for *DNAse*-treated data. The output file “*merged_nodups*” was subjected to the 3D-DNA pipeline v201008 (https://github.com/theaidenlab/3d-dna) with options *“-i 100000 --sort-output”* to obtain the final genome assembly (v2.0). The Hi-C contact map was visualized and reviewed by Juicebox v2.20.00 (Durand et al., 2016b). Genome quality and completeness were analyzed by QUAST v5.2.0 (Gurevich et al., 2013) and BUSCO v5.4.7 (Simao et al., 2015) based on the plant-specific Embryophyta odb10 dataset which includes 1,614 single-copy orthologs. Mapping back rates were obtained by mapping a total 102 million WGS read data from Garassino et al. (2022) using bowtie2 v2.5.1 (Langmead and Salzberg, 2012) with parameters “*--very-sensitive --no-unal -k 20*”. Genome circular plots were drawn using Circos v0.69-9 (Krzywinski et al., 2009).

### Transcriptome sequencing, transcript assembly and other data acquisition

To aid genome annotation, we used the ONT sequencing technology to generate long-read transcriptome data for six samples of *H. incana* leaves collected from two conditions, low (200 µmol m^-2^ s^-1^) and high-light (1,800 µmol m^-2^ s^-1^) six weeks after planting. Leaf samples were collected from the 3^rd^ -5^th^ compound leaves counting from the top of each plant (referred to as functional leaves) and snap-frozen in liquid N_2_ and stored at -80°C until further processing. RNA isolation was performed following the TRIzol™ Reagent RNA isolation protocol (Thermo Fisher Scientific, Waltham, MA, USA) combined with the RQ1 RNase-Free DNase Protocol (Promega, Madison, WA, USA) using ∼100 mg ground tissue. The Nanopore sequencing library preparation was conducted following the PCR-cDNA Barcoding Kit (SQK-PCB111.24) protocol. Sequencing was performed on the MinION Mk1B platform using flow cells (FLO-MIN106D) and was base-called with Dorado simplex v0.3.4 (https://github.com/nanoporetech/dorado). Chopper v0.8.0 (https://github.com/wdecoster/chopper) was used for trimming the raw reads, which then were used as direct evidence in our genome annotation pipeline. See **Supplemental Fig. S23** for data quality assessment.

We further obtained the *H. incana* leaf transcriptome data from Mabry et al. (2020) and Garassino et al. (2022) and whole-canopy data from Garassino et al. (2023). These Illumina short-reads and their assembled transcripts were utilized during genome annotation and annotation quality check. Transcriptomes were assembled using the Trinity pipeline v2.15.0 (Haas et al., 2013) with default settings. Additionally, the previously assembled transcripts for *H. incana* samples representing above and underground tissues obtained from Hasnaoui et al. (2022) were also utilized in genome annotation. It is important to note here that the data from Garassino et al. (2022) and Garassino et al. (2023) were from the same *H. incana* Nijmegen accession, while data from Mabry et al. (2020) and Hasnaoui et al. (2022) were derived from different accessions.

### Repeat and gene annotation of the *Hirschfeldia incana* genome

Repeats and transposable elements in the genome were masked with RepeatModeler v2.0.3/RepeatMasker v4.1.2 and RepeatProteinMask (Tarailo-Graovac and Chen, 2009). Firstly, the *ab initio* prediction program RepeatModeler was employed to build a *de novo* repeat library based on the *H. incana* genome. Then, using a custom library that consisted of *de novo* identified repeats, Dfam v3.3 and RepBaseRepeatMaskerEdition-20181026 as database, RepeatMasker was run to find and classify repetitive elements in the genome. The centromere location of each chromosome was identified following the approach used for radish genomes (Jeong et al., 2016; Cho et al., 2022). Briefly, the centromeric tandem repeats (CENTs) from *R. sativus* and *Brassica* genomes were blasted against the *H. incana* genome with an E-value cutoff of 1×10^-10^. The blast hit results were analyzed to identify the likely centromeric regions of the *H. incana* chromosomes.

For *de novo* gene prediction, the PASA pipeline v2.4.1 (Haas et al., 2008) was used to train a model using the available RNA-seq data as direct evidence and the resulting PASA models were then used to train AUGUSTUS v3.1.0 (Stanke and Morgenstern, 2005). Other *de novo* gene predictions were performed using SNAP v20131129 (Korf, 2004), GeneMark v4.72 (Bruna et al., 2020) and GlimmerHMM v3.0.4 (Majoros et al., 2004). Protein evidence used during gene prediction was collected from the UniProtKb/SwissProt curated protein database Release 2024_02 (UniProt, 2015), Viridiplantae dataset from OrthoDB v11 (Kuznetsov et al., 2023) and proteins from related species including *A. thaliana*, *B. rapa*, *B. oleracea*, *B. nigra* and *H. incana* v1.0. We combined the predicted gene models from different programs, transcript evidence and protein evidence to produce consensus gene sets by using EvidenceModeler v2.1.0 (Haas et al., 2008) stand alone and within the Funannotate pipeline v1.8.16 (https://github.com/nextgenusfs/funannotate). This combination step was performed by giving different weights to different predictions, as follows *“--weights augustus:2 pasa:10 snap:1 transcripts:6 proteins:3 GeneMark:1 GlimmerHMM:1*” to reduce spurious gene prediction by *de novo* prediction programs. The BRAKER v3.0.7 (Hoff et al., 2019) and HELIXER v0.3.2 (Holst et al., 2023) annotation pipelines were also tested and compared with the results from EvidenceModeler and Funannotate pipelines to choose the best annotation set. The final annotation was updated by PASA to add untranslated regions (UTR) data and to fix gene models that were not in agreement with the RNA-seq data. Initially, the annotation was done for the ONT-derived genome assembly v1.5, then lifted over to the final Hi-C genome assembly (v2.0) using LIFTOFF v1.6.3 (Shumate and Salzberg, 2021). The BUSCO assessment based on 1,614 Embryophyta single-copy orthologs and OMArk v0.3.0 based on 17,999 conserved orthologs of the Brassicaceae family (Nevers et al., 2022) were used to analyze and compare the annotations.

### Gene functional annotation

The *H. incana* predicted proteins were blasted against the Swiss-Prot release 2022_04 (O’Donovan et al., 2002) and TrEMBL release 2022_01 (O’Donovan et al., 2002) using Diamond BLASTP v2.0.14 (Buchfink et al., 2021) with the following settings “*-e 1e-5 -k 1*”. To predict protein function (both protein domains and associated GO terms), we used InterProScan-5.66-98.0 (Zdobnov and Apweiler, 2001) to blast the *H. incana* proteins against several databases with the options*“-goterms*”. We utilized all 17 databases provided with InterProScan to maximize the annotation. KEGG mapping was done using BlastKOALA v2.2 (Kanehisa et al., 2016) with “*plants*” as taxonomy group and searched against the “*family_eukaryotes*” KEGG gene databases.

### Orthogroup classification

To infer the orthology of *H. incana* and other Brassicaceae genomes, primary (longest variant) protein sequences were used for orthogroup clustering by OrthoFinder v2.5.5 (Emms and Kelly, 2019) with default settings and the “-*M msa*” option to infer maximum likelihood (ML) gene trees from multiple sequence alignment. Additionally, OrthoFinder was used for identification of single-copy orthologs across genomes and reconstruction of a species tree based on the identified single-copy orthologs. This species tree was used to compare with those generated from IQ-TREE v2.0.6 (Minh et al., 2020) and ASTRAL v5.7.1 (Zhang et al., 2018) to infer the relationship between *H. incana* and other species within the Brassiceae used in this study (details in “**Nuclear phylogenetic analyses**” section.

### Genome synteny and duplication analyses

Macro- and micro-synteny of the genomes of *H. incana* and other Brassicaceae species was analyzed by SynMap (Lyons et al., 2008), SynFind (Tang et al., 2015) on the CoGe v7 (Castillo et al., 2018) and MCscan v0.8 (Tang et al., 2008) python version (https://github.com/tanghaibao/jcvi/wiki/MCscan-(Python-version)). Additionally, MCScanX (accessed Dec 2023) (Wang et al., 2012) and DupGen_finder (accessed Dec 2023) (Qiao et al., 2019) were employed for various analyses. Modes of duplicated gene copies were analyzed by DupGen_finder with default parameters using the *A. thaliana* genome as reference. For each Brassiceae genome, gene duplications were classified into WTD/WGD, tandem, proximal, transposed, and dispersed duplicates.

### Estimation of *Ks* ratios of WGT/WGD duplicated gene pairs

The *Ks* (the ratio of number of substitutions per synonymous site) values were computed for WGT/WGD gene pairs identified by DupGen_finder using *KaKs*_Calculator v2.0 (Wang et al., 2010) following the pipeline in Qiao et al. (2019). This employed MAFFT v7.480 (Katoh et al., 2002) and PAL2NAL v14 (Suyama et al., 2006) and the γ-MYN method (Wang et al., 2009a). To identify the *Ks* peaks corresponding to the recent WGT/WGD events in the *H. incana* genome, we fitted the *Ks* distribution using a Gaussian mixture model (GMM) as described in Qiao et al. (2019). To infer species divergence, *Ks* values between syntenic gene pairs were also calculated by CodeML (Yang, 2007) in SynMap running on the CoGe v7 (https://genomevolution.org/coge/).

### Ancestral genomic blocks and karyotype evolution of the *Hirschfeldia incana* genome

The ancestral tPCK genomic blocks (Schranz et al., 2006; Lysak et al., 2016) were used to analyze the *H. incana* genome structure. We utilized the updated genomic blocks for the *Brassica* genomes in He et al. (2021) to determine the intervals and boundaries of the 26 ancestral genomic blocks in the *H. incana* genomes. The triplicated *H. incana* genomic blocks were then classified into three sub-genomes, LF, MF_1_ and MF_2_ sub-genomes based on gene retention rate compared to the ancestral blocks. Briefly, we aligned the *H. incana* genome to the 26 tPCK ancestral genomic blocks (as target/reference) using SynMap. This analysis was ran together with the program FractBias (Joyce et al., 2017) on the CoGe v7 using a window size of 100 genes. The fractionation bias rate was calculated for syntenic genes in the target genome. Syntenic depth was set to 1:3 based on the ploidy level between the ancestral genomes and *H. incana*. Additionally, to elucidate genome rearrangement among *H. incana*, *R. sativus, S. arvensis* and three *Brassica* A/C/B genomes, we employed the IAGS pipeline (accessed Feb 2024) (Gao et al., 2022) to reconstruct their common ancestral genome using the orthologous results from OrthoFinder and the non-overlapping syntenic blocks detected by Drimm-Synteny (accessed Feb 2024) (Pham and Pevzner, 2010).

### Nuclear phylogenetic analyses

To reconstruct the species tree, single-copy orthologs were identified by OrthoFinder v2.5.5 across selected genomes as described earlier. This used MAFFT v7.480 (Katoh et al., 2002) for sequence alignment and FastTree v2 (Price et al., 2009) for the phylogenetic tree inference. For IQ-TREE analysis, coding or protein sequences were aligned by MAFFT with the option the option “*G-INS-i*”, then poorly aligned regions were trimmed by trimAL v1.4.rev22 (Capella-Gutierrez et al., 2009) with the option “*-automated1*”. The alignment files then were subjected to IQ-TREE v2.2.0 (Trifinopoulos et al., 2016) with default settings (1,000 bootstrap iterations).

For phylogenetic incongruency analyses, coding-sequences from the 5,765 “strict single-copy genes” identified among six species (*A. thaliana, B. rapa, B. oleracea, B. nigra, R. sativus* and *H. incana*) using OrthoFinder were aligned using MACSE v2.06 (Ranwez et al., 2018). Gene trees were reconstructed using RAxML-NG v1.1.0 (Kozlov et al., 2019) with substitution model GTR+G and 1000 bootstrap replicates while setting *A. thaliana* as the outgroup. A species tree was reconstructed with ASTRAL v5.7.8 (Zhang et al., 2018) based on all gene trees. The tree topology was further used as input for DensiTree v3.0.3 (Bouckaert, 2010) and PhyParts v0.0.1 (Smith et al., 2015) to assess conflict among the gene trees. For each bipartition, the software assesses the number of gene trees that support the main topology, the most common alternative topology, all other topologies, and the number of gene trees that are not informative for the respective bipartition. Here, we also employed three support levels (no threshold, bootstrap support < 50%, < 85%) to count as uninformative. PhyPartsPieCharts (https://github.com/mossmatters/phyloscripts/tree/master/phypartspiecharts, accessed Feb 2024) was used for visualization.

Genes retained in all three sub-genomes (triads) across all species were obtained from SynMap analysis using the ancestral block as reference against the target genomes. We focused our analysis on 90 genes from shared ancestral block F. Codon-aware alignments were created using MACSE v2.06, and gene trees were reconstructed using RAxML-NG v1.1.0 as described above. We then reconstructed two different types of trees. First, a species tree was reconstructed using ASTRAL-pro v1.15.1.3 (Zhang et al., 2020), which allows for multi-copy genes; here the three gene copies in each Brassiceae species were considered paralogs. Second, a sub-genome tree was reconstructed using ASTRAL v5.7.8, in which the three copies in each species were assigned to their respective sub-genomes LF, MF_1_ and MF_2_. The final consensus trees were visualized by FigTree v1.4.3 (http://evomics.org/resources/software/molecular-evolution-software/figtree/).

### Endoreduplication analysis by flow cytometry

Endoreduplication (endopolyploidy) analysis was performed using leaf samples from three species, *B. rapa* (R-o-18 accession), *B. nigra* (DG1 accession) and *H. incana* (Nijmegen accession) by flow cytometry (Plant Cytometry, Didam, The Netherlands). Leaf samples were collected from plants grown under low (200 µmol m^-2^ s^-1^) and high-light (1,800 µmol m^-2^ s^-1^) at 30 days after sowing. Three leaf developmental stages (very young, young and mature) as marked in **Fig. 6A** were used.

### Gene expression analysis

For analysis of gene expression in leaf tissues, we utilized whole canopy transcriptome data reported in Garassino et al. (2023) from two contrasting light conditions, low (200 µmol m^-2^ s^-1^) and high-light (1,800 µmol m^-2^ s^-1^). RNA-seq read quality before and after trimming was assessed by FastQC v0.11.9 (http://www.bioinformatics.babraham.ac.uk/projects/fastqc). Adapter sequences and low-quality reads were trimmed using Trimmomatic v0.39 (Bolger et al., 2014) and the following parameters: “ILLUMINACLIP: 2:20:10 SLIDINGWINDOW:4:15 LEADING:5 TRAILING:5 MINLEN:50.” To estimate transcript abundance, cleaned reads were mapped onto *H. incana* gene models using Bowtie2 v2.4.5 (Langmead and Salzberg, 2012) with default settings. The mapping BAM files were sorted by SAMTOOLS-1.19.2 (Li et al., 2009) and subjected to RSEM v1.3.3 (Li and Dewey, 2011) for quantification of transcript abundance, normalized as transcripts per million transcripts (TPM). Differentially expressed genes were identified using DESeq2 package (Love et al., 2014) with an FDR corrected *p*-value ≤0.05 and |fold-change| ≥2. Additionally, we employed Mercator v4.6 (Lohse et al., 2014) to functionally annotate and categorize the identified differentially expressed genes.

### Other quantification and statistical analyses

Venn diagrams were generated using the online tools (http://bioinformatics.psb.ugent.be/webtools/Venn) and InteractiVenn (Heberle et al., 2015). GO term and KEGG pathway enrichment of gene sets were performed using the DAVID bioinformatics resources v2023q4 (Huang et al., 2009) and ShinyGO v0.80 (Ge et al., 2020). All analyses in the Linux environment were performed on local servers running Ubuntu 16.04.6 LTS hosted by the Biosystematics Group at Wageningen University, the Netherlands. All statistical analyses, unless otherwise stated, were performed in Microsoft Excel v18.2311.1071.0 and R v4.0.2 with RStudio v2022.07.2-576 (https://www.rstudio.com).

### Data availability and accession numbers

Data supporting the findings in this work are available here and in Supplemental Data files. The raw WGS and transcriptome data from Illumina and Nanopore platforms used for this study have been deposited at NCBI SRA database under BioProject PRJNA1045848. The final assembly and annotation (v2.0) of the *H. incana* genome can be downloaded from Figshare at https://doi.org/10.6084/m9.figshare.25574799. The final genome sequence has also been deposited at DDBJ/ENA/GenBank under the accession JBAWST000000000. The version described in this paper is version JBAWST010000000. Sequence alignment files and machine-readable phylogenetic trees related to phylogenetic analyses presented in this work are available via Figshare at https://doi.org/10.6084/m9.figshare.25574811.

The *A. thaliana* araport11 genome data were downloaded from Phytozome 13 (https://phytozome-next.jgi.doe.gov/info/Athaliana_Araport11). The *T. parvula* genome v8.4 were downloaded from http://thellungiella.org. The *B. rapa* Chiifu genome (v4.0 and v4.1), *B. oleracea* JZS genome v2.0 and Broccoli HDEM data were downloaded from the Brassicaceae Database (http://brassicadb.cn/#/). The *B. nigra* NI100 v2.0 and C2 was downloaded from the Crucifer Genome Initiative https://cruciferseq.ca. The *B. rapa* chinensis Pakchoi genome was obtained from https://doi.org/10.6084/m9.figshare.19589524.v2. The *B. oleracea* genomes for accessions Cauliflower Korso and Kale were respectively downloaded from the Brassica oleracea Genome Database http://www.bogdb.com/genome/cauliflower and Ensembl Genomes https://ftp.ensemblgenomes.ebi.ac.uk/pub/plants/release-58/, respectively. The *R. sativus* genomes for accessions WK10039 (v1.0 and v2.0) and NAULB were respectively obtained from https://doi.org/10.6084/m9.figshare.21671201.v2, https://www.ncbi.nlm.nih.gov/datasets/genome/GCF_000801105.2/, https://download.cncb.ac.cn/gwh/Plants/Raphanus_sativus_NAU-LB_GWHCBIT00000000. The genomes of *B. nigra* Giebra and *S. arvensis* were obtained from https://doi.org/10.6084/m9.figshare.21442935.v1. The genomes of *B. tournefortii*, *Sinapis* sp. HIR3, *H. incana* HIR1 were obtained from https://doi.org/10.6084/m9.figshare.21671201.v2.

## ACKNOWLEDGEMENTS

The authors would like to thank Sandra Smit, Francesco Garassino, Harm Nijveen and Rik Janssen for generous help and suggestions on genome annotation strategies; Xinyou Yin for providing helpful comments and suggestions on our manuscript; Freek Bakker, Klaas Bouwmeester and Tao Feng for interesting suggestions and discussions on phylogenetic and gene function analyses; Felix Akens for kind help with plant growth experiment; and Tianpeng Wang for his kind help and discussion of the ancestral genomic block analysis. We acknowledge the members of the Biosystematics Group at Wageningen University for their helpful and insightful comments and discussion during our regular meetings.

## AUTHOR CONTRIBUTIONS

MES and MGMA conceived the project; MES supervised the data analysis and manuscript preparation; NVH and MES coordinated the genome sequencing and assembly, annotated the genome and analyzed the data; NVH prepared the first draft of the paper and figures with the inputs from other authors; FFMB, NVH, SBL and MGMA contributed to whole-genome Nanopore sequencing; RL, PV, NVH and LC contributed to transcriptome Nanopore sequencing; NW and NVH contributed to nuclear phylogenetic analysis; FW, TW and NVH generated the 7^th^ and 8^th^ generation of *H. incana* Nijmegen accession used for Hi-C sequencing; LV, JH and HvA contributed to chloroplast analysis; MR and PCS contributed to microCT imaging data analysis of additional chloroplast density experiment; SBL provided samples for endoreduplication analysis; DdR and SMD participated in various meetings and discussions during genome sequencing and assembly of this project. All authors read, edited, and approved the final manuscript.

## SUPPLEMENTAL DATA

**Supplemental Figure file contains the following 23 figures:**

**- Figure S1.** Summary of the Nanopore ONT whole-genome sequencing data (combined) (upper panel) and Hi-C data base quality (lower panel).

**- Figure S2.** Size estimation of the *H. incana* genome using GenomeScope v2.0.

**- Figure S3.** Comparison among the assemblies from the Brassiceae.

**- Figure S4.** Comparison of different annotation approaches of the *H. incana* genome v1.5.

**- Figure S5.** Comparison of annotated proteomes and coding sequences (CDS) from selected Brassicaceae genomes..

**- Figure S6.** Dotplot showing synteny between *H. incana* v1.0 and v2.0, analyzed by SynMap program.

**- Figure S7.** OMArk proteome assessment of selected Brassicaceae genomes.

**- Figure S8.** Macro-synteny between *A. thaliana* and *H. incana*.

**- Figure S9.** Intra-genomic syntenic (self-comparison) dotplot the *H. incana* genome v2.0.

**- Figure S10.** Macro-synteny between *B. rapa* and *H. incana*.

**- Figure S11.** Gene duplication modes of the *H. incana* genome v2.0 compared with other Brassicaceae genomes.

**- Figure S12.** Syntenic relationship between the *H. incana* genome v2.0 and the Brassica tPCK ancestral genomic blocks (Bra-ACK).

**- Figure S13.** Additional plot showing syntenic relationship between the *H. incana* genome v2.0 and the Brassica tPCK ancestral genomic blocks (Bra-ACK).

**- Figure S14.** Evolutionary history and genome rearrangement estimation of the six Brassiceae genomes.

**- Figure S15.** Genome syntenic dotplot and ribbons between *R. sativus* and *H. incana* v2.0.

**- Figure S16.** PhyParts tree topology quantification.

**- Figure S17.** The identification of the tPCK ancestral genomic block F in the six Brassiceae genomes.

**- Figure S18.** Species tree reconstructed from 90 triad gene sets from three sub-genomes of each Brassiceae species.

**- Figure S19.** GO biological process (upper panel) and KEGG pathway (lower panel) enrichment analyses of all well-retained 2,103 triad genes.

**- Figure S20.** GO biological process (upper panel) and KEGG pathway (lower panel) enrichment analyses of two-copy retained 6,457 dyad genes.

**- Figure S21.** A summary of *H. incana* whole canopy transcriptome data.

**- Figure S22.** GO biological process enrichment analyses of 3,527, 1,932 and 1,310 single-copy genes found on the LF, MF1 and MF2 sub-genomes.

**- Figure S23.** Nanopore RNA sequencing data statistics.

**Supplemental Table file contains the following 18 tables:**

**- Table S1.** Sequencing data information for the assembly of the *H. incana* genome v2.0.

**- Table S2.** Comparison of assemblies by QUAST.

**- Table S3.** Mapping rates of Illumina data onto the three assemblies.

**- Table S4.** BUSCO completeness of the three *H. incana* assemblies.

**- Table S5.** Repeat masking of the *H. incana* genome v2.0.

**- Table S6.** BUSCO completeness assessment of the final *H. incana* genome annotation v2.0 (lifted from v1.5).

**- Table S7.** Summary of functional annotation of the *H. incana* gene set.

**- Table S8.** Assessment of selected proteomes from Brassicaceae by OMArk.

**- Table S9.** OrthoFinder ortholog groups clustering results.

**- Table S10.** Intervals and boundaries of the 26 ancestral genomic blocks in the *H. incana* genome.

**- Table S11.** A list of 5,675 single-copy genes identified among the selected genomes.

**- Table S12.** Sub-genome assignment of 5,675 single-copy genes identified the *H. incana* genome.

**- Table S13.** A set of 90 syntenic orthologous gene groups identified across sub-genomes of six selected Brassiceae species.

**- Table S14.** KEGG pathway and GO term enrichment of upregulated genes in high-light compared to low-light condition in *H. incana*.

**- Table S15.** Sub-genome biased gene expression analysis (pairwise comparisons).

**- Table S16.** Sub-genome biased gene expression analysis (all three sub-genome comparison).

**- Table S17.** Endoreduplication analysis of *B. rapa*, *B. nigra* and *H. incana* under low-light and high-light conditions.

**- Table S18.** A list of 2,012 up- and 2,126 downregulated genes identified in whole canopy transcriptome data.

## REFERENCES

Arias, T., and Pires, J.C. (2018). A fully resolved chloroplast phylogeny of the *Brassica* crops and wild relatives (Brassicaceae: Brassiceae): Novel clades and potential taxonomic implications. Taxon 61, 980–988.

Ashburner, M., Ball, C.A., Blake, J.A., Botstein, D., Butler, H., Cherry, J.M., Davis, A.P., Dolinski, K., Dwight, S.S., Eppig, J.T., Harris, M.A., Hill, D.P., Issel-Tarver, L., Kasarskis, A., Lewis, S., Matese, J.C., Richardson, J.E., Ringwald, M., Rubin, G.M., and Sherlock, G. (2000). Gene ontology: tool for the unification of biology. The Gene Ontology Consortium. Nat. Genet. 25, 25–29.

Auguy, F., Fahr, M., Moulin, P., Brugel, A., Laplaze, L., Mzibri, M.E., Filali-Maltouf, A., Doumas, P., and Smouni, A. (2013). Lead tolerance and accumulation in *Hirschfeldia incana*, a Mediterranean Brassicaceae from metalliferous mine spoils. PLoS One 8, e61932.

Barker, M.S., Li, Z., Kidder, T.I., Reardon, C.R., Lai, Z., Oliveira, L.O., Scascitelli, M., and Rieseberg, L.H. (2016). Most Compositae (Asteraceae) are descendants of a paleohexaploid and all share a paleotetraploid ancestor with the Calyceraceae. Am. J. Bot. 103, 1203–1211.

Belser, C., Istace, B., Denis, E., Dubarry, M., Baurens, F.C., Falentin, C., Genete, M., Berrabah, W., Chevre, A.M., Delourme, R., Deniot, G., Denoeud, F., Duffe, P., Engelen, S., Lemainque, A., Manzanares-Dauleux, M., Martin, G., Morice, J., Noel, B., Vekemans, X., D’Hont, A., Rousseau-Gueutin, M., Barbe, V., Cruaud, C., Wincker, P., and Aury, J.M. (2018). Chromosome-scale assemblies of plant genomes using nanopore long reads and optical maps. Nat Plants 4, 879–887.

Bolger, A.M., Lohse, M., and Usadel, B. (2014). Trimmomatic: a flexible trimmer for Illumina sequence data. Bioinformatics 30, 2114–2120.

Bonke, M., Thitamadee, S., Mahonen, A.P., Hauser, M.T., and Helariutta, Y. (2003). APL regulates vascular tissue identity in Arabidopsis. Nature 426, 181–186.

Bouckaert, R.R. (2010). DensiTree: making sense of sets of phylogenetic trees. Bioinformatics 26, 1372–1373.

Brown, N.J., Parsley, K., and Hibberd, J.M. (2005). The future of C4 research--maize, Flaveria or Cleome? Trends Plant Sci. 10, 215–221.

Bruna, T., Lomsadze, A., and Borodovsky, M. (2020). GeneMark-EP+: eukaryotic gene prediction with self-training in the space of genes and proteins. NAR Genom Bioinform 2, lqaa026.

Buchfink, B., Reuter, K., and Drost, H.G. (2021). Sensitive protein alignments at tree-of-life scale using DIAMOND. Nat. Methods 18, 366–368.

Cai, X., Wu, J., Liang, J., Lin, R., Zhang, K., Cheng, F., and Wang, X. (2020). Improved *Brassica oleracea* JZS assembly reveals significant changing of LTR-RT dynamics in different morphotypes. Theor. Appl. Genet. 133, 3187–3199.

Canvin, D.T., Berry, J.A., Badger, M.R., Fock, H., and Osmond, C.B. (1980). Oxygen exchange in leaves in the light. Plant Physiol. 66, 302–307.

Capella-Gutierrez, S., Silla-Martinez, J.M., and Gabaldon, T. (2009). trimAl: a tool for automated alignment trimming in large-scale phylogenetic analyses. Bioinformatics 25, 1972–1973.

Castillo, A.I., Nelson, A.D.L., Haug-Baltzell, A.K., and Lyons, E. (2018). A tutorial of diverse genome analysis tools found in the CoGe web-platform using *Plasmodium* spp. as a model. Database 2018, bay030.

Chen, K.M., Holmstrom, M., Raksajit, W., Suorsa, M., Piippo, M., and Aro, E.M. (2010). Small chloroplast-targeted DnaJ proteins are involved in optimization of photosynthetic reactions in *Arabidopsis thaliana*. BMC Plant Biol. 10, 43.

Cheng, F., Wu, J., and Wang, X. (2014). Genome triplication drove the diversification of *Brassica* plants. Hortic Res 1, 14024.

Cheng, F., Wu, J., Fang, L., Sun, S., Liu, B., Lin, K., Bonnema, G., and Wang, X. (2012). Biased gene fractionation and dominant gene expression among the subgenomes of *Brassica rapa*. PLoS One 7, e36442.

Cheng, F., Sun, R., Hou, X., Zheng, H., Zhang, F., Zhang, Y., Liu, B., Liang, J., Zhuang, M., Liu, Y., Liu, D., Wang, X., Li, P., Liu, Y., Lin, K., Bucher, J., Zhang, N., Wang, Y., Wang, H., Deng, J., Liao, Y., Wei, K., Zhang, X., Fu, L., Hu, Y., Liu, J., Cai, C., Zhang, S., Zhang, S., Li, F., Zhang, H., Zhang, J., Guo, N., Liu, Z., Liu, J., Sun, C., Ma, Y., Zhang, H., Cui, Y., Freeling, M.R., Borm, T., Bonnema, G., Wu, J., and Wang, X. (2016). Subgenome parallel selection is associated with morphotype diversification and convergent crop domestication in *Brassica rapa* and *Brassica oleracea*. Nat. Genet. 48, 1218–1224.

Cho, A., Jang, H., Baek, S., Kim, M.J., Yim, B., Huh, S., Kwon, S.H., Yu, H.J., and Mun, J.H. (2022). An improved *Raphanus sativus* cv. WK10039 genome localizes centromeres, uncovers variation of DNA methylation and resolves arrangement of the ancestral Brassica genome blocks in radish chromosomes. Theor. Appl. Genet. 135, 1731–1750.

Coombe, L., Warren, R.L., Wong, J., Nikolic, V., and Birol, I. (2023). ntLink: A Toolkit for *De Novo* Genome Assembly Scaffolding and Mapping Using Long Reads. Curr Protoc 3, e733.

Davletova, S., Schlauch, K., Coutu, J., and Mittler, R. (2005). The zinc-finger protein Zat12 plays a central role in reactive oxygen and abiotic stress signaling in Arabidopsis. Plant Physiol. 139, 847–856.

De Smet, R., Adams, K.L., Vandepoele, K., Van Montagu, M.C., Maere, S., and Van de Peer, Y. (2013). Convergent gene loss following gene and genome duplications creates single-copy families in flowering plants. Proc. Natl. Acad. Sci. U. S. A. 110, 2898–2903.

Driguez, P., Bougouffa, S., Carty, K., Putra, A., Jabbari, K., Reddy, M., Soppe, R., Cheung, M.S., Fukasawa, Y., and Ermini, L. (2021). LeafGo: Leaf to Genome, a quick workflow to produce high-quality de novo plant genomes using long-read sequencing technology. Genome Biol. 22, 256.

Durand, N.C., Shamim, M.S., Machol, I., Rao, S.S., Huntley, M.H., Lander, E.S., and Aiden, E.L. (2016a). Juicer Provides a One-Click System for Analyzing Loop-Resolution Hi-C Experiments. Cell Syst 3, 95–98.

Durand, N.C., Robinson, J.T., Shamim, M.S., Machol, I., Mesirov, J.P., Lander, E.S., and Aiden, E.L. (2016b). Juicebox Provides a Visualization System for Hi-C Contact Maps with Unlimited Zoom. Cell Syst 3, 99–101.

Edger, P.P., Hall, J.C., Harkess, A., Tang, M., Coombs, J., Mohammadin, S., Schranz, M.E., Xiong, Z., Leebens-Mack, J., Meyers, B.C., Sytsma, K.J., Koch, M.A., Al-Shehbaz, I.A., and Pires, J.C. (2018). Brassicales phylogeny inferred from 72 plastid genes: A reanalysis of the phylogenetic localization of two paleopolyploid events and origin of novel chemical defenses. Am. J. Bot. 105, 463–469.

Edwards, E.J. (2019). Evolutionary trajectories, accessibility and other metaphors: the case of C4 and CAM photosynthesis. New Phytol. 223, 1742–1755.

Emms, D.M., and Kelly, S. (2019). OrthoFinder: phylogenetic orthology inference for comparative genomics. Genome Biol. 20, 238.

Feng, X., Chen, Q., Wu, W., Wang, J., Li, G., Xu, S., Shao, S., Liu, M., Zhong, C., Wu, C.I., Shi, S., and He, Z. (2024). Genomic evidence for rediploidization and adaptive evolution following the whole-genome triplication. Nat Commun 15, 1635.

Forsythe, E.S., Nelson, A.D.L., and Beilstein, M.A. (2020). Biased Gene Retention in the Face of Introgression Obscures Species Relationships. Genome Biol. Evol. 12, 1646–1663.

Gao, S., Yang, X., Sun, J., Zhao, X., Wang, B., and Ye, K. (2022). IAGS: Inferring Ancestor Genome Structure under a Wide Range of Evolutionary Scenarios. Mol Biol Evol 39.

Garassino, F., Luoni, S.B., Cumerlato, T., Marquez, F.R., Harbinson, J., Aarts, M.G.M., Nijveen, H., and Smit, S. (2023). Comparative transcriptomics of *Hirschfeldia incana* and relatives highlights differences in photosynthetic pathways. bioRxiv, 2023.2010.2018.562717.

Garassino, F., Wijfjes, R.Y., Boesten, R., Reyes Marquez, F., Becker, F.F.M., Clapero, V., van den Hatert, I., Holmer, R., Schranz, M.E., Harbinson, J., de Ridder, D., Smit, S., and Aarts, M.G.M. (2022). The genome sequence of *Hirschfeldia incana*, a new Brassicaceae model to improve photosynthetic light-use efficiency. Plant J. 112, 1298–1315.

Ge, S.X., Jung, D., and Yao, R. (2020). ShinyGO: a graphical gene-set enrichment tool for animals and plants. Bioinformatics 36, 2628–2629.

Gish, L.A., Gagne, J.M., Han, L., Deyoung, B.J., and Clark, S.E. (2013). WUSCHEL-responsive At5g65480 interacts with CLAVATA components *in vitro* and in transient expression. PLoS One 8, e66345.

Gowik, U., Brautigam, A., Weber, K.L., Weber, A.P., and Westhoff, P. (2011). Evolution of C4 photosynthesis in the genus Flaveria: how many and which genes does it take to make C4? Plant Cell 23, 2087–2105.

Guerreiro, R., Bonthala, V.S., Schluter, U., Hoang, N.V., Triesch, S., Schranz, M.E., Weber, A.P.M., and Stich, B. (2023). A genomic panel for studying C3-C4 intermediate photosynthesis in the Brassiceae tribe. Plant Cell Environ. 46, 3611–3627.

Guo, N., Wang, S., Gao, L., Liu, Y., Wang, X., Lai, E., Duan, M., Wang, G., Li, J., Yang, M., Zong, M., Han, S., Pei, Y., Borm, T., Sun, H., Miao, L., Liu, D., Yu, F., Zhang, W., Ji, H., Zhu, C., Xu, Y., Bonnema, G., Li, J., Fei, Z., and Liu, F. (2021). Genome sequencing sheds light on the contribution of structural variants to *Brassica oleracea* diversification. BMC Biol. 19, 93.

Gurevich, A., Saveliev, V., Vyahhi, N., and Tesler, G. (2013). QUAST: quality assessment tool for genome assemblies. Bioinformatics 29, 1072–1075.

Haas, B.J., Salzberg, S.L., Zhu, W., Pertea, M., Allen, J.E., Orvis, J., White, O., Buell, C.R., and Wortman, J.R. (2008). Automated eukaryotic gene structure annotation using EVidenceModeler and the Program to Assemble Spliced Alignments. Genome Biol. 9, R7.

Haas, B.J., Papanicolaou, A., Yassour, M., Grabherr, M., Blood, P.D., Bowden, J., Couger, M.B., Eccles, D., Li, B., Lieber, M., MacManes, M.D., Ott, M., Orvis, J., Pochet, N., Strozzi, F., Weeks, N., Westerman, R., William, T., Dewey, C.N., Henschel, R., LeDuc, R.D., Friedman, N., and Regev, A. (2013). *De novo* transcript sequence reconstruction from RNA-seq using the Trinity platform for reference generation and analysis. Nat. Protoc. 8, 1494–1512.

Hao, Y., Mabry, M.E., Edger, P.P., Freeling, M., Zheng, C., Jin, L., VanBuren, R., Colle, M., An, H., Abrahams, R.S., Washburn, J.D., Qi, X., Barry, K., Daum, C., Shu, S., Schmutz, J., Sankoff, D., Barker, M.S., Lyons, E., Pires, J.C., and Conant, G.C. (2021). The contributions from the progenitor genomes of the mesopolyploid Brassiceae are evolutionarily distinct but functionally compatible. Genome Res. 31, 799–810.

Hasnaoui, S.E., Fahr, M., Zouine, M., and Smouni, A. (2022). De Novo Transcriptome Assembly, Gene Annotations, and Characterization of Functional Profiling Reveal Key Genes for Lead Alleviation in the Pb Hyperaccumulator Greek Mustard (*Hirschfeldia incana* L.). Curr. Issues Mol. Biol. 44, 4658–4675.

He, Z., Ji, R., Havlickova, L., Wang, L., Li, Y., Lee, H.T., Song, J., Koh, C., Yang, J., Zhang, M., Parkin, I.A.P., Wang, X., Edwards, D., King, G.J., Zou, J., Liu, K., Snowdon, R.J., Banga, S.S., Machackova, I., and Bancroft, I. (2021). Genome structural evolution in *Brassica* crops. Nat Plants 7, 757–765.

Heberle, H., Meirelles, G.V., da Silva, F.R., Telles, G.P., and Minghim, R. (2015). InteractiVenn: a web-based tool for the analysis of sets through Venn diagrams. BMC Bioinformatics 16, 1–7.

Hendriks, K.P., Kiefer, C., Al-Shehbaz, I.A., Bailey, C.D., Hooft van Huysduynen, A., Nikolov, L.A., Nauheimer, L., Zuntini, A.R., German, D.A., Franzke, A., Koch, M.A., Lysak, M.A., Toro-Nunez, O., Ozudogru, B., Invernon, V.R., Walden, N., Maurin, O., Hay, N.M., Shushkov, P., Mandakova, T., Schranz, M.E., Thulin, M., Windham, M.D., Resetnik, I., Spaniel, S., Ly, E., Pires, J.C., Harkess, A., Neuffer, B., Vogt, R., Brauchler, C., Rainer, H., Janssens, S.B., Schmull, M., Forrest, A., Guggisberg, A., Zmarzty, S., Lepschi, B.J., Scarlett, N., Stauffer, F.W., Schonberger, I., Heenan, P., Baker, W.J., Forest, F., Mummenhoff, K., and Lens, F. (2023). Global Brassicaceae phylogeny based on filtering of 1,000-gene dataset. Curr. Biol. 33, 4052–4068 e4056.

Hoang, N.V., Sogbohossou, E.O.D., Xiong, W., Simpson, C.J.C., Singh, P., Walden, N., van den Bergh, E., Becker, F.F.M., Li, Z., Zhu, X.G., Brautigam, A., Weber, A.P.M., van Haarst, J.C., Schijlen, E., Hendre, P.S., Van Deynze, A., Achigan-Dako, E.G., Hibberd, J.M., and Schranz, M.E. (2023). The *Gynandropsis gynandra* genome provides insights into whole-genome duplications and the evolution of C4 photosynthesis in Cleomaceae. Plant Cell 35, 1334–1359.

Hoff, K.J., Lomsadze, A., Borodovsky, M., and Stanke, M. (2019). Whole-Genome Annotation with BRAKER. Methods Mol. Biol. 1962, 65–95.

Holst, F., Bolger, A., Günther, C., Maß, J., Triesch, S., Kindel, F., Kiel, N., Saadat, N., Ebenhöh, O., Usadel, B., Schwacke, R., Bolger, M., Weber, A.P.M., and Denton, A.K. (2023). Helixer–*de novo* Prediction of Primary Eukaryotic Gene Models Combining Deep Learning and a Hidden Markov Model. bioRxiv, 2023.2002.2006.527280.

Huang, C.H., Sun, R., Hu, Y., Zeng, L., Zhang, N., Cai, L., Zhang, Q., Koch, M.A., Al-Shehbaz, I., Edger, P.P., Pires, J.C., Tan, D.Y., Zhong, Y., and Ma, H. (2016). Resolution of Brassicaceae Phylogeny Using Nuclear Genes Uncovers Nested Radiations and Supports Convergent Morphological Evolution. Mol Biol Evol 33, 394–412.

Huang, D.W., Sherman, B.T., and Lempicki, R.A. (2009). Systematic and integrative analysis of large gene lists using DAVID bioinformatics resources. Nat. Protoc. 4, 44–57.

Hutin, C., Nussaume, L., Moise, N., Moya, I., Kloppstech, K., and Havaux, M. (2003). Early light-induced proteins protect Arabidopsis from photooxidative stress. Proc. Natl. Acad. Sci. U. S. A. 100, 4921–4926.

International Wheat Genome Sequencing, C. (2014). A chromosome-based draft sequence of the hexaploid bread wheat (*Triticum aestivum*) genome. Science 345, 1251788.

Jeong, Y.M., Kim, N., Ahn, B.O., Oh, M., Chung, W.H., Chung, H., Jeong, S., Lim, K.B., Hwang, Y.J., Kim, G.B., Baek, S., Choi, S.B., Hyung, D.J., Lee, S.W., Sohn, S.H., Kwon, S.J., Jin, M., Seol, Y.J., Chae, W.B., Choi, K.J., Park, B.S., Yu, H.J., and Mun, J.H. (2016). Elucidating the triplicated ancestral genome structure of radish based on chromosome-level comparison with the *Brassica* genomes. Theor. Appl. Genet. 129, 1357–1372.

Jiao, Y., Wickett, N.J., Ayyampalayam, S., Chanderbali, A.S., Landherr, L., Ralph, P.E., Tomsho, L.P., Hu, Y., Liang, H., Soltis, P.S., Soltis, D.E., Clifton, S.W., Schlarbaum, S.E., Schuster, S.C., Ma, H., Leebens-Mack, J., and dePamphilis, C.W. (2011). Ancestral polyploidy in seed plants and angiosperms. Nature 473, 97–100.

Johnson, M.G., Pokorny, L., Dodsworth, S., Botigue, L.R., Cowan, R.S., Devault, A., Eiserhardt, W.L., Epitawalage, N., Forest, F., Kim, J.T., Leebens-Mack, J.H., Leitch, I.J., Maurin, O., Soltis, D.E., Soltis, P.S., Wong, G.K., Baker, W.J., and Wickett, N.J. (2019). A Universal Probe Set for Targeted Sequencing of 353 Nuclear Genes from Any Flowering Plant Designed Using k-Medoids Clustering. Syst. Biol. 68, 594–606.

Joyce, B.L., Haug-Baltzell, A., Davey, S., Bomhoff, M., Schnable, J.C., and Lyons, E. (2017). FractBias: a graphical tool for assessing fractionation bias following polyploidy. Bioinformatics 33, 552–554.

Kagale, S., Robinson, S.J., Nixon, J., Xiao, R., Huebert, T., Condie, J., Kessler, D., Clarke, W.E., Edger, P.P., Links, M.G., Sharpe, A.G., and Parkin, I.A. (2014). Polyploid evolution of the Brassicaceae during the Cenozoic era. Plant Cell 26, 2777–2791.

Kanehisa, M., and Goto, S. (2000). KEGG: kyoto encyclopedia of genes and genomes. Nucleic Acids Res. 28, 27–30.

Kanehisa, M., Sato, Y., and Morishima, K. (2016). BlastKOALA and GhostKOALA: KEGG Tools for Functional Characterization of Genome and Metagenome Sequences. J. Mol. Biol. 428, 726–731.

Katoh, K., Misawa, K., Kuma, K., and Miyata, T. (2002). MAFFT: a novel method for rapid multiple sequence alignment based on fast Fourier transform. Nucleic Acids Res. 30, 3059–3066.

Kokot, M., Dlugosz, M., and Deorowicz, S. (2017). KMC 3: counting and manipulating k-mer statistics. Bioinformatics 33, 2759–2761.

Korf, I. (2004). Gene finding in novel genomes. BMC Bioinformatics 5, 59.

Kozlov, A.M., Darriba, D., Flouri, T., Morel, B., and Stamatakis, A. (2019). RAxML-NG: a fast, scalable and user-friendly tool for maximum likelihood phylogenetic inference. Bioinformatics 35, 4453–4455.

Krzywinski, M., Schein, J., Birol, I., Connors, J., Gascoyne, R., Horsman, D., Jones, S.J., and Marra, M.A. (2009). Circos: an information aesthetic for comparative genomics. Genome Res. 19, 1639–1645.

Kuznetsov, D., Tegenfeldt, F., Manni, M., Seppey, M., Berkeley, M., Kriventseva, E.V., and Zdobnov, E.M. (2023). OrthoDB v11: annotation of orthologs in the widest sampling of organismal diversity. Nucleic Acids Res. 51, D445–D451.

Langmead, B., and Salzberg, S.L. (2012). Fast gapped-read alignment with Bowtie 2. Nat. Methods 9, 357–359.

Lawson, T., Kramer, D.M., and Raines, C.A. (2012). Improving yield by exploiting mechanisms underlying natural variation of photosynthesis. Curr. Opin. Biotechnol. 23, 215–220.

Li, B., and Dewey, C.N. (2011). RSEM: accurate transcript quantification from RNA-Seq data with or without a reference genome. BMC Bioinformatics 12, 323.

Li, H., Handsaker, B., Wysoker, A., Fennell, T., Ruan, J., Homer, N., Marth, G., Abecasis, G., Durbin, R., and Genome Project Data Processing, S. (2009). The Sequence Alignment/Map format and SAMtools. Bioinformatics 25, 2078–2079.

Li, W., Challa, G.S., Zhu, H., and Wei, W. (2016). Recurrence of Chromosome Rearrangements and Reuse of DNA Breakpoints in the Evolution of the Triticeae Genomes. G3 (Bethesda) 6, 3837–3847.

Liu, D., Hunt, M., and Tsai, I.J. (2018). Inferring synteny between genome assemblies: a systematic evaluation. BMC Bioinformatics 19, 26.

Liu, S., Liu, Y., Yang, X., Tong, C., Edwards, D., Parkin, I.A., Zhao, M., Ma, J., Yu, J., Huang, S., Wang, X., Wang, J., Lu, K., Fang, Z., Bancroft, I., Yang, T.J., Hu, Q., Wang, X., Yue, Z., Li, H., Yang, L., Wu, J., Zhou, Q., Wang, W., King, G.J., Pires, J.C., Lu, C., Wu, Z., Sampath, P., Wang, Z., Guo, H., Pan, S., Yang, L., Min, J., Zhang, D., Jin, D., Li, W., Belcram, H., Tu, J., Guan, M., Qi, C., Du, D., Li, J., Jiang, L., Batley, J., Sharpe, A.G., Park, B.S., Ruperao, P., Cheng, F., Waminal, N.E., Huang, Y., Dong, C., Wang, L., Li, J., Hu, Z., Zhuang, M., Huang, Y., Huang, J., Shi, J., Mei, D., Liu, J., Lee, T.H., Wang, J., Jin, H., Li, Z., Li, X., Zhang, J., Xiao, L., Zhou, Y., Liu, Z., Liu, X., Qin, R., Tang, X., Liu, W., Wang, Y., Zhang, Y., Lee, J., Kim, H.H., Denoeud, F., Xu, X., Liang, X., Hua, W., Wang, X., Wang, J., Chalhoub, B., and Paterson, A.H. (2014). The *Brassica oleracea* genome reveals the asymmetrical evolution of polyploid genomes. Nat Commun 5, 3930.

Lohse, M., Nagel, A., Herter, T., May, P., Schroda, M., Zrenner, R., Tohge, T., Fernie, A.R., Stitt, M., and Usadel, B. (2014). Mercator: a fast and simple web server for genome scale functional annotation of plant sequence data. Plant Cell Environ. 37, 1250–1258.

Love, M.I., Huber, W., and Anders, S. (2014). Moderated estimation of fold change and dispersion for RNA-seq data with DESeq2. Genome Biol. 15, 550.

Lyons, E., Pedersen, B., Kane, J., and Freeling, M. (2008). The Value of Nonmodel Genomes and an Example Using SynMap Within CoGe to Dissect the Hexaploidy that Predates the Rosids. Trop. Plant Biol. 1, 181–190.

Lysak, M.A., Mandakova, T., and Schranz, M.E. (2016). Comparative paleogenomics of crucifers: ancestral genomic blocks revisited. Curr. Opin. Plant Biol. 30, 108–115.

Mabry, M.E., Brose, J.M., Blischak, P.D., Sutherland, B., Dismukes, W.T., Bottoms, C.A., Edger, P.P., Washburn, J.D., An, H., Hall, J.C., McKain, M.R., Al-Shehbaz, I., Barker, M.S., Schranz, M.E., Conant, G.C., and Pires, J.C. (2020). Phylogeny and multiple independent whole-genome duplication events in the Brassicales. Am. J. Bot. 107, 1148–1164.

Majoros, W.H., Pertea, M., and Salzberg, S.L. (2004). TigrScan and GlimmerHMM: two open source ab initio eukaryotic gene-finders. Bioinformatics 20, 2878–2879.

Minh, B.Q., Schmidt, H.A., Chernomor, O., Schrempf, D., Woodhams, M.D., von Haeseler, A., and Lanfear, R. (2020). IQ-TREE 2: New Models and Efficient Methods for Phylogenetic Inference in the Genomic Era. Mol Biol Evol 37, 1530–1534.

Miyagishima, S.Y., Froehlich, J.E., and Osteryoung, K.W. (2006). PDV1 and PDV2 mediate recruitment of the dynamin-related protein ARC5 to the plastid division site. Plant Cell 18, 2517–2530.

Mravec, J., Skupa, P., Bailly, A., Hoyerova, K., Krecek, P., Bielach, A., Petrasek, J., Zhang, J., Gaykova, V., Stierhof, Y.D., Dobrev, P.I., Schwarzerova, K., Rolcik, J., Seifertova, D., Luschnig, C., Benkova, E., Zazimalova, E., Geisler, M., and Friml, J. (2009). Subcellular homeostasis of phytohormone auxin is mediated by the ER-localized PIN5 transporter. Nature 459, 1136–1140.

Nevers, Y., Rossier, V., Train, C.M., Altenhoff, A., Dessimoz, C., and Glover, N. (2022). Multifaceted quality assessment of gene repertoire annotation with OMArk. bioRxiv, 2022.2011.2025.517970.

Nikolov, L.A., Shushkov, P., Nevado, B., Gan, X., Al-Shehbaz, I.A., Filatov, D., Bailey, C.D., and Tsiantis, M. (2019). Resolving the backbone of the Brassicaceae phylogeny for investigating trait diversity. New Phytol. 222, 1638–1651.

O’Donovan, C., Martin, M.J., Gattiker, A., Gasteiger, E., Bairoch, A., and Apweiler, R. (2002). High-quality protein knowledge resource: SWISS-PROT and TrEMBL. Brief. Bioinform. 3, 275–284.

Parkin, I.A., Koh, C., Tang, H., Robinson, S.J., Kagale, S., Clarke, W.E., Town, C.D., Nixon, J., Krishnakumar, V., Bidwell, S.L., Denoeud, F., Belcram, H., Links, M.G., Just, J., Clarke, C., Bender, T., Huebert, T., Mason, A.S., Pires, J.C., Barker, G., Moore, J., Walley, P.G., Manoli, S., Batley, J., Edwards, D., Nelson, M.N., Wang, X., Paterson, A.H., King, G., Bancroft, I., Chalhoub, B., and Sharpe, A.G. (2014). Transcriptome and methylome profiling reveals relics of genome dominance in the mesopolyploid *Brassica oleracea*. Genome Biol. 15, R77.

Perumal, S., Koh, C.S., Jin, L., Buchwaldt, M., Higgins, E.E., Zheng, C., Sankoff, D., Robinson, S.J., Kagale, S., Navabi, Z.K., Tang, L., Horner, K.N., He, Z., Bancroft, I., Chalhoub, B., Sharpe, A.G., and Parkin, I.A.P. (2020). A high-contiguity *Brassica nigra* genome localizes active centromeres and defines the ancestral *Brassica* genome. Nat Plants 6, 929–941.

Pham, S.K., and Pevzner, P.A. (2010). DRIMM-Synteny: decomposing genomes into evolutionary conserved segments. Bioinformatics 26, 2509–2516.

Price, M.N., Dehal, P.S., and Arkin, A.P. (2009). FastTree: computing large minimum evolution trees with profiles instead of a distance matrix. Mol Biol Evol 26, 1641–1650.

Qiao, X., Li, Q., Yin, H., Qi, K., Li, L., Wang, R., Zhang, S., and Paterson, A.H. (2019). Gene duplication and evolution in recurring polyploidization-diploidization cycles in plants. Genome Biol. 20, 38.

Sawchuk, M.G., Edgar, A., and Scarpella, E. (2013). Patterning of leaf vein networks by convergent auxin transport pathways. PLoS Genet. 9, e1003294.

Schluter, U., Bouvier, J.W., Guerreiro, R., Malisic, M., Kontny, C., Westhoff, P., Stich, B., and Weber, A.P.M. (2023). Brassicaceae display variation in efficiency of photorespiratory carbon-recapturing mechanisms. J. Exp. Bot. 74, 6631–6649.

Scholes, D.R., and Paige, K.N. (2015). Plasticity in ploidy: a generalized response to stress. Trends Plant Sci. 20, 165–175.

Schranz, M.E., Lysak, M.A., and Mitchell-Olds, T. (2006). The ABC’s of comparative genomics in the Brassicaceae: building blocks of crucifer genomes. Trends Plant Sci. 11, 535–542.

Shumate, A., and Salzberg, S.L. (2021). Liftoff: accurate mapping of gene annotations. Bioinformatics 37, 1639–1643.

Simao, F.A., Waterhouse, R.M., Ioannidis, P., Kriventseva, E.V., and Zdobnov, E.M. (2015). BUSCO: assessing genome assembly and annotation completeness with single-copy orthologs. Bioinformatics 31, 3210–3212.

Smith, M.L., and Hahn, M.W. (2021). New Approaches for Inferring Phylogenies in the Presence of Paralogs. Trends Genet. 37, 174–187.

Smith, S.A., Moore, M.J., Brown, J.W., and Yang, Y. (2015). Analysis of phylogenomic datasets reveals conflict, concordance, and gene duplications with examples from animals and plants. BMC Evol. Biol. 15, 150.

Stanke, M., and Morgenstern, B. (2005). AUGUSTUS: a web server for gene prediction in eukaryotes that allows user-defined constraints. Nucleic Acids Res. 33, W465–467.

Stevens, A.V., Nicotra, A.B., Godfree, R.C., and Guja, L.K. (2020). Polyploidy affects the seed, dormancy and seedling characteristics of a perennial grass, conferring an advantage in stressful climates. Plant Biol. (Stuttg.) 22, 500–513.

Suyama, M., Torrents, D., and Bork, P. (2006). PAL2NAL: robust conversion of protein sequence alignments into the corresponding codon alignments. Nucleic Acids Res. 34, W609–612.

Tang, H., Bowers, J.E., Wang, X., Ming, R., Alam, M., and Paterson, A.H. (2008). Synteny and collinearity in plant genomes. Science 320, 486–488.

Tang, H., Bomhoff, M.D., Briones, E., Zhang, L., Schnable, J.C., and Lyons, E. (2015). SynFind: Compiling Syntenic Regions across Any Set of Genomes on Demand. Genome Biol. Evol. 7, 3286–3298.

Tarailo-Graovac, M., and Chen, N. (2009). Using RepeatMasker to identify repetitive elements in genomic sequences. Curr Protoc Bioinformatics Chapter 4, 4 10 11-14 10 14.

Theeuwen, T.P.J.M., Logie, L.L., Harbinson, J., and Aarts, M.G.M. (2022). Genetics as a key to improving crop photosynthesis. J. Exp. Bot. 73, 3122–3137.

Tholen, D., Boom, C., and Zhu, X.G. (2012). Opinion: prospects for improving photosynthesis by altering leaf anatomy. Plant Sci. 197, 92–101.

Trifinopoulos, J., Nguyen, L.T., von Haeseler, A., and Minh, B.Q. (2016). W-IQ-TREE: a fast online phylogenetic tool for maximum likelihood analysis. Nucleic Acids Res. 44, W232–235.

UniProt, C. (2015). UniProt: a hub for protein information. Nucleic Acids Res. 43, D204–212.

Van de Peer, Y., Mizrachi, E., and Marchal, K. (2017). The evolutionary significance of polyploidy. Nat. Rev. Genet. 18, 411–424.

Vaser, R., Sovic, I., Nagarajan, N., and Sikic, M. (2017). Fast and accurate de novo genome assembly from long uncorrected reads. Genome Res. 27, 737–746.

Vieira, P., De Clercq, A., Stals, H., Van Leene, J., Van De Slijke, E., Van Isterdael, G., Eeckhout, D., Persiau, G., Van Damme, D., Verkest, A., Antonino de Souza, J.D., Junior, Glab, N., Abad, P., Engler, G., Inze, D., De Veylder, L., De Jaeger, G., and Engler, J.D. (2014). The Cyclin-Dependent Kinase Inhibitor KRP6 Induces Mitosis and Impairs Cytokinesis in Giant Cells Induced by Plant-Parasitic Nematodes in Arabidopsis. Plant Cell 26, 2633–2647.

Vurture, G.W., Sedlazeck, F.J., Nattestad, M., Underwood, C.J., Fang, H., Gurtowski, J., and Schatz, M.C. (2017). GenomeScope: fast reference-free genome profiling from short reads. Bioinformatics 33, 2202–2204.

Waidmann, S., Beziat, C., Ferreira Da Silva Santos, J., Feraru, E., Feraru, M.I., Sun, L., Noura, S., Boutte, Y., and Kleine-Vehn, J. (2023). Endoplasmic reticulum stress controls PIN-LIKES abundance and thereby growth adaptation. Proc. Natl. Acad. Sci. U. S. A. 120, e2218865120.

Walden, N., and Schranz, M.E. (2023). Synteny Identifies Reliable Orthologs for Phylogenomics and Comparative Genomics of the Brassicaceae. Genome Biol. Evol. 15.

Wang, D., Zhang, Y., Zhang, Z., Zhu, J., and Yu, J. (2010). KaKs_Calculator 2.0: a toolkit incorporating gamma-series methods and sliding window strategies. Genomics Proteomics Bioinformatics 8, 77–80.

Wang, D.P., Wan, H.L., Zhang, S., and Yu, J. (2009a). Gamma-MYN: a new algorithm for estimating Ka and Ks with consideration of variable substitution rates. Biol. Direct 4, 20.

Wang, X., Gowik, U., Tang, H., Bowers, J.E., Westhoff, P., and Paterson, A.H. (2009b). Comparative genomic analysis of C4 photosynthetic pathway evolution in grasses. Genome Biol. 10, R68.

Wang, X., Wang, H., Wang, J., Sun, R., Wu, J., Liu, S., Bai, Y., Mun, J.H., Bancroft, I., Cheng, F., Huang, S., Li, X., Hua, W., Wang, J., Wang, X., Freeling, M., Pires, J.C., Paterson, A.H., Chalhoub, B., Wang, B., Hayward, A., Sharpe, A.G., Park, B.S., Weisshaar, B., Liu, B., Li, B., Liu, B., Tong, C., Song, C., Duran, C., Peng, C., Geng, C., Koh, C., Lin, C., Edwards, D., Mu, D., Shen, D., Soumpourou, E., Li, F., Fraser, F., Conant, G., Lassalle, G., King, G.J., Bonnema, G., Tang, H., Wang, H., Belcram, H., Zhou, H., Hirakawa, H., Abe, H., Guo, H., Wang, H., Jin, H., Parkin, I.A., Batley, J., Kim, J.S., Just, J., Li, J., Xu, J., Deng, J., Kim, J.A., Li, J., Yu, J., Meng, J., Wang, J., Min, J., Poulain, J., Wang, J., Hatakeyama, K., Wu, K., Wang, L., Fang, L., Trick, M., Links, M.G., Zhao, M., Jin, M., Ramchiary, N., Drou, N., Berkman, P.J., Cai, Q., Huang, Q., Li, R., Tabata, S., Cheng, S., Zhang, S., Zhang, S., Huang, S., Sato, S., Sun, S., Kwon, S.J., Choi, S.R., Lee, T.H., Fan, W., Zhao, X., Tan, X., Xu, X., Wang, Y., Qiu, Y., Yin, Y., Li, Y., Du, Y., Liao, Y., Lim, Y., Narusaka, Y., Wang, Y., Wang, Z., Li, Z., Wang, Z., Xiong, Z., Zhang, Z., and Brassica rapa Genome Sequencing Project, C. (2011). The genome of the mesopolyploid crop species *Brassica rapa*. Nat. Genet. 43, 1035–1039.

Wang, Y., Tang, H., Debarry, J.D., Tan, X., Li, J., Wang, X., Lee, T.H., Jin, H., Marler, B., Guo, H., Kissinger, J.C., and Paterson, A.H. (2012). MCScanX: a toolkit for detection and evolutionary analysis of gene synteny and collinearity. Nucleic Acids Res. 40, e49.

Wu, A., Allu, A.D., Garapati, P., Siddiqui, H., Dortay, H., Zanor, M.I., Asensi-Fabado, M.A., Munne-Bosch, S., Antonio, C., Tohge, T., Fernie, A.R., Kaufmann, K., Xue, G.P., Mueller-Roeber, B., and Balazadeh, S. (2012). JUNGBRUNNEN1, a reactive oxygen species-responsive NAC transcription factor, regulates longevity in Arabidopsis. Plant Cell 24, 482–506.

Wu, J., Liang, J., Lin, R., Cai, X., Zhang, L., Guo, X., Wang, T., Chen, H., and Wang, X. (2022). Investigation of *Brassica* and its relative genomes in the post-genomics era. Hortic Res 9, uhac182.

Xiong, H., Wang, D., Shao, C., Yang, X., Yang, J., Ma, T., Davis, C.C., Liu, L., and Xi, Z. (2022). Species Tree Estimation and the Impact of Gene Loss Following Whole-Genome Duplication. Syst. Biol. 71, 1348–1361.

Xu, H., Wang, C., Shao, G., Wu, S., Liu, P., Cao, P., Jiang, P., Wang, S., Zhu, H., Lin, X., Tauqeer, A., Lin, Y., Chen, W., Huang, W., Wen, Q., Chang, J., Zhong, F., and Wu, S. (2022). The reference genome and full-length transcriptome of pakchoi provide insights into cuticle formation and heat adaption. Hortic Res 9, uhac123.

Xu, L., Wang, Y., Dong, J., Zhang, W., Tang, M., Zhang, W., Wang, K., Chen, Y., Zhang, X., He, Q., Zhang, X., Wang, K., Wang, L., Ma, Y., Xia, K., and Liu, L. (2023). A chromosome-level genome assembly of radish (*Raphanus sativus* L.) reveals insights into genome adaptation and differential bolting regulation. Plant Biotechnol. J. 21, 990–1004.

Yang, T., Cai, B., Jia, Z., Wang, Y., Wang, J., King, G.J., Ge, X., and Li, Z. (2023). Sinapis genomes provide insights into whole-genome triplication and divergence patterns within tribe Brassiceae. Plant J. 113, 246–261.

Yang, Z. (2007). PAML 4: phylogenetic analysis by maximum likelihood. Mol Biol Evol 24, 1586–1591.

Yang, Z., Yan, B., Dong, H., He, G., Zhou, Y., and Sun, J. (2021). BIC1 acts as a transcriptional coactivator to promote brassinosteroid signaling and plant growth. EMBO J. 40, e104615.

Yim, W.C., Swain, M.L., Ma, D., An, H., Bird, K.A., Curdie, D.D., Wang, S., Ham, H.D., Luzuriaga-Neira, A., Kirkwood, J.S., Hur, M., Solomon, J.K.Q., Harper, J.F., Kosma, D.K., Alvarez-Ponce, D., Cushman, J.C., Edger, P.P., Mason, A.S., Pires, J.C., Tang, H., and Zhang, X. (2022). The final piece of the Triangle of U: Evolution of the tetraploid *Brassica carinata* genome. Plant Cell 34, 4143–4172.

Zdobnov, E.M., and Apweiler, R. (2001). InterProScan--an integration platform for the signature-recognition methods in InterPro. Bioinformatics 17, 847–848.

Zedek, F., Plackova, K., Vesely, P., Smerda, J., Smarda, P., Horova, L., and Bures, P. (2020). Endopolyploidy is a common response to UV-B stress in natural plant populations, but its magnitude may be affected by chromosome type. Am. J. Bot. 126, 883–889.

Zhang, C., Rabiee, M., Sayyari, E., and Mirarab, S. (2018). ASTRAL-III: polynomial time species tree reconstruction from partially resolved gene trees. BMC Bioinformatics 19, 153.

Zhang, C., Scornavacca, C., Molloy, E.K., and Mirarab, S. (2020). ASTRAL-Pro: Quartet-Based Species-Tree Inference despite Paralogy. Mol Biol Evol 37, 3292–3307.

Zhang, L., Liang, J., Chen, H., Zhang, Z., Wu, J., and Wang, X. (2023). A near-complete genome assembly of *Brassica rapa* provides new insights into the evolution of centromeres. Plant Biotechnol. J. 21, 1022–1032.

Zhu, X.G., Wang, Y., Ort, D.R., and Long, S.P. (2013). e-Photosynthesis: a comprehensive dynamic mechanistic model of C3 photosynthesis: from light capture to sucrose synthesis. Plant Cell Environ. 36, 1711–1727.

