## Supplemental Figure file for "Expanding the Triangle of U: The genome assembly of *Hirschfeldia incana* provides insights into chromosomal evolution, phylogenomics and high photosynthesis-related traits"

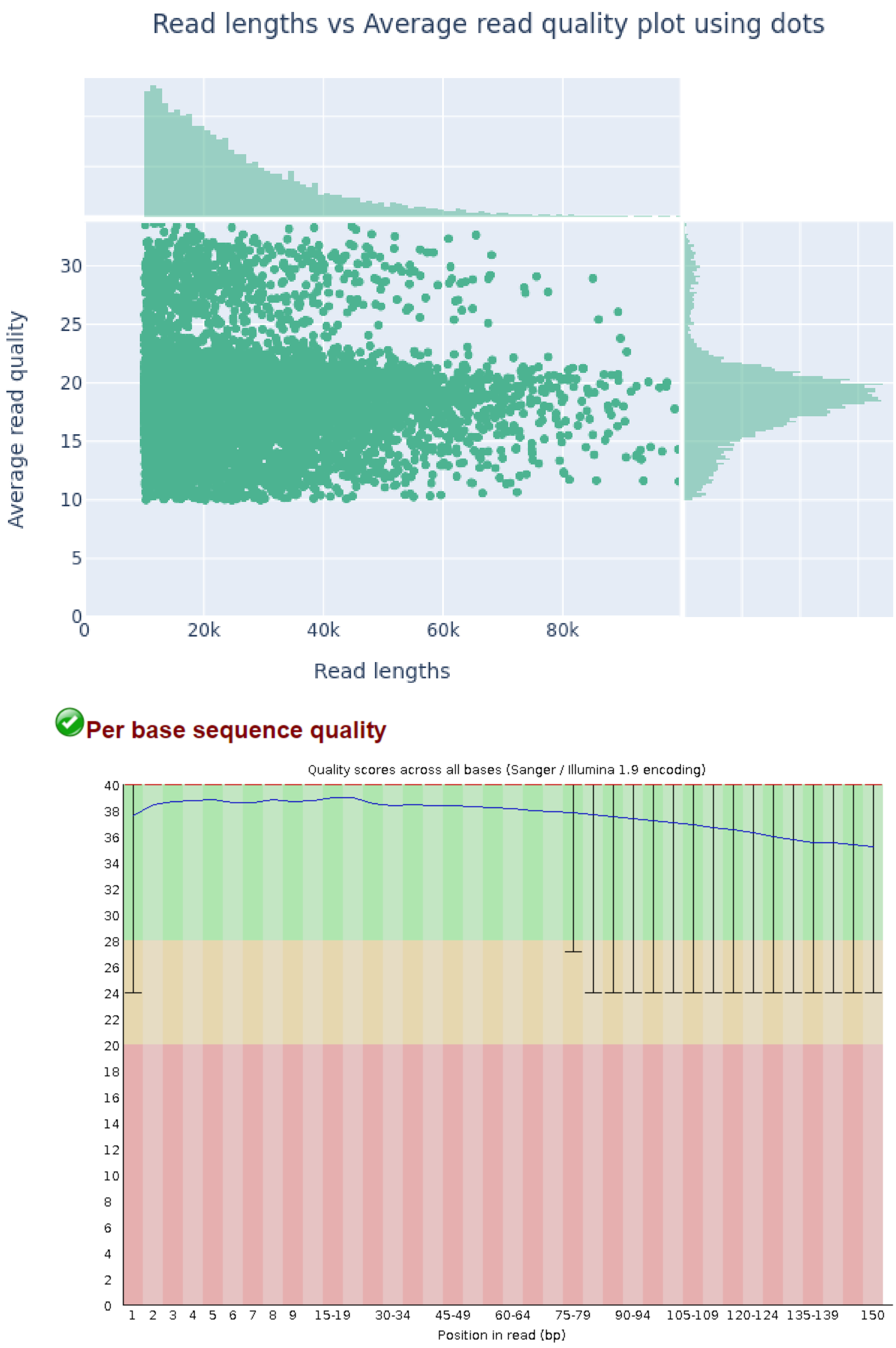


**Figure S1. Summary of the combined Nanopore ONT whole-genome sequencing data (upper panel) and Hi-C data base quality (lower panel).** A total of ~20 Gb long-read ONT data (N50 = 26 kb, ~48x genome coverage) and 124 million Hi-C Illumina reads (150 bp, ~44x genome overage) were generated.

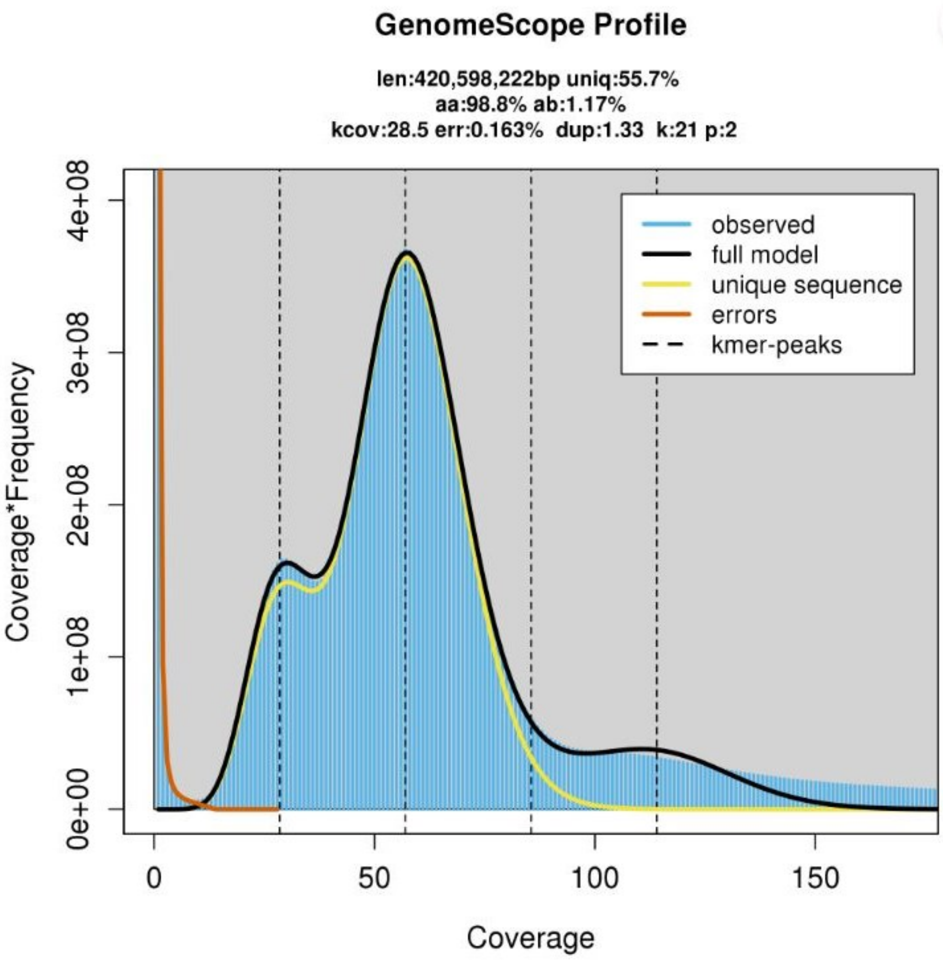


**Figure S2. Size estimation of the *H. incana* genome** using GenomeScope v2.0 and 102 millions WGS Illumina read data published in Garassino et al. (2022).

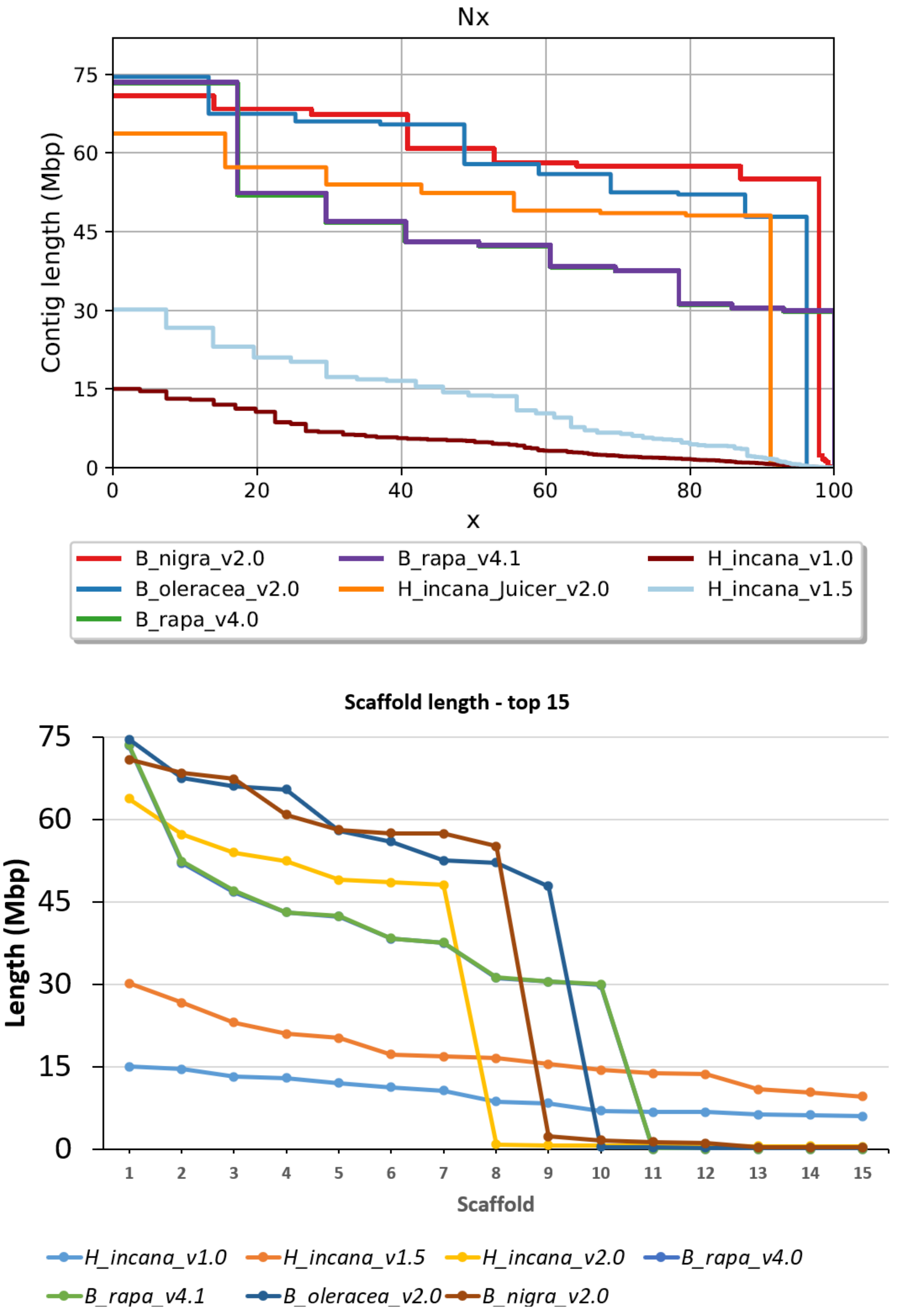


**Figure S3. Comparison among the assemblies from the Brassiceae.** Total length comparison (upper panel) and top 15 longest scaffolds (lower panel). In the upper panel: x denotes % of assembly.

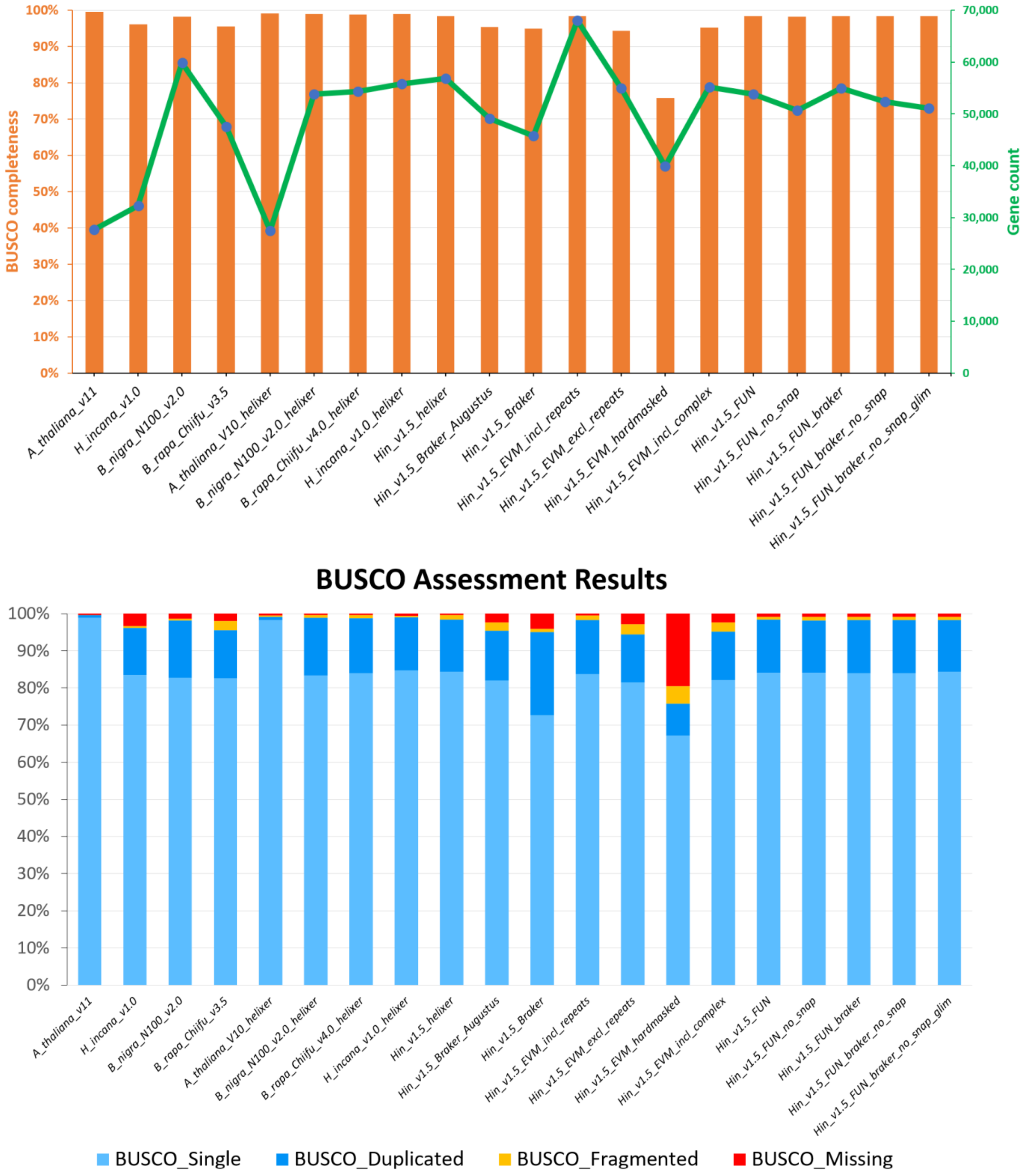


**Figure S4. Comparison of different annotation approaches of the *H. incana* genome v1.5.** The genomes of *A. thaliana* Col-0, *B. rapa* Chiifu, *B. oleracea* JZS, *B. nigra* NI100. Annotation with “helixer” in the names indicate those that were reannotated using Helixer (Holst et al., 2023). EVM denotes EvidenceModeler (Haas et al., 2008). FUN indicates Funannotate derived annotation (https://github.com/nextgenusfs/funannotate). All FUN annotation sets were obtained from PREDICT step of the pipeline. The final annotation was derived from Hin_v1.5_FUN after the UPDATE step by PASA to add untranslated regions (UTR) data and fix gene models that were not in agreement with the RNA-seq data. This update annotation was then lifted over to genome v2.0 (Hi-C) to obtain the final annotation. *Hin: H. incana.*

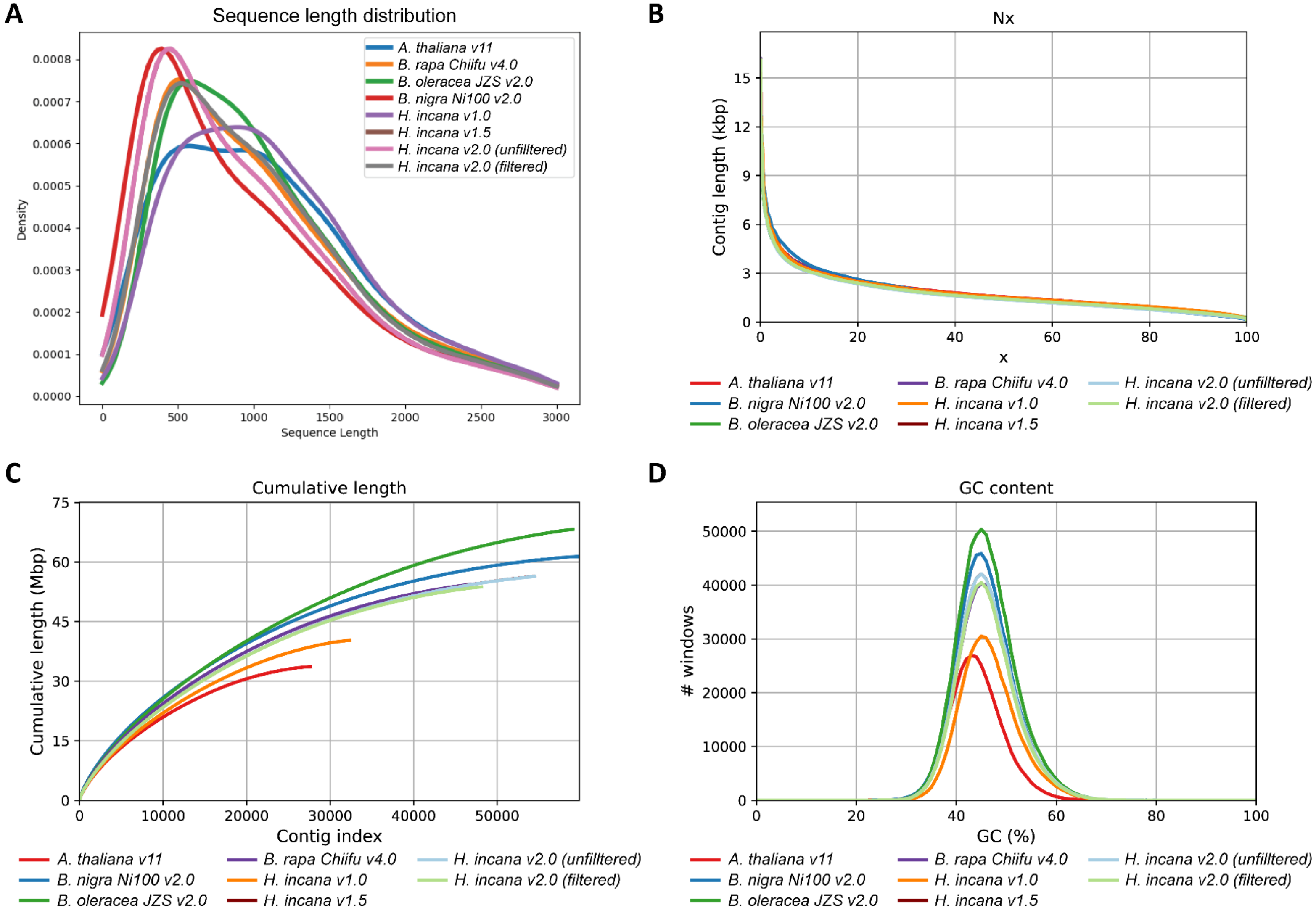


**Figure S5. Comparison of annotated proteomes and coding sequences (CDS) from selected Brassicaceae genomes. (A)** Protein length distribution. **(B)** Cumulative length in percentage of CDS. x denotes % of assembly. **(C)** Cumulative length in Mb of CDS. **(D)** GC content of CDS. Note that the distribution of *H. incana* v1.5 and v2.0 are very similar and largely overlapped.

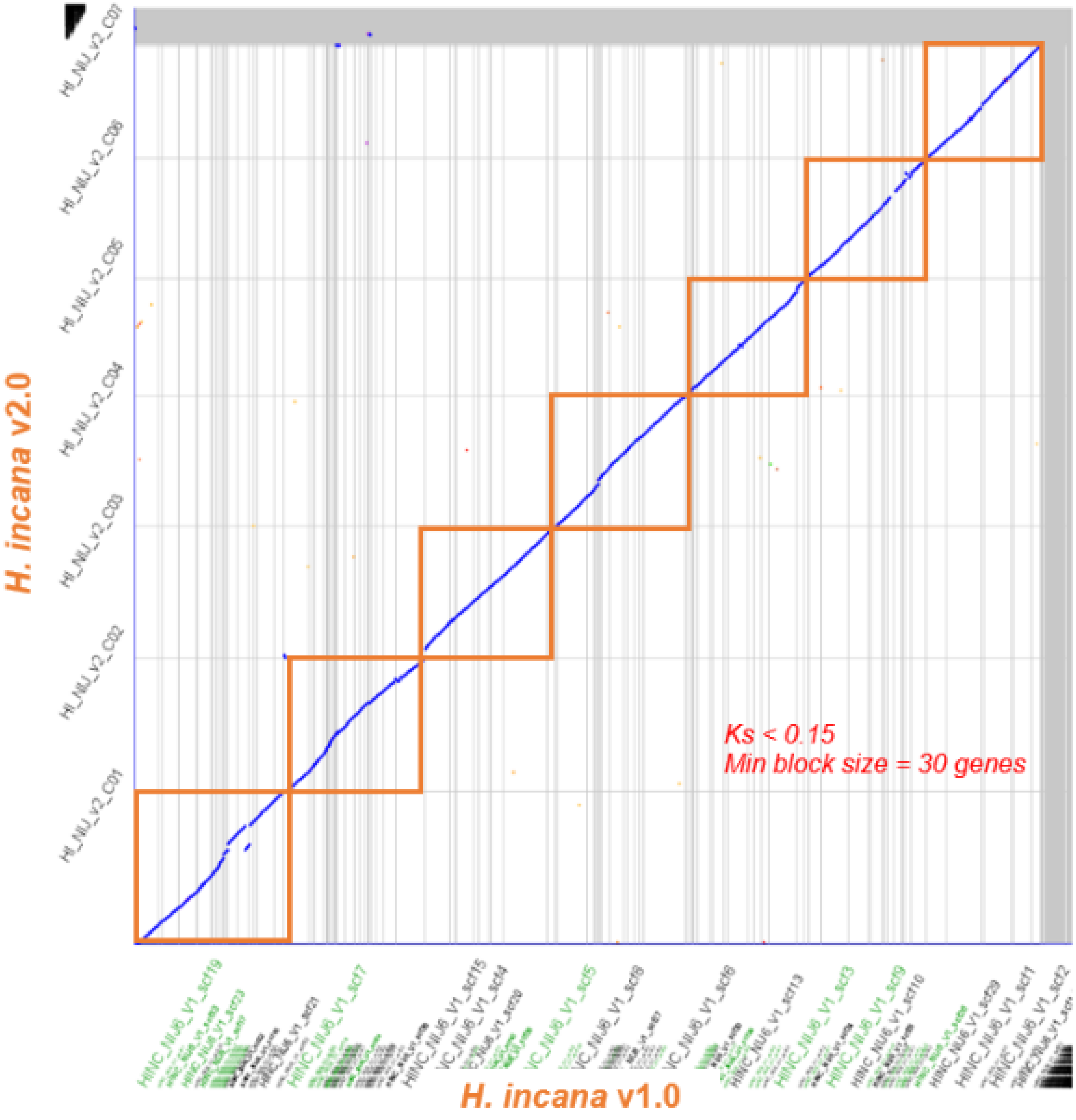


**Figure S6. Dotplot showing synteny between *H. incana* v1.0 and v2.0, analyzed by SynMap program** on the CoGe v7 (https://genomevolution.org/coge/) using a syntenic block size minimum = 30 genes. Only "true orthologs" between the two assemblies were used**.**

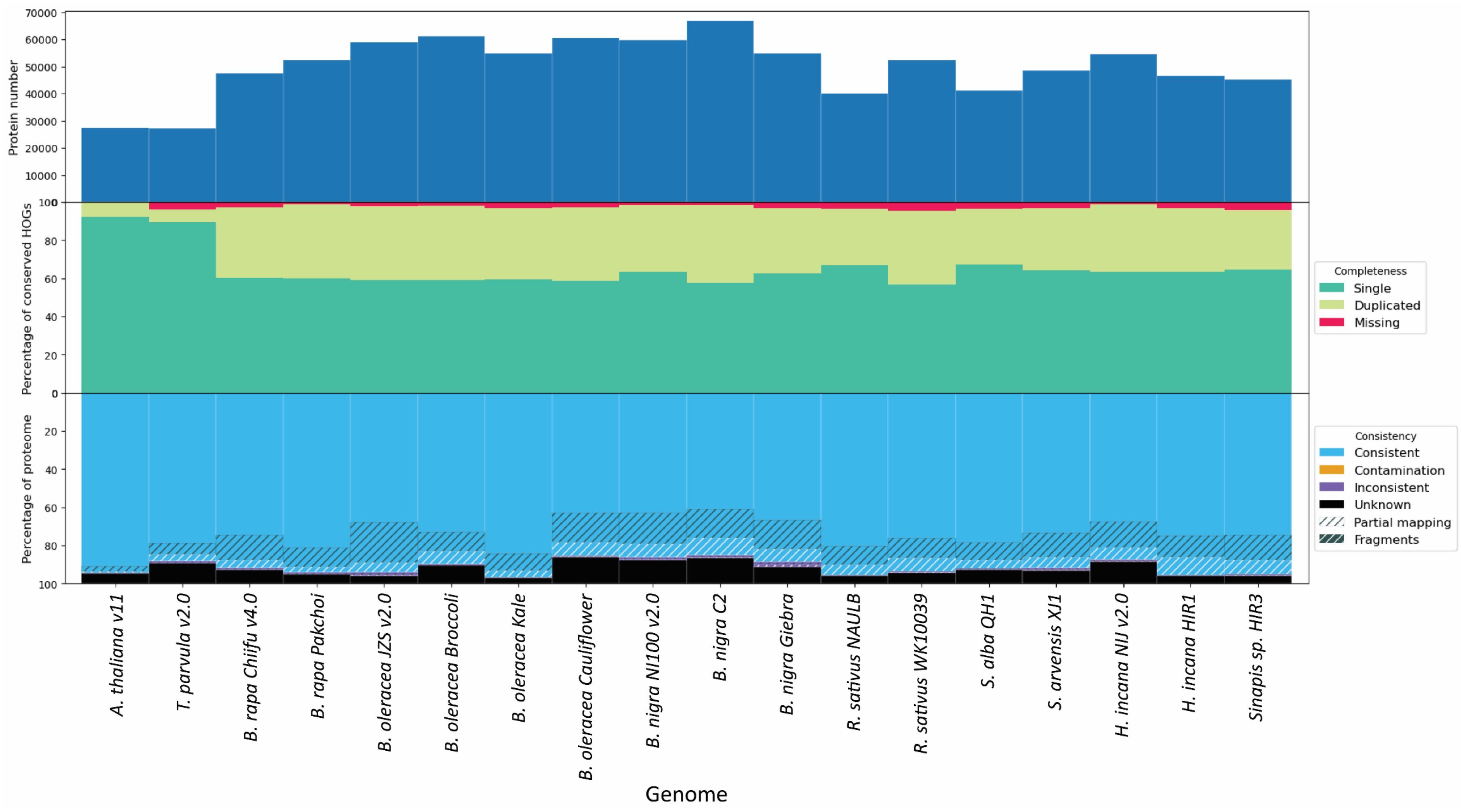


**Figure S7. OMARK proteome assessment of selected Brassicaceae genomes.** The OMArk v0.3.0 based on 17,999 conserved orthologs of the Brassicaceae family (Nevers et al., 2022) were used to analyze and compare the annotations.

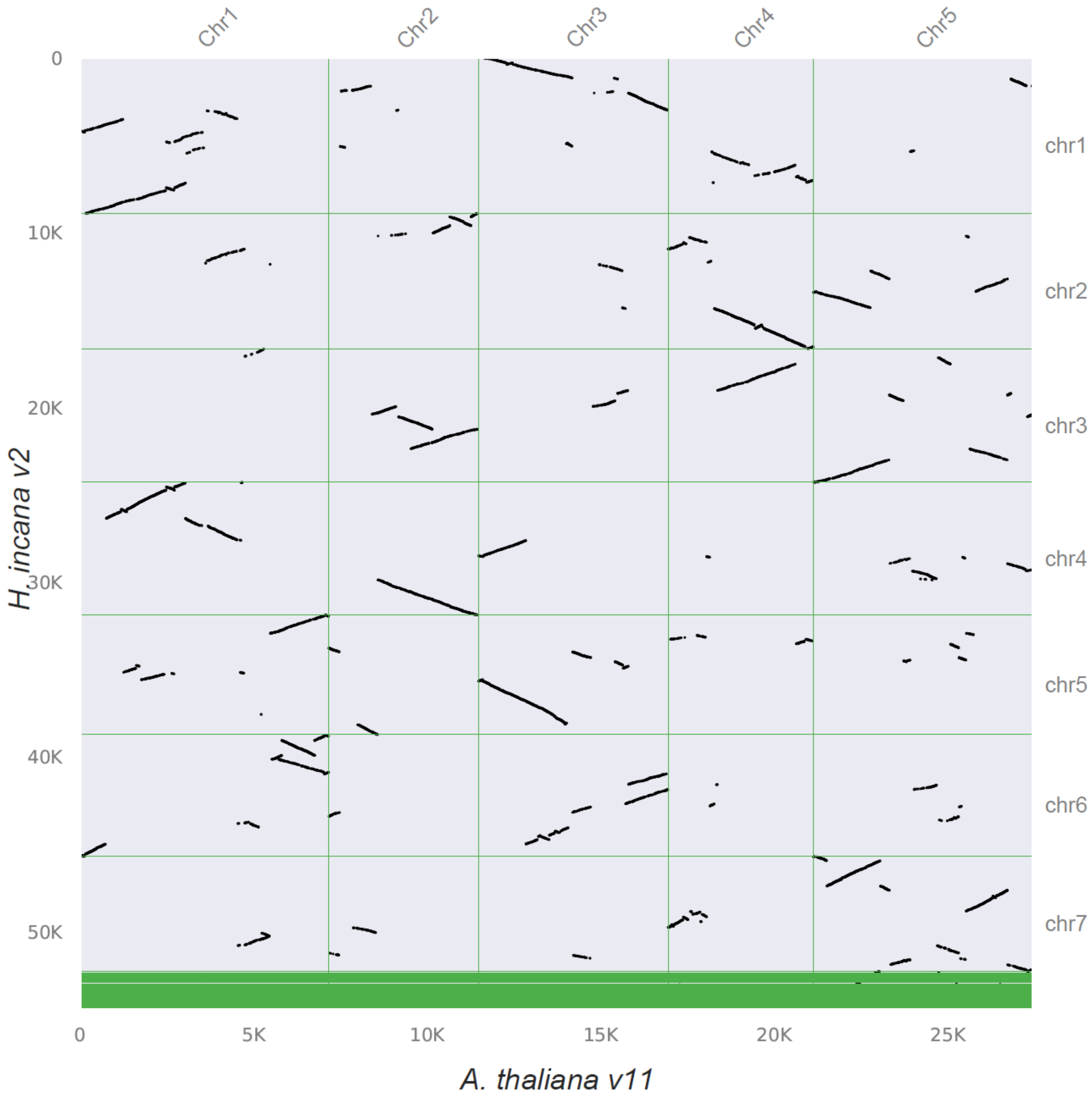


**Figure S8. Macro-synteny between *A. thaliana* and *H. incana*.** The syntenic depth ratio was set to 1 vs. 3 to use only true orthologous genes between the 2 genomes. Plot was generated by MCscan v0.8 (Tang et al., 2008) python version (https://github.com/tanghaibao/jcvi/wiki/MCscan-(Python-version))

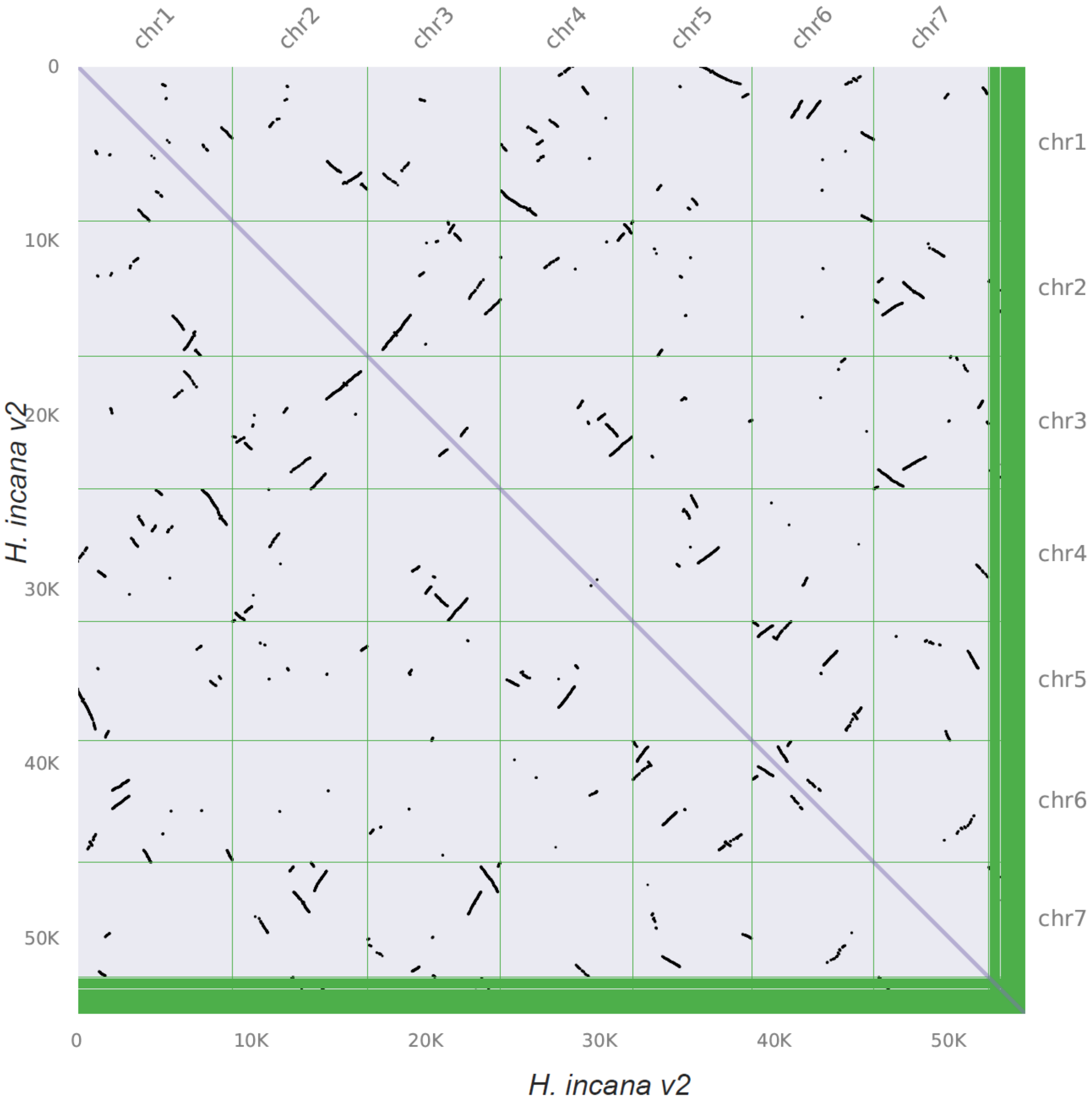


**Figure S9. Intra-genomic syntenic (self-comparison) dotplot the *H. incana* genome v2.0.** Plot was generated by MCscan v0.8 (Tang et al., 2008) python version (https://github.com/tanghaibao/jcvi/wiki/MCscan-(Python-version))

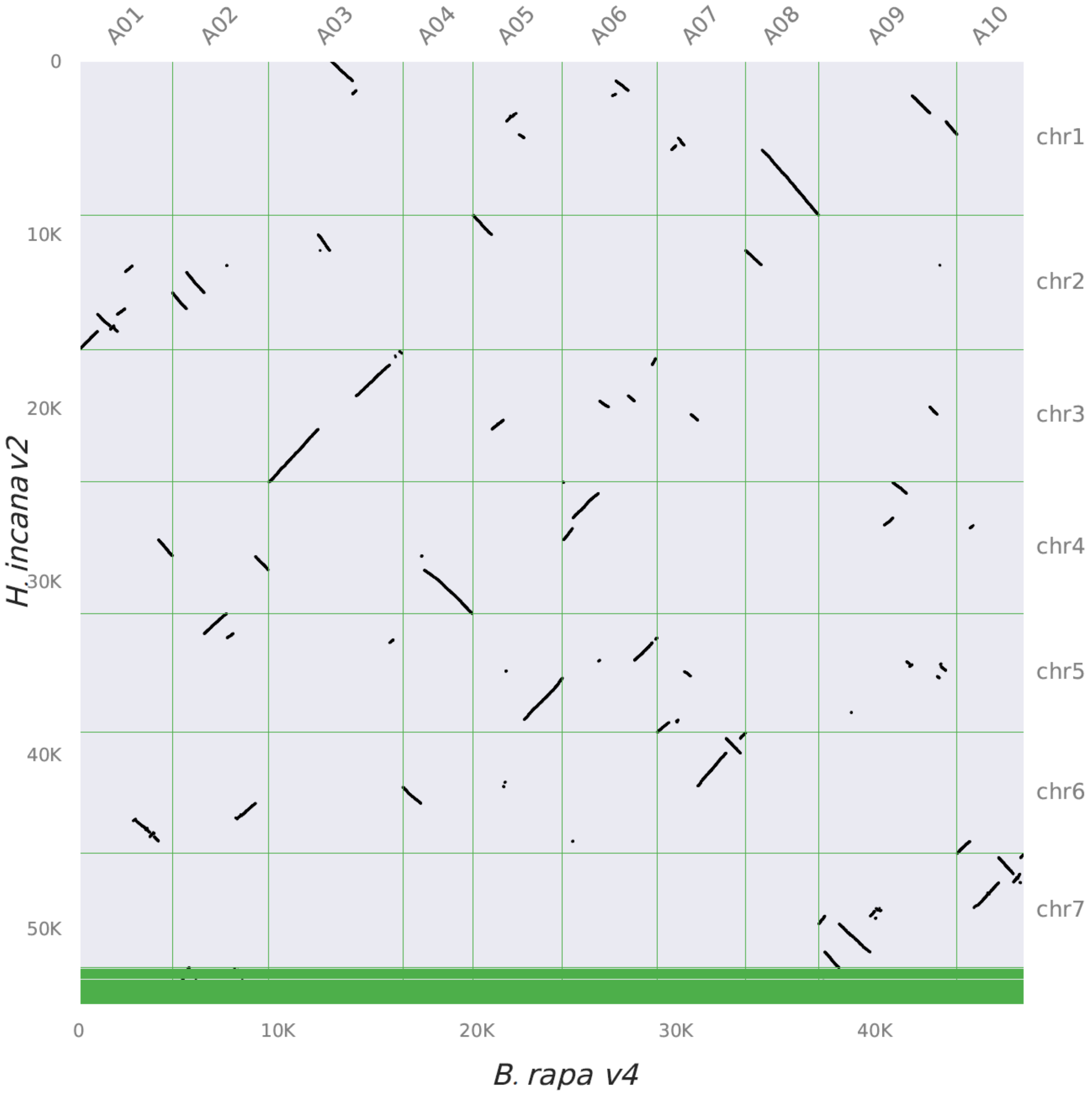


**Figure S10. Macro-synteny between *B. rapa* and *H. incana*.** The syntenic depth ratio was set to 1 vs. 1 to use only true orthologous genes between the 2 genomes. Plot was generated by MCscan v0.8 (Tang et al., 2008) python version (https://github.com/tanghaibao/jcvi/wiki/MCscan-(Python-version))

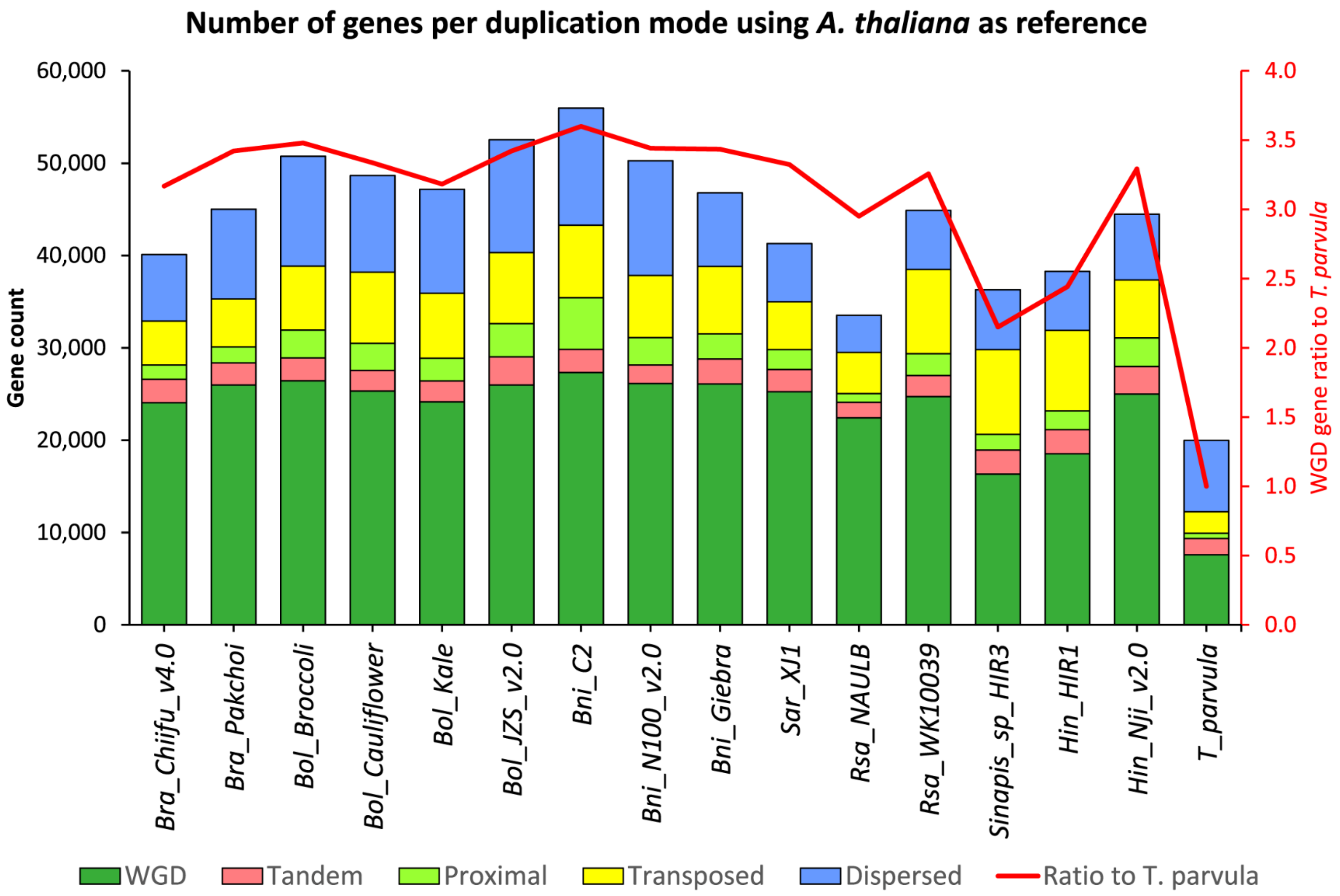


**Figure S11. Gene duplication modes of the *H. incana* genome v2.0 compared with other Brassicaceae genomes.** Modes of duplicated gene copies were analyzed by DupGen_finder (Qiao et al., 2019) with default parameters using the *A. thaliana* genome as reference. *Tpa: T. parvula, Bra: B. rapa, Bol: B. oleracea, Bni: B. nigra, Rsa: R. sativus, Sar: S. arvensis,* and *Hin: H. incana.* Note that *A. thaliana* and *T. parvula* did not experience the WGT *Br-α* event.

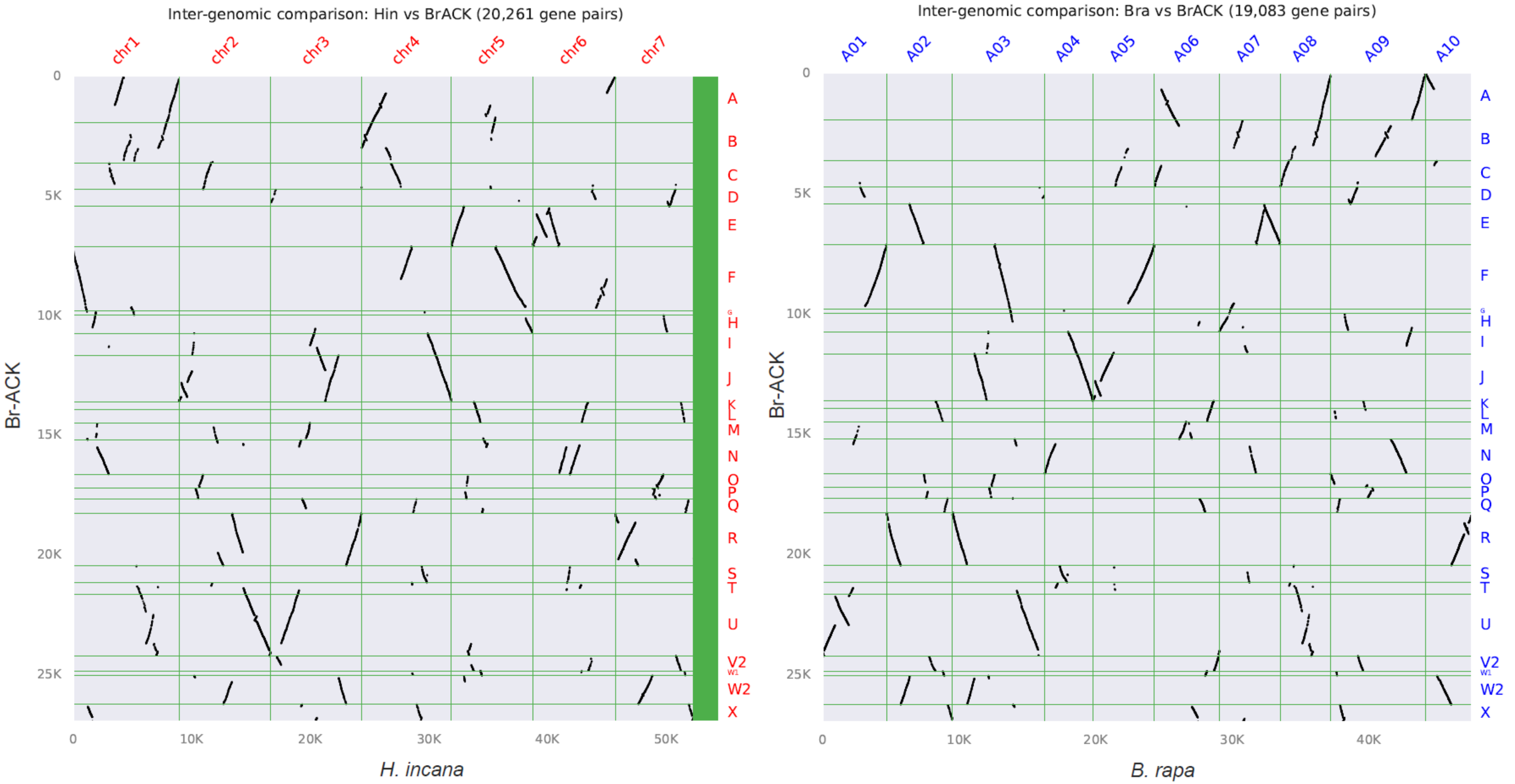


**Figure S12. Syntenic relationship between the *H. incana* genome v2.0 and the Brassica tPCK ancestral genomic blocks (Bra-ACK).** For comparison, the analysis was also done for the *B. rapa* genome v4.0. Plot was generated by MCscan v0.8 (Tang et al., 2008) python version (https://github.com/tanghaibao/jcvi/wiki/MCscan-(Python-version))

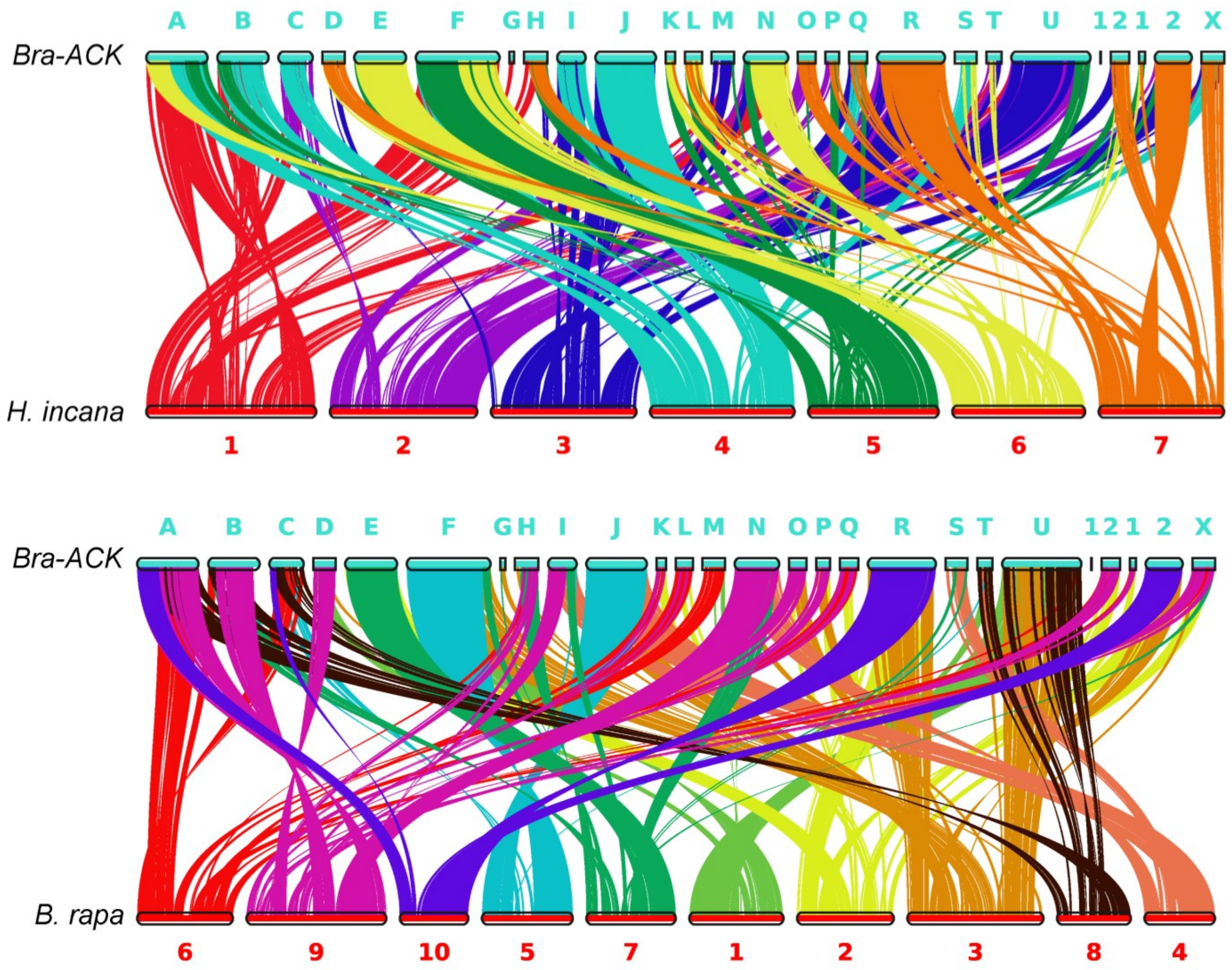


**Figure S13. Additional plot showing syntenic relationship between the *H. incana* genome v2.0 and the Brassica tPCK ancestral genomic blocks (Bra-ACK).** Same genomes and comparisons as in **Figure S12** were used. The blocks between U and X are V1, V2, W1 and W2.

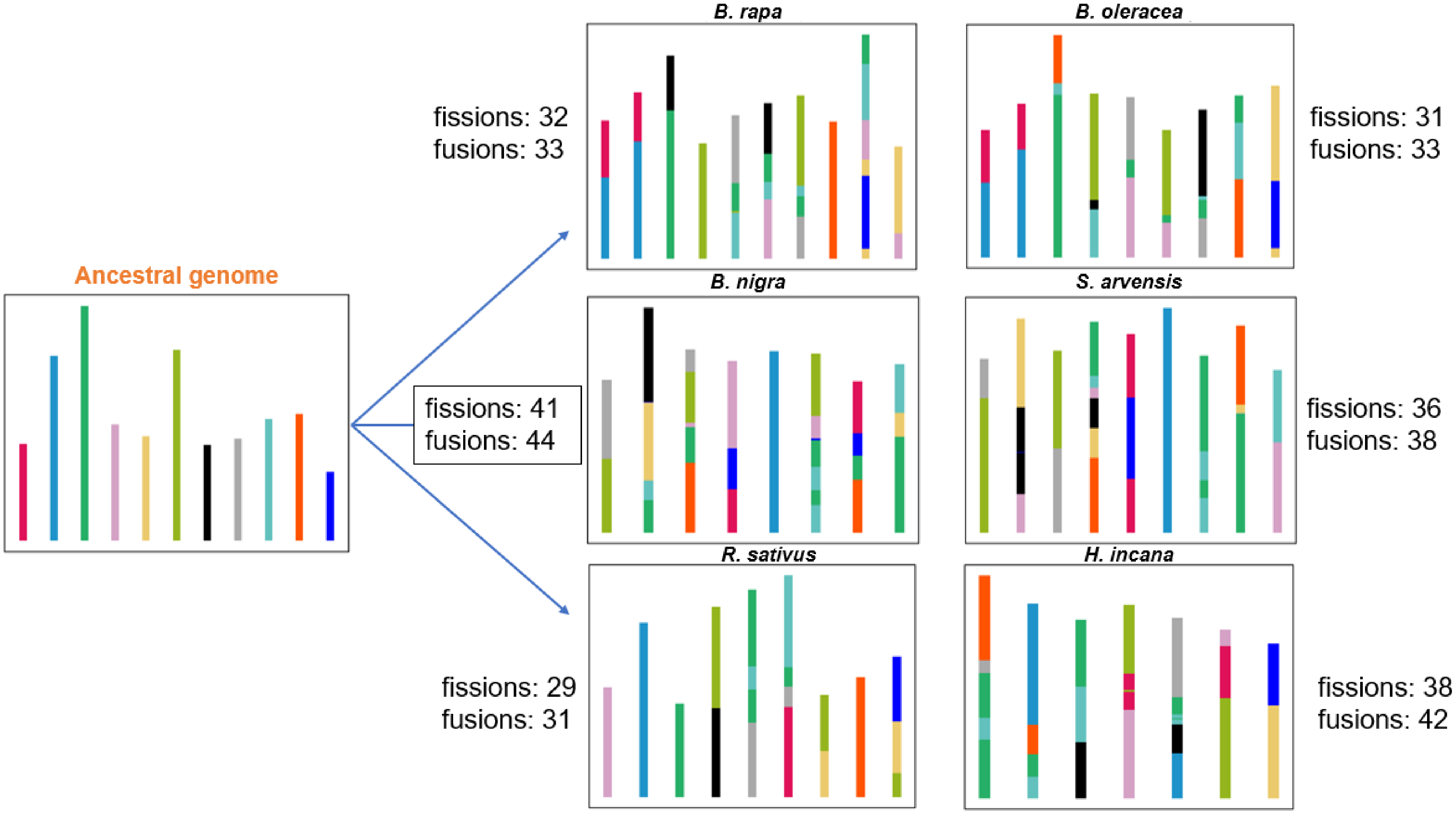


**Figure S14. Evolutionary history and genome rearrangement estimation of the six Brassiceae genomes.** The genomes of *B. rapa* Chiifu, *B. oleracea* JZS, *B. nigra* NI100, *S. arvensis* XJ1, *R. sativus* NAULB and *H. incana* NIJ were used. The IAGS pipeline (accessed Feb 2024) (Gao et al., 2022) was run based on the orthologous results from OrthoFinder and the non-overlapping syntenic blocks detected by Drimm-Synteny (accessed Feb 2024) (Pham and Pevzner, 2010).

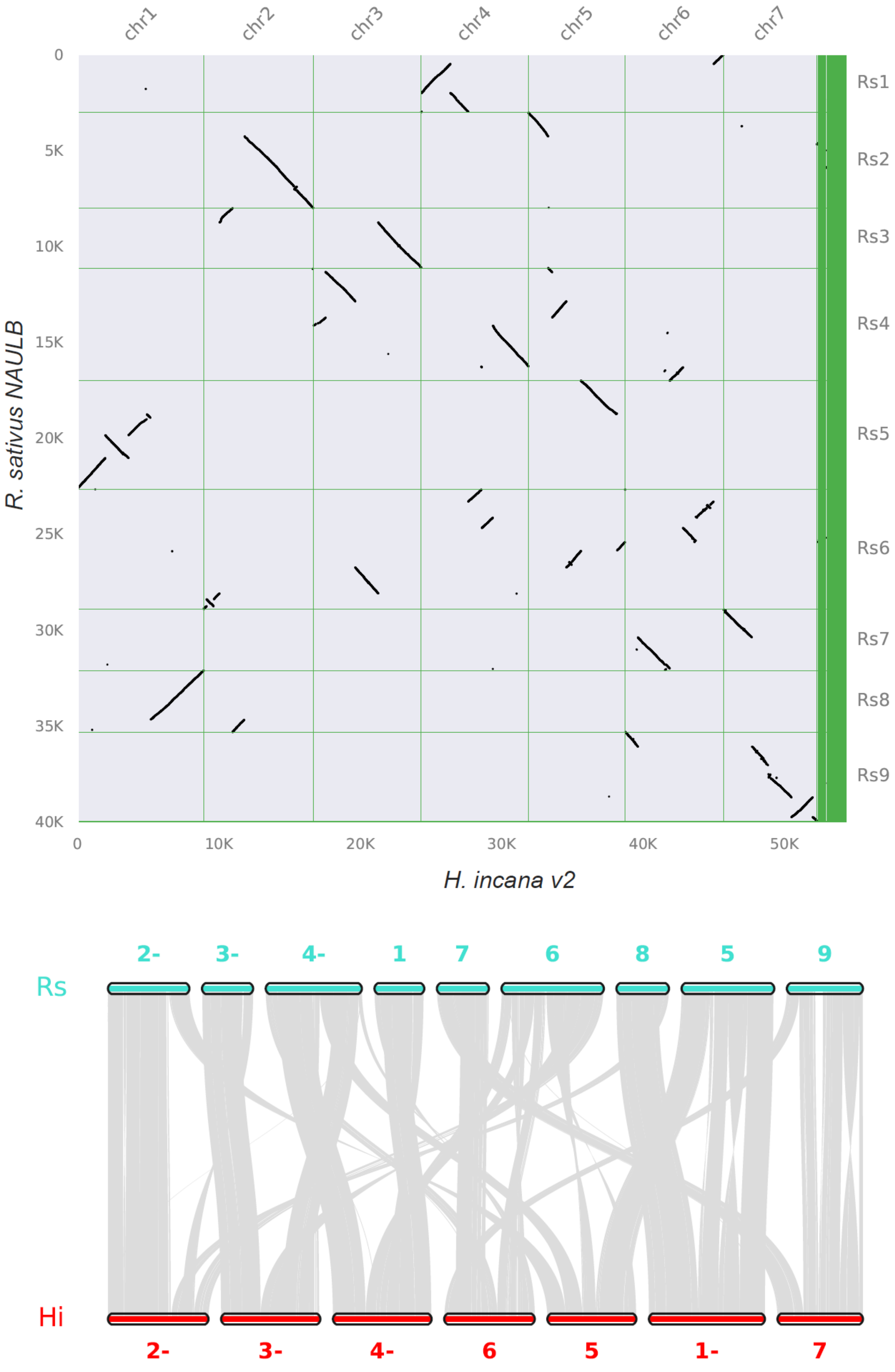


**Figure S15. Genome syntenic dotplot and ribbons between *R. sativus* and *H. incana* v2.0.** The genomes of *R. sativus* NAULB and *H. incana* NIJ were used. "-" denotes inverted sequence.

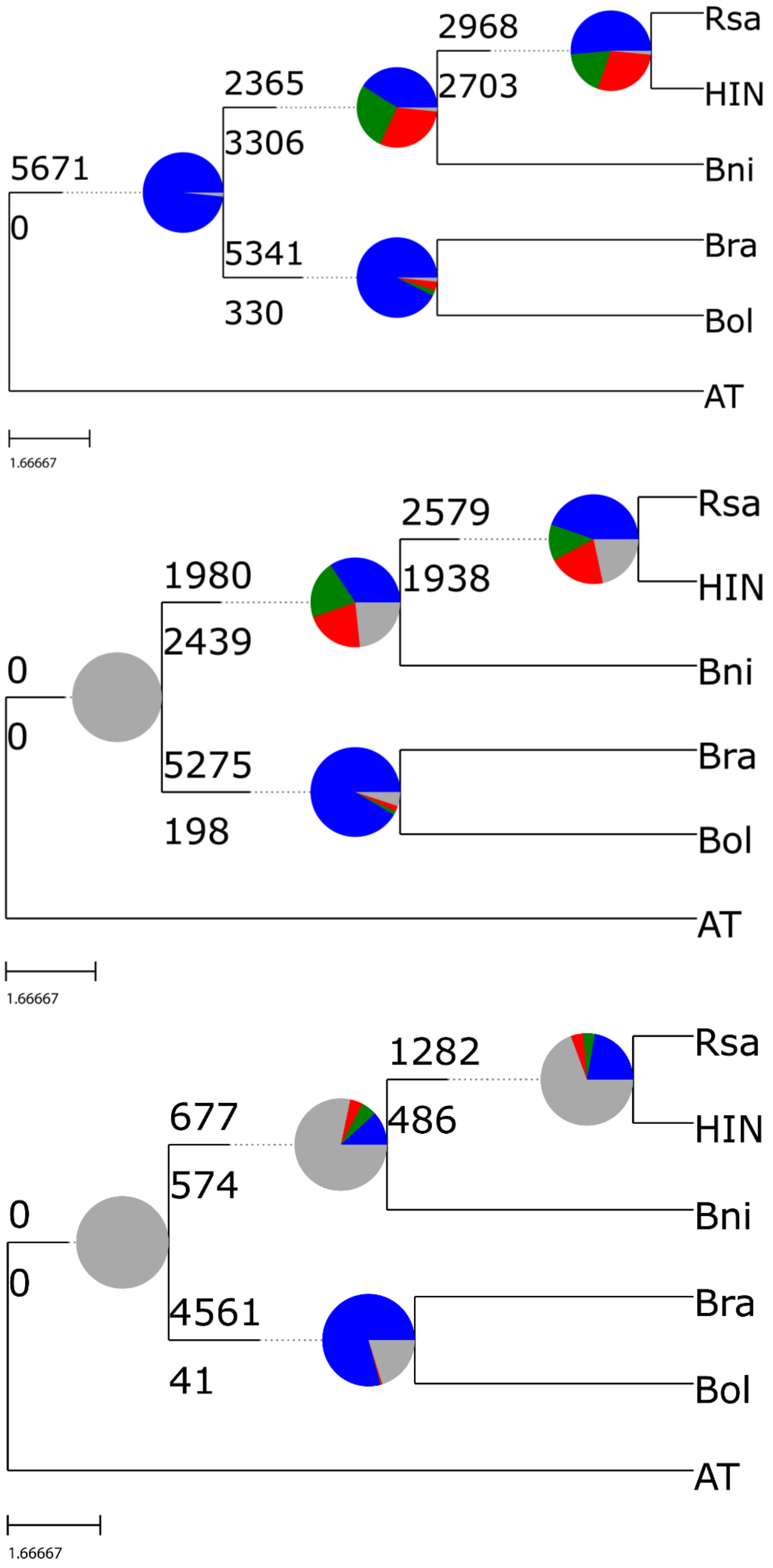


**Figure S16. PhyParts tree topology quantification** using all gene trees (top panel), gene trees with a bootstrap ≥50% (middle panel) and gene trees with a bootstrap ≥85% (bottom panel). The genomes of *A. thaliana* Col-0 (AT), *B. rapa* Chiifu (Bra), *B. oleracea* JZS (Bol), *B. nigra* NI100 (Bni), *R. sativus* NAULB (Rsa) and *H. incana* NIJ (HIN) were used.

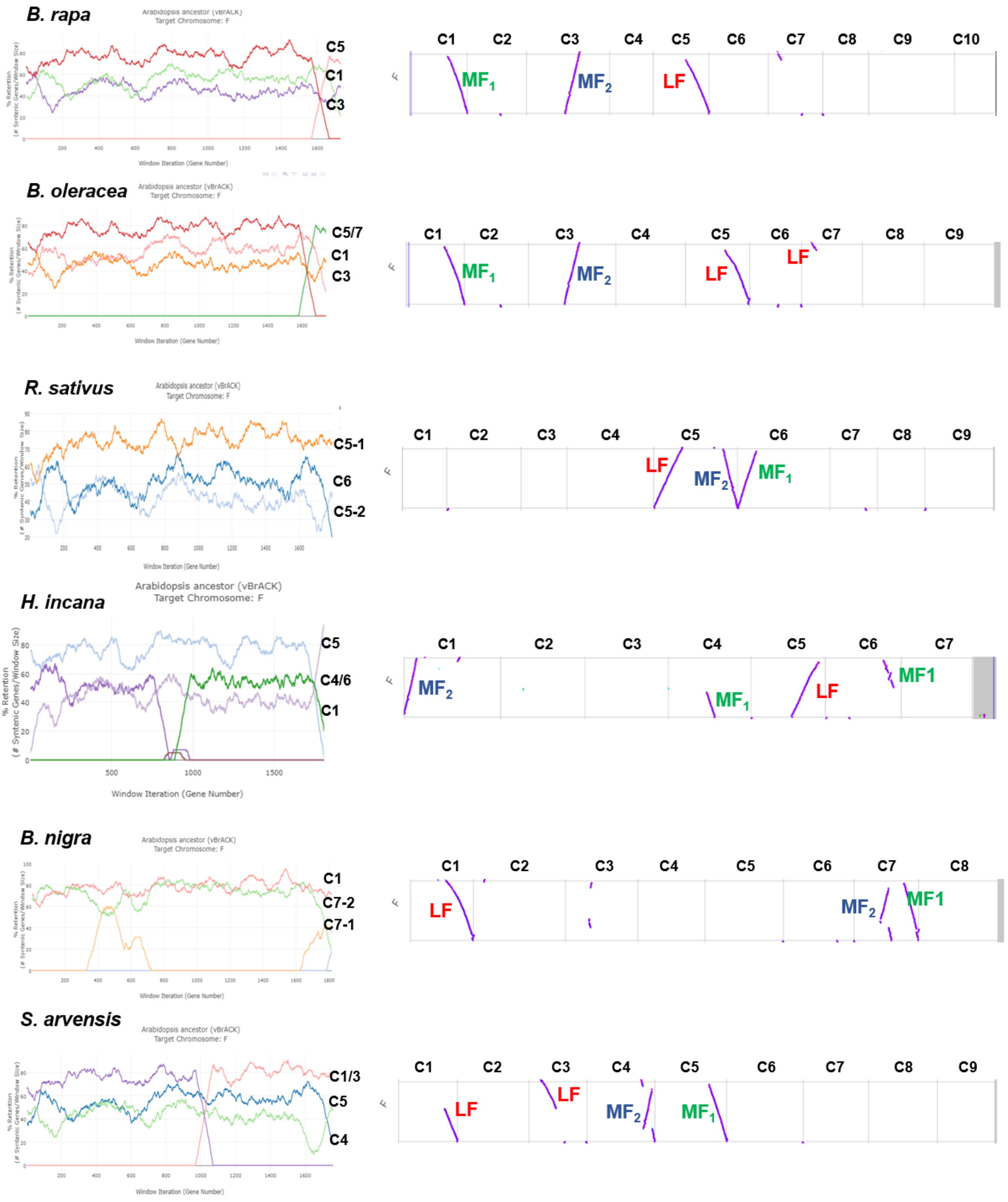


**Figure S17. The identification of the tPCK ancestral genomic block F in the six Brassiceae genomes. Left panel:** three triplicated ancestral genomic block F showing different fractionation rate corresponding to three sub-genome origins, LF, MF$_1$ and MF$_2$. **Right panel:** locations of the three triplicated ancestral genomic block F on each Brassiceae genome. The genomes of *B. rapa* Chiifu, *B. oleracea* JZS, *B. nigra* NI100, *S. arvensis* XJ1, *R. sativus* NAULB and *H. incana* NIJ were used. The plots were generated by SynMap ran together with FractBias (Joyce et al., 2017) on the CoGe v7. Note that for *B. nigra* genome, the fragment C7-1 (MF$_2$) was not fully shown in the left panel.

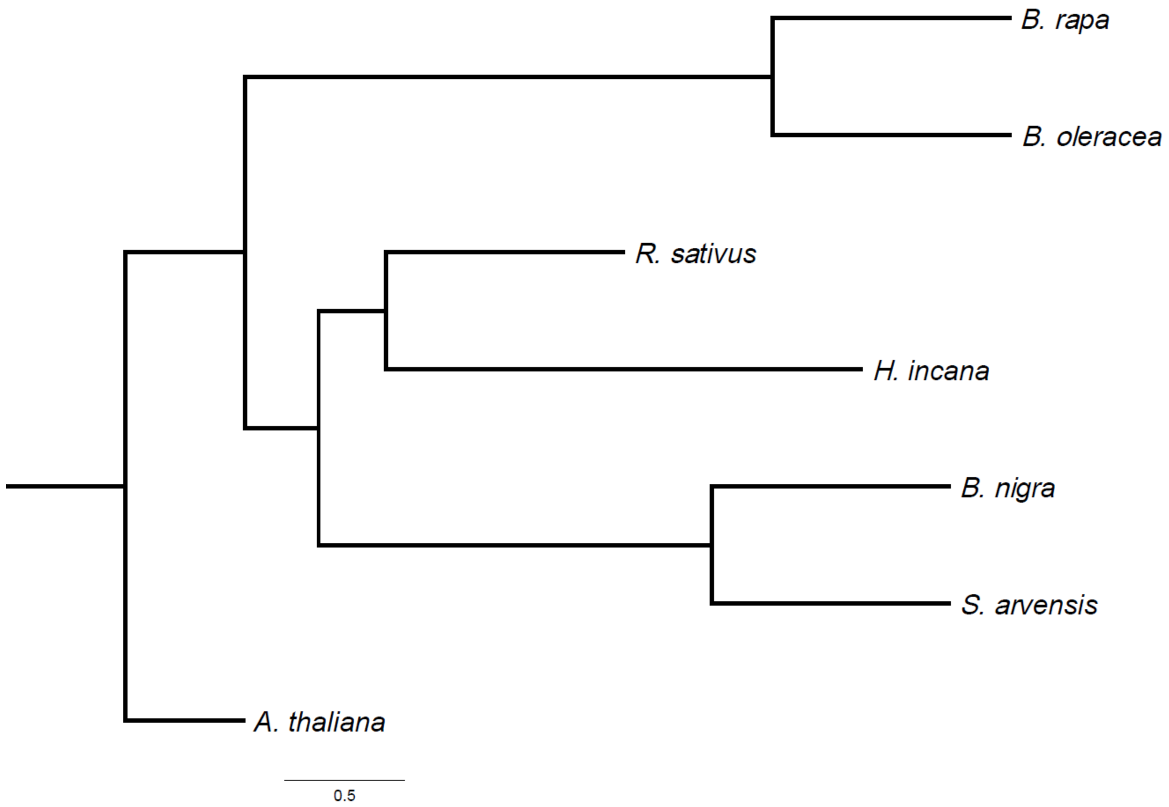


**Figure S18. Species tree reconstructed from 90 triad gene sets from three sub-genomes of each Brassiceae species** (as shown in **Figure S17**). All posterior probability supporting values are 1 (100%) and not shown. Branch length represents coalescence units.

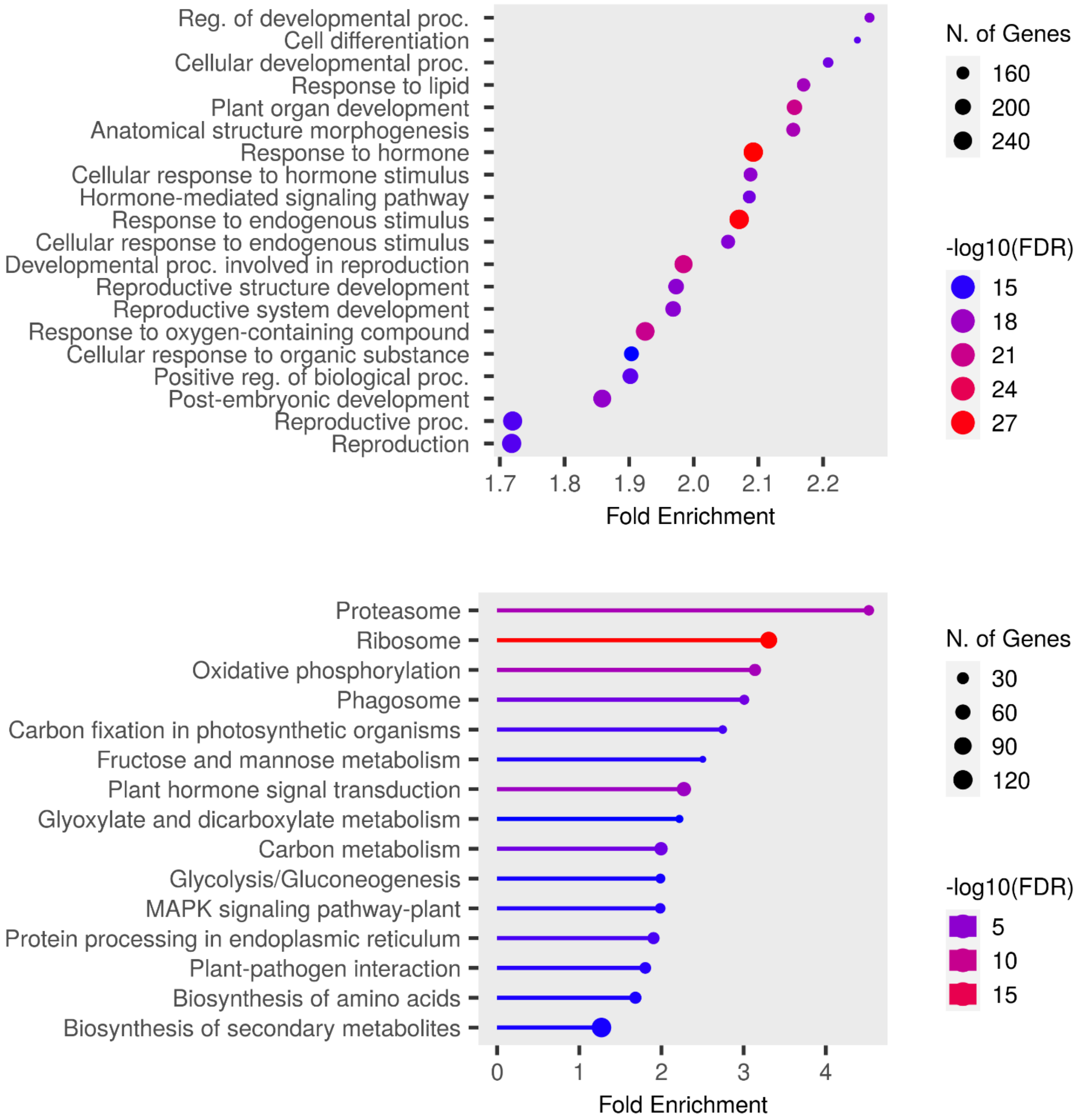


**Figure S19. GO biological process (upper panel) and KEGG pathway (lower panel) enrichment analyses of all well-retained 2,103 triad genes** highlighting topmost significant GO terms (FDR-corrected $p \leq 0.05$).

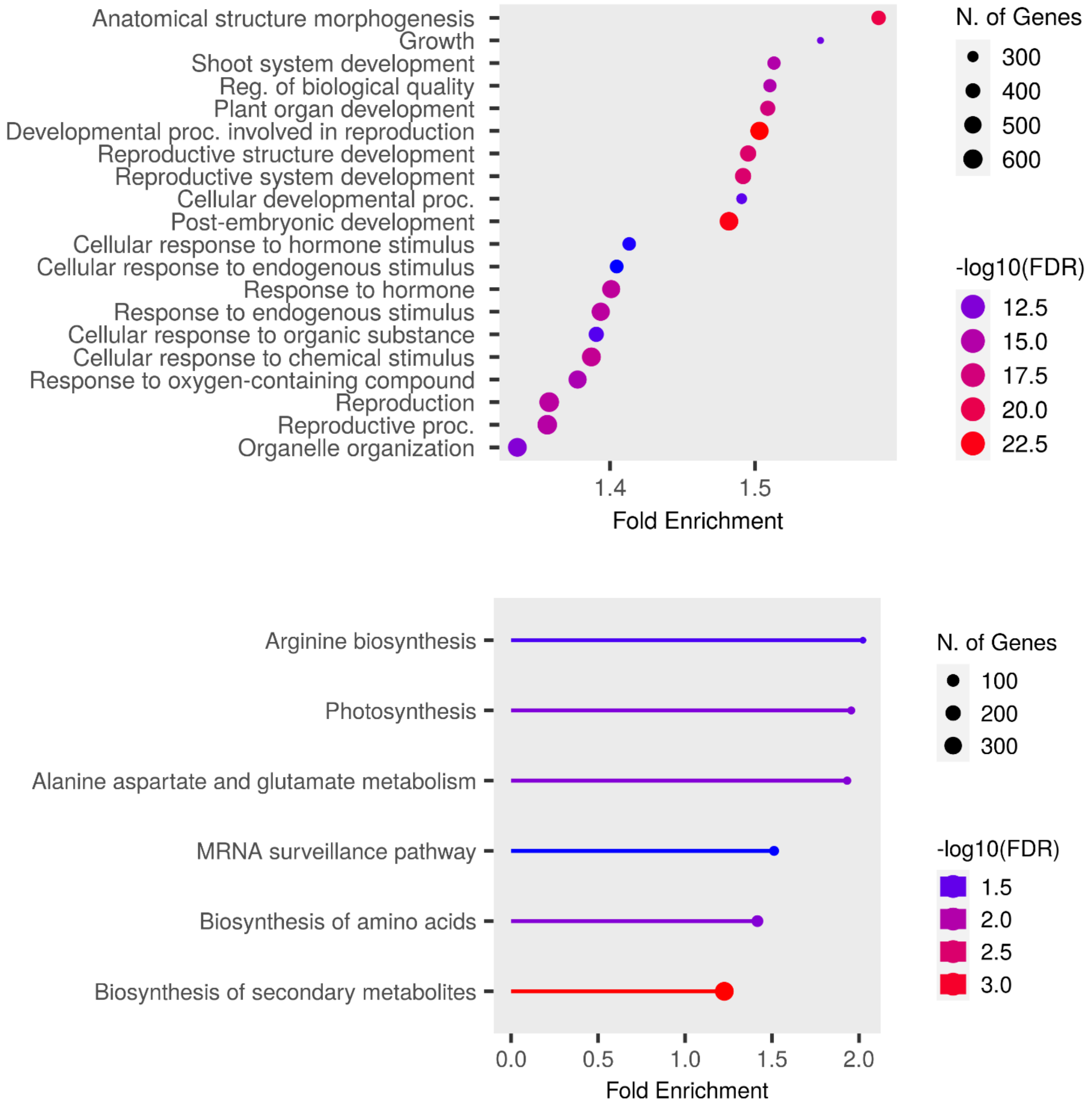


**Figure S20. GO biological process (upper panel) and KEGG pathway (lower panel) enrichment analyses of two-copy retained 6,457 dyad genes** highlighting topmost significant GO terms (FDR-corrected $p \leq 0.05$).

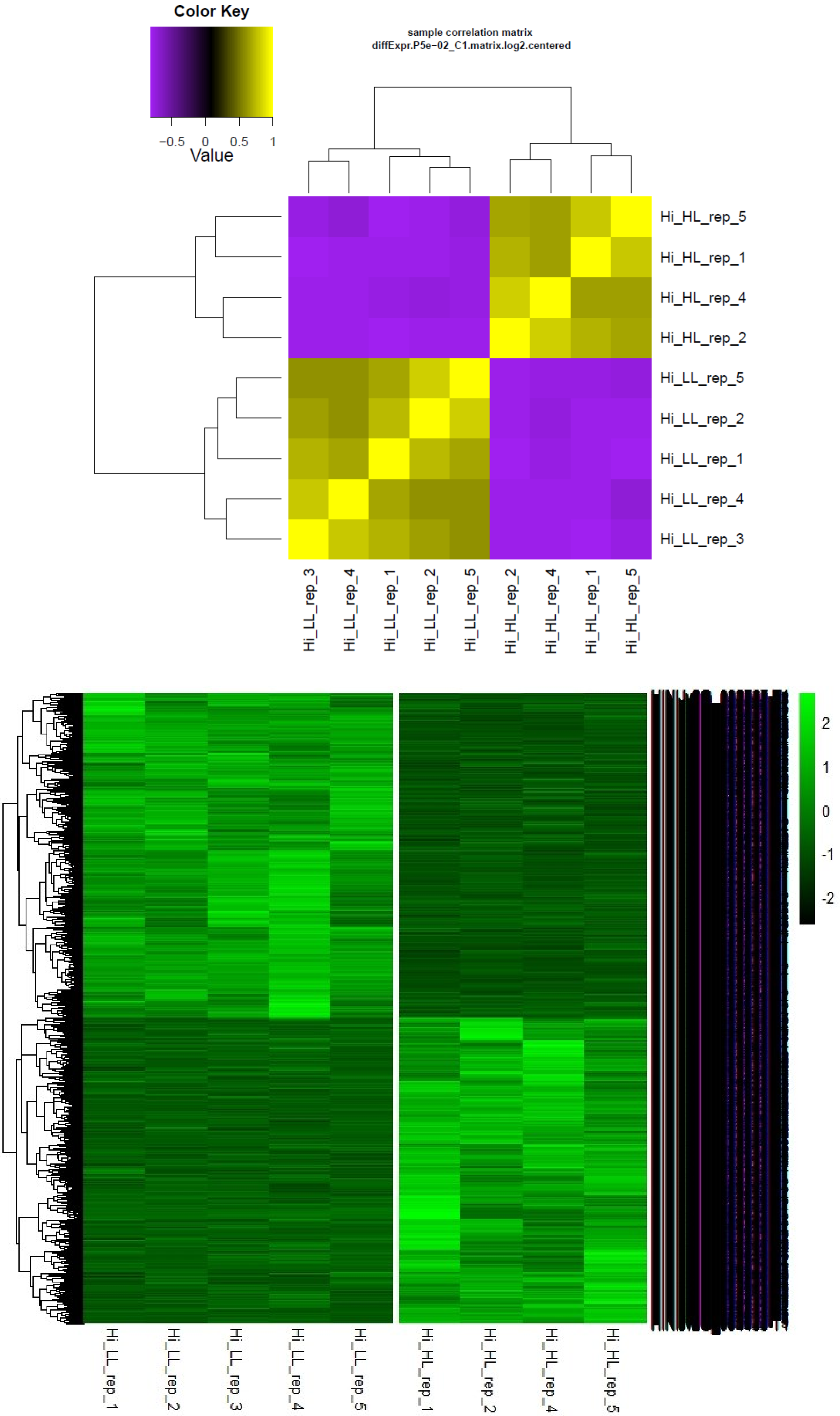


**Figure S21. A summary of *H. incana* whole canopy transcriptome data.** The *H. incana* RNA-seq sample correlation (upper panel) and the expression of the identified DEGs in two contrasting low (LL) and high-light (HL) conditions (lower panel). Gene expression was row-normalized. HI: *Hirschfeldia incana*.

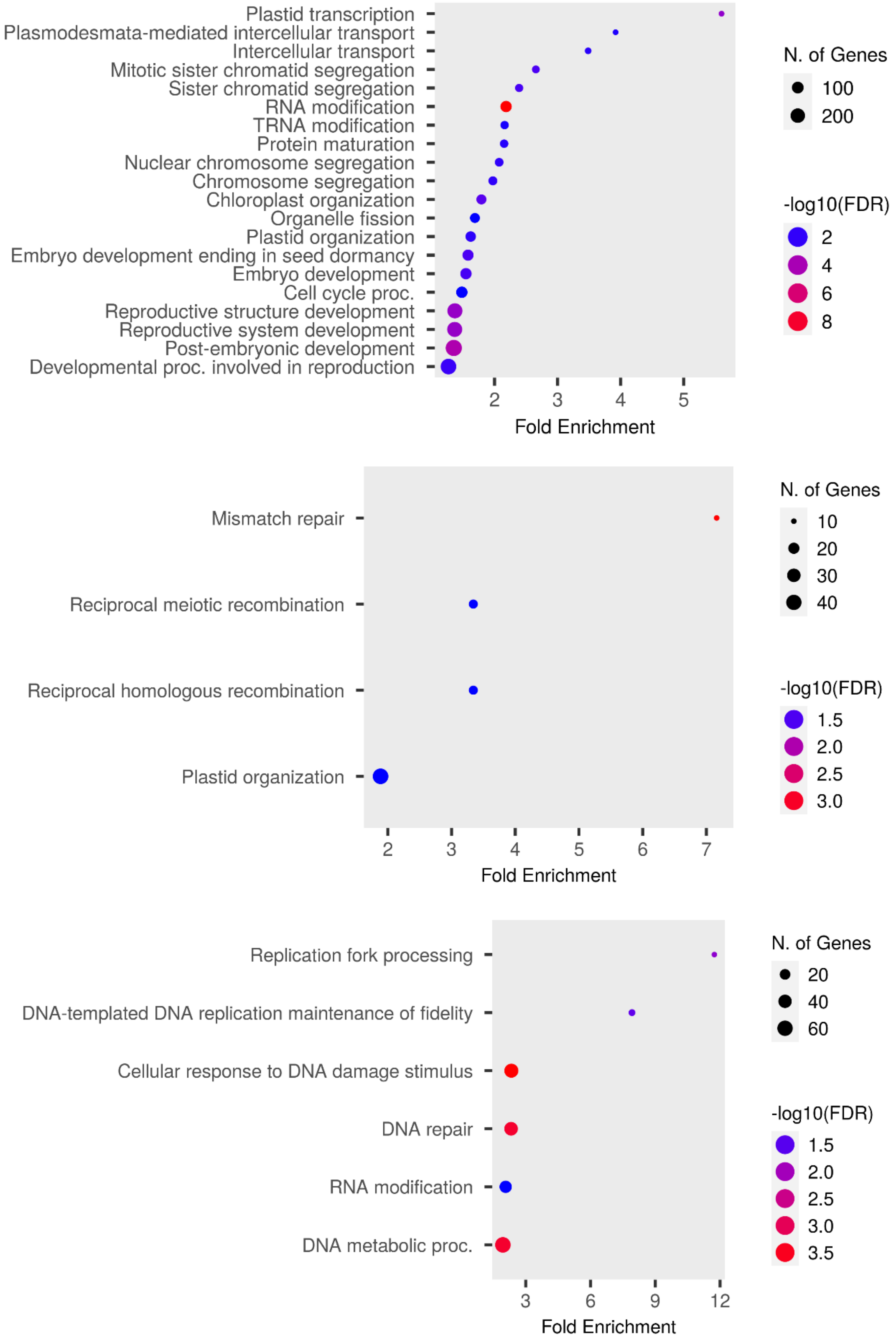


**Figure S22. GO biological process enrichment analyses of 3,527, 1,932 and 1,310 single-copy genes found on the LF, $MF_1$ and $MF_2$ sub-genomes**, respectively, highlighting significant GO terms (FDR-corrected $p \leq 0.05$).

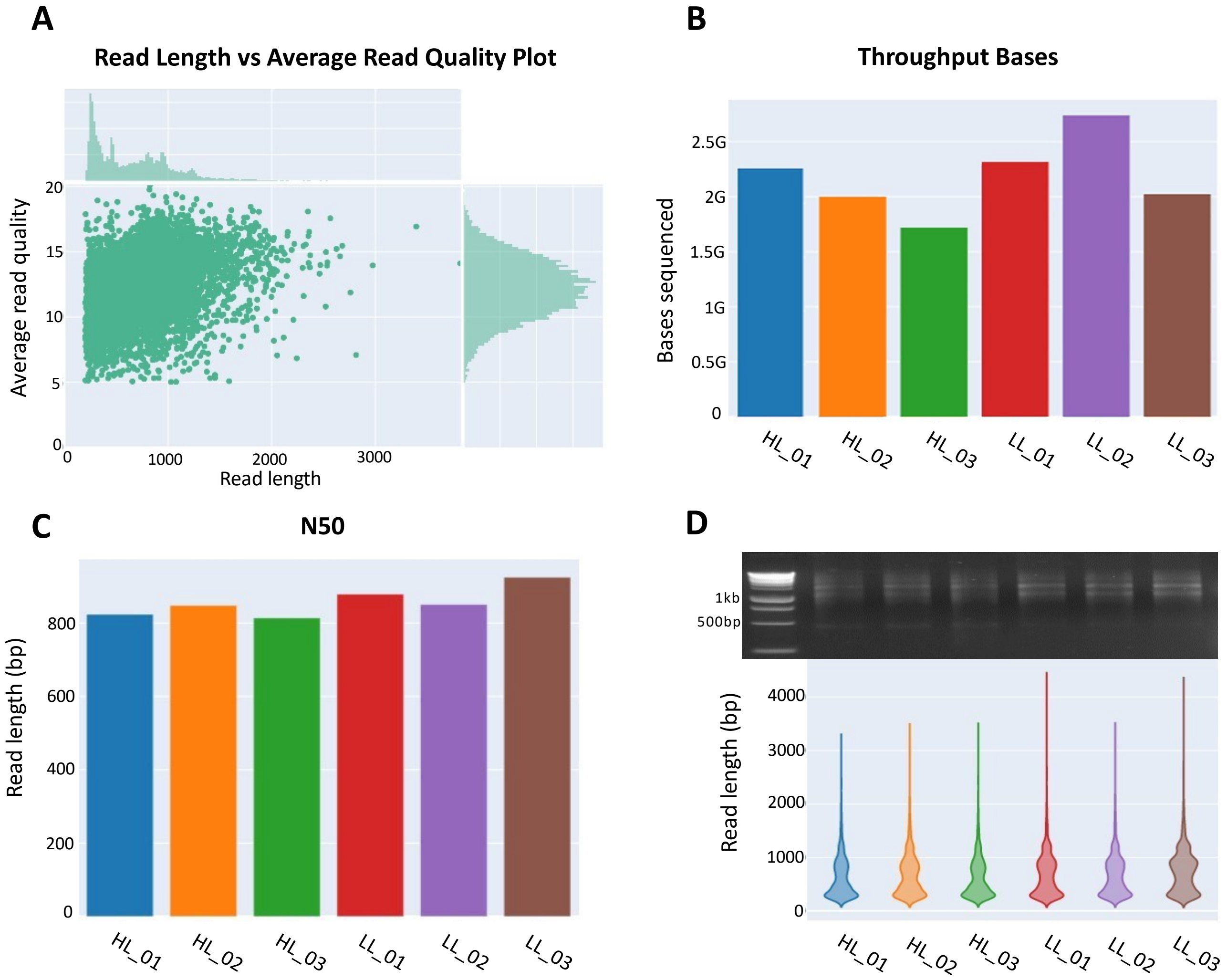


**Figure S23. Nanopore RNA sequencing data statistics.** (**A**) general read length and quality distribution dot plot. (**B**) comparison of sequenced sample size in bases bar-chart. (**C**) comparison of N50 values across samples bar-chart. (**D**) Comparison of sequencing libraries and read length distribution across samples. A 1kb DNA ladder was used. HL and LL denote high-light and low-light, respectively.
